# Conserved Cell Type Signatures Across the Brainstem and Spinal Cord in the Mouse Central Nervous System

**DOI:** 10.64898/2026.06.09.728039

**Authors:** Yuan Gao, Evgenii Kegeles, Jingyi Xie, Changkyu Lee, Yunfan Kong, Sam A. McClelland, Matthew T. Schmitz, Nelson J. Johansen, Judith Baka, Tamara Casper, Michael Clark, Katherine A. Fancher, Jessica Gloe, Jeff Goldy, Junitta Guzman, Carliana Halterman, Windy Ho, Marcus Hooper, Kelly Jin, Matthew Jungert, Rachel McCue, Nick Pena, Elliot Phillips, Augustin Ruiz, Nadiya V. Shapovalova, Daniel Sokolovsky, Emma D. Thomas, Amy Torkelson, Ruyi Yang, Shuguang Yu, Nick Dee, Kimberly A. Smith, Trygve E. Bakken, Bosiljka Tasic, Zhigang He, Hongkui Zeng, Zizhen Yao, Cindy T. J. van Velthoven

**Affiliations:** Allen Institute, Brain Science, Seattle, WA, USA; F.M. Kirby Neurobiology Center, Boston Children’s Hospital, Boston, MA, USA; Department of Neurology and Ophthalmology, Harvard Medical School, Boston, MA, USA; PhD Program in Biological and Biomedical Sciences, Harvard Medical School, Boston, MA, USA; Allen Institute, Brain Health, Seattle, WA, USA; Allen Institute, Neural Dynamics, WA, USA

**Keywords:** Single-cell transcriptomics, Spinal cord, Brainstem, Spatial transcriptomics

## Abstract

Understanding how cell types are organized across the central nervous system (CNS) is key to uncovering neural function. Here, we integrate single-nucleus Multiome (RNA+ATAC) sequencing, spatial transcriptomics, and computational analyses to map conserved cell type signatures in the adult mouse brainstem and spinal cord. We identify a shared core of neuronal and non-neuronal cell types, alongside region-specific specializations reflecting distinct functions. Spatial data reveal conserved cellular niches across the brainstem–spinal cord boundary, indicating a continuous organizational logic. Cross-region comparisons uncover recurrent gene expression modules and signaling programs that may support shared circuit features. Chromatin accessibility profiling highlights cell-type-specific regulatory programs and implicates *Hox* transcription factors in positional identity. Notably, cell-type and positional identities are largely orthogonal, with varying regional influence across neuronal classes: motor neurons show strong positional coupling, whereas glutamatergic and GABAergic interneurons show minimal entrainment. This work provides a reference for the shared molecular architecture of these CNS regions.

## INTRODUCTION

The spinal cord and brainstem together form the core of the vertebrate sensorimotor system, orchestrating fundamental processes including the integration of sensory signals, the execution of voluntary movement, and the maintenance of vital autonomic functions such as breathing, heart rate, and blood pressure^1–3^. Understanding how these regions perform such complex tasks requires a comprehensive knowledge of their constituent cell populations and their organization. These regions are anatomically continuous, yet they have historically been studied in isolation, within distinct research communities, model systems, and classification frameworks developing largely independently. Understanding how this extended neural axis is organized at the cellular level, including how its component cell types give rise to such diverse and precisely coordinated functions, remains a central challenge in neuroscience.

Historically, cell types in the spinal cord have been classified using various parameters, including location, morphology, embryonic lineage, electrophysiological properties, and connectivity^4,5^. While these approaches have provided valuable insights, the overarching principles that bridge molecularly defined subtypes with their connectivity, physiology, and function have remained unclear.

The advent of high-throughput single-cell and single-nucleus RNA sequencing (sc/snRNA-seq) has transformed this landscape by enabling unbiased, comprehensive characterization of gene expression across hundreds of thousands of individual cells simultaneously^6,7^. Applied to the spinal cord, these methods have produced increasingly complete molecular atlases that consolidate previously fragmented classification schemes into unified taxonomies grounded in transcriptional identity^8–11^. Crucially, these atlases have demonstrated that transcriptionally defined cell types correspond closely to functionally and developmentally characterized populations, validating the molecular approach as a principled framework for understanding neural diversity. On the other hand, brain-centric single-cell profiling has been applied to the brainstem, revealing that regions such as the ventrolateral medulla, a key regulator of cardiovascular and respiratory function and a major source of descending input to the spinal cord, contain a large diversity of molecularly distinct neuronal and non-neuronal subtypes^12–14^. Targeted profiling of specific brainstem populations, including spinally-projecting neurons and aminergic cell groups, has begun to link molecular identity to circuit function^15^.

Despite this progress, the organizational principles that structure neural diversity across this extended axis remain unclear. Thus, we set out to ask the following questions. Are the cell types of the brainstem and spinal cord organized according to the same rules? To what extent is cell type identity determined by intrinsic transcriptional programs versus positional signals that vary along the neuraxis? And how does molecular identity relate to the functional specialization of circuits at different levels of the axis?

Here we address these questions by generating and integrating a comprehensive single-nucleus RNA sequencing atlas spanning the full extent of the mouse brainstem and spinal cord. By profiling 109,373 nuclei from spinal cord and integrating this dataset with existing spinal cord and brainstem resources, we uncover homologous sets of transcriptomic cell types between brainstem and spinal cord and establish a unified molecular taxonomy comprising neuronal and non-neuronal cell types organized into a hierarchy of classes, subclasses, and supertypes. We leverage existing and new spatial transcriptomic data to map this taxonomy onto the tissue, revealing that the organizational logic of the spinal cord, including the conserved dorsal-ventral arrangement of inhibitory and excitatory populations and the niche structure of local cellular environments, extends continuously into the brainstem without a sharp boundary. Using chromatin accessibility profiling from paired gene expression and ATAC sequencing data, we characterize the cell-type-specific regulatory programs that underlie this taxonomy and identify transcription factors and cis-regulatory elements that define each cell type’s identity.

Finally, we show that neuronal identity across this axis is organized along two partially orthogonal axes: a positional axis defined by combinatorial HOX gene expression that varies by segment, and a cell-type axis defined by neurotransmitter identity, ion channel complement, and transcription factor programs that are broadly conserved across segments. The degree of coupling between these axes varies systematically by cell class. Motor neurons maintain essentially identical molecular programs regardless of segment, while glutamatergic interneuron diversity is strongly positionally entrained, with implications for understanding how the same fundamental circuit motifs can subserve different functions at different levels of the neuraxis. Together, these results provide a molecular foundation for understanding the organization and function of one of the most evolutionarily conserved and functionally critical regions of the vertebrate central nervous system.

## RESULTS

### Creation of a mouse spinal cord cell-type atlas

To systematically characterize the cellular diversity of the mouse brainstem and spinal cord, we constructed a multimodal single-cell atlas integrating transcriptomic, chromatin accessibility, and spatial transcriptomic data (**Figure 1a**). We generated snMultiome data from mouse spinal cord covering the cervical, thoracic, lumbar, and sacral regions. After stringent QC (see **Methods**), we compiled a dataset of 379,889 cells/nuclei, comprising 109,373 newly generated spinal cord nuclei and 270,516 cells/nuclei from published sources^8,11–13,15–21^ (**Figure S1**, **Supplementary Tables 1–2**). These datasets were integrated using scVI to correct for batch effects, and hierarchical clustering was performed on the integrated latent space (**Figure S2**). Multimodal integration of gene expression and chromatin accessibility data was further performed using MultiVI, enabling the inference of cell-type- and region-specific gene regulatory networks (GRNs) (**Figure 1a**).

**Figure 1.**
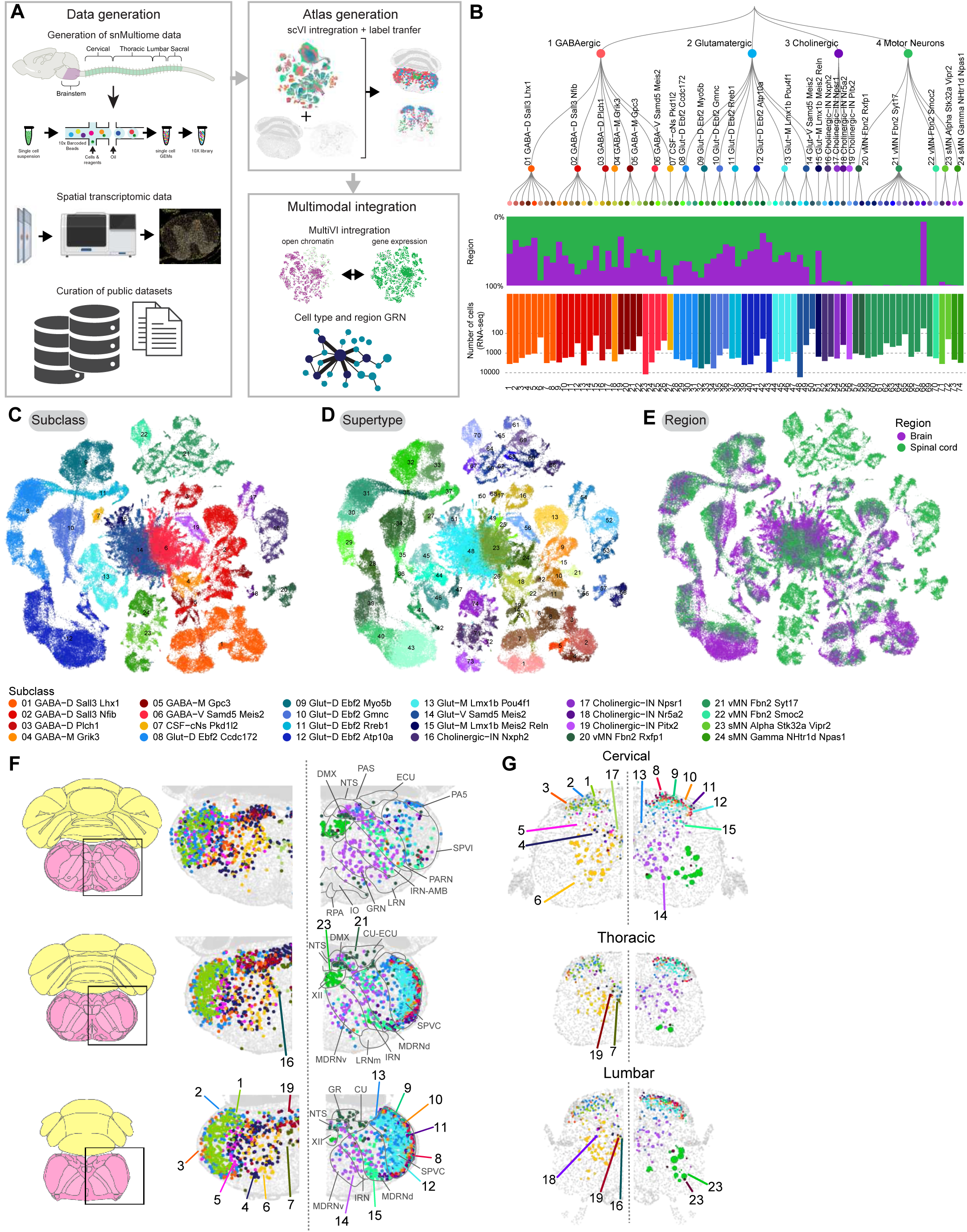
Construction of a multimodal single-cell atlas of the mouse hindbrain and spinal cord. **(A)** Schematic overview of the study workflow. Single-nucleus multiome data generation from cervical, thoracic, lumbar, and sacral regions of the spinal cord (top left). Spatial transcriptomic data acquisition and processing (middle left). Published dataset curation and integration (bottom left). Spatial data integration (top right). Multimodal integration of chromatin accessibility and gene expression data for GRN inference using MultiVI, (bottom right). **(B)** Hierarchical taxonomy of CNS cell types. Dendrogram illustrating the transcriptomic relationships among identified cell subclasses, grouped into four major classes: GABAergic, Glutamatergic, Cholinergic, and Motor Neurons. Bar plots below show the proportional contribution of each region (middle) and the absolute number of nuclei (bottom) per subclass. **(C-E)** UMAP embeddings of all profiled cells colored by (C) subclass identity (24 subclasses indicated in legend), (D) supertype identity illustrating fine-grained transcriptomic diversity within major cell classes, and (E colored by region of origin demonstrating the regional composition of the integrated atlas. **(F,G)** Spatial distribution of cell subclasses mapped onto coronal tissue sections, showing anatomically resolved localization of neuronal populations across brainstem (F) and spinal cord segments (G). The panel on the left shows the GABAergic and Cholinergic IN subclass and the panel on the right shows the Glutamatergic and Motor Neuron subclasses at the same level.

We established a transcriptomic taxonomy of spinal cord and related brainstem regions with four nested levels of classification, encompassing 5 classes, 38 subclasses, 93 supertypes, and 387 clusters (**Figure 1b, S1a, Supplementary Table 3**). The multidimensional relationships among supertypes are depicted using a hierarchical tree, UMAP embeddings, and constellation diagrams (**Figures 1b-e, S1a-f,h**). Cell-type annotations were assigned by label transfer from the Allen Brain Cell-Whole Mouse Brain (ABC-WMB atlas) and, where available, from original dataset annotations (**Figures S3, S4**). Regional contribution and cell number varied substantially across subclasses, reflecting the known anatomical specialization of the hindbrain and spinal cord (**Figure 1b**). To map cell type distributions across the spinal cord, we generated Xenium in situ transcriptomic data from transverse sections spanning the full rostral-caudal extent of the mouse spinal cord using the Xenium Prime 5K mouse pan-tissue panel supplemented with custom genes, with cell segmentation performed using a custom CellPose-based model (**Supplementary Table 5**; see **Methods**). These Xenium data, together with MERFISH data spanning the brainstem, were integrated with the RNA-seq reference atlas using scVI, and cell-type labels were transferred within the integrated latent space to annotate the spatial location of each supertype (see Methods).

Neuronal cell types constitute a large proportion of the spinal cord cell-type atlas, including 4 classes, 24 subclasses, 74 supertypes, and 310 clusters (**Figure 1b**). To further investigate the neuronal diversity, we generated re-embedded UMAPs (in 2D and 3D) for neuronal types described above, to reveal fine-grained relationships between neuronal types within and between regions in conjunction with the spatial transcriptomics data (**Figure 1c-e**). This analysis reveals a strong correspondence between transcriptomic specificity and relatedness and spatial specificity and relatedness among neuronal subclasses. Notably, neurons from the hindbrain and spinal cord show high transcriptomic similarity and cluster closely together, indicating shared molecular identities across these regions (**Figure 1c-e**). Spatial transcriptomic data further mapped these subclasses onto coronal tissue sections, demonstrating anatomically coherent localization of neuronal populations across brainstem and spinal cord segments (**Figure 1f,g**).

The four neuronal classes, GABAergic, glutamatergic, cholinergic interneurons, and motor neurons, show distinct spatial distributions that reflect their functional roles across the brainstem-spinal cord axis (**Figure 1f,g; Supp. Fig. S6**). GABAergic neurons, comprising 6 subclasses and 27 supertypes, are distributed across the dorsal and intermediate spinal cord and their brainstem equivalents, where they serve primarily inhibitory roles in sensory gating, pain modulation, and motor coordination. Glutamatergic neurons, with 8 subclasses and 24 supertypes, span the full dorsoventral extent from superficial dorsal horn to the ventral zone, encompassing populations involved in nociceptive relay, mechanosensory processing, locomotor rhythm generation, and ascending projection pathways (**Figure S6**). Cholinergic neurons comprise 4 subclasses including the V0c partition cell interneurons, which modulate motor neuron excitability via C-bouton synapses, and are enriched in intermediate and ventral regions of both spinal cord and brainstem^18,22^. Motor neurons form a fourth class of 5 subclasses, including visceral motor neurons (preganglionic autonomic neurons of the intermediolateral column) and skeletal motor neurons (alpha and gamma subtypes innervating extrafusal and intrafusal muscle fibers respectively). These are spatially restricted to ventral lamina IX and the intermediolateral column in the spinal cord, with corresponding populations in medullary motor nuclei in the brainstem. Cross-dataset correspondence with previously published spinal cord atlases confirms that these molecularly defined classes align with established functional and developmental categories (**Supp. Fig. S6**), providing a validated molecular reference for the spatial analyses described below.

Non-neuronal cells constitute a substantial and diverse component of the spinal cord cell-type atlas and encompass multiple glial, vascular, and immune-related populations. These cell types play essential roles in neural development, homeostasis, metabolic support, and injury response, and display distinct molecular and spatial characteristics within the spinal cord. All non-neuronal cell types across the spinal cord are classified into 1 class, 14 subclasses and 77 clusters (**Figure S6a-d**). Astrocyte and oligodendrocyte lineage populations exhibited regionally biased distributions of white matter versus grey matter (**Figure S6e,g,h**). Ependymal cells, arachnoid barrier cells (ABC), and Vascular Leptomeningeal Cells (VLMC) were located near the central canal, and meninges as expected (**Figure S6f,i,j**). Vascular cells and microglia were broadly dispersed, consistent with their respective roles (**Figure S6k-n**).

### Spatial organization of neuronal cell types in the spinal cord and brainstem

Spatial transcriptomics analysis revealed a highly structured organization of neuronal cell types within the spinal cord, reflecting both regional patterning and functional specialization (**Figure 2**). Neuronal subclasses and supertypes exhibited distinct spatial distributions along the dorsoventral and mediolateral axes, consistent with known anatomical and developmental organization of the spinal cord.

**Figure 2.**
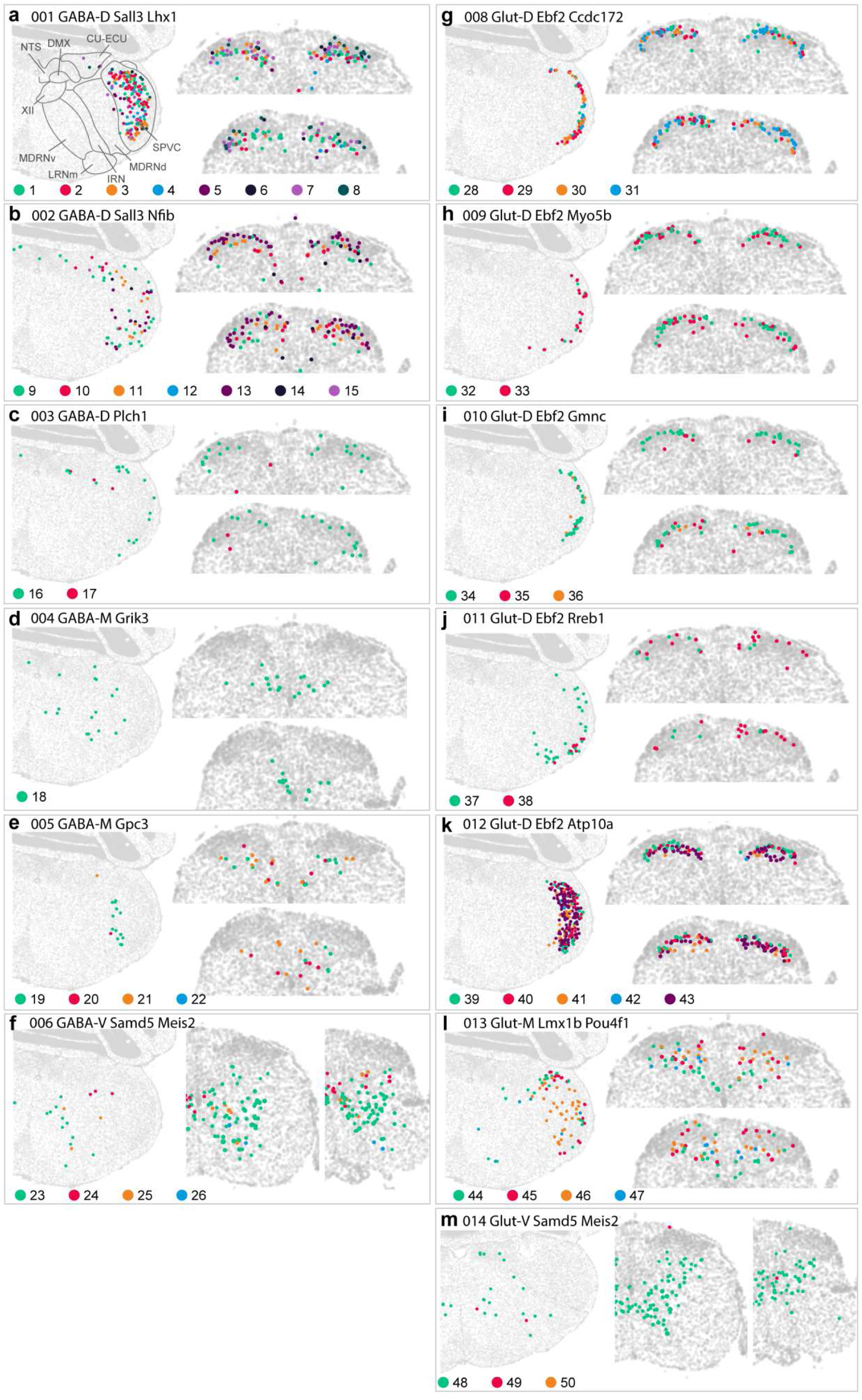
Spatial distribution of neuronal subclasses across CNS tissue sections. (A–M) Spatial transcriptomic maps showing the anatomical localization of individual neuronal subclasses projected onto coronal sections of the brain and spinal cord. Each panel displays cells belonging to a single transcriptomically defined subclass, with individual supertypes distinguished by color. Subclasses are grouped by neurotransmitter identity and spatial location: **GABAergic subclasses (A–F): (A)** 01 GABA-D Sall3 Lhx1 (supertypes 1–8), **(B)** 02 GABA-D Sall3 Nfib (supertypes 9–15), **(C)** 03 GABA-D Plch1 (supertypes 16–17), **(D)** 04 GABA-M Grik3 (supertype 18), **(E)** 05 GABA-M Gpc3 (supertypes 19–22), **(F)** 06 GABA-V Samd5 Meis2 (supertypes 23–26). **Glutamatergic subclasses (G–M): (G)** 08 Glut-D Ebf2 Ccdc172 (supertypes 28–31), **(h)** 09 Glut-D Ebf2 Myo5b (supertypes 32–33), **(i)** 10 Glut-D Ebf2 Gmnc (supertypes 34–36), **(j)** 11 Glut-D Ebf2 Rreb1 (supertypes 37–38), **(k)** 12 Glut-D Ebf2 Atp10a (supertypes 39–43), **(l)** 13 Glut-M Lmx1b Pou4f1 (supertypes 44–47), **(m)** 14 Glut-V Samd5 Meis2 (supertypes 48–50). Colored dots represent individual cells, with each color corresponding to a distinct supertype as indicated in the legend below each panel. Gray tissue outlines depict reference anatomy.

GABAergic subclasses occupied largely non-overlapping spatial territories organized along the dorsoventral axis, with systematic correspondence between spatial position and functional identity (**Figure 2a-f**). Dorsal GABAergic subclasses (01-03, GABA-D Sall3 and Plch1, **Figure 2a-c**) were concentrated in superficial laminae, with individual supertypes mapping to known functional populations: supertypes 002 (GABA-D Nrgn) and 003 (GABA-D Rxfp2 Syt10) occupied the most superficial positions consistent with their annotation as pain-gating interneurons in laminae I-II^9^, while supertype 008 (GABA-D Sstr2 Pnoc), which encodes nociceptin/orphanin and has been shown to gate mechanical allodynia, was similarly confined to superficial dorsal positions^23,24^ (**Figure S7**, **Supplemental Table 4**). Supertype 016 (GABA-D Npy Ecel1), which gates mechanical and cold pain^16,23,25,26^, co-occupied lamina I with these pain-gating populations, consistent with a convergence of multiple inhibitory mechanisms at the first nociceptive relay. Deeper dorsal GABAergic populations including 013 (GABA-D Prox1 Gal), which expresses dynorphin and has been implicated in gating mechanical allodynia through dynorphin-mediated inhibition^9,27^, and 014 (GABA-D Kcnq4), a candidate inhibitory neuron of the low-threshold mechanoreceptor (LTMR)-recipient zone in laminae III-IV, occupied positions deeper in the dorsal horn consistent with their roles in modulating ascending sensory transmission at the level of the low-threshold mechanoreceptor input zone rather than primary afferent terminals directly^9,28^.

At intermediate dorsoventral positions, GABA-M subclasses (04-05, **Figure 2d,e**) populated laminae IV-VI, a region classically associated with sensorimotor integration^29^. Supertypes within these subclasses including 018 (GABA-M Klhl14), 020 (GABA-M Nxph2 Gpc3), and 021 (GABA-M Rab38) are annotated as premotor inhibitory interneurons (**Figure S7**, **Supplemental Table 4**), consistent with their intermediate laminar position between the dorsal sensory-processing zone and the ventral motor output zone. This spatial positioning is consistent with their proposed role in integrating descending motor commands with ascending sensory signals to coordinate motor output^30^. The most ventral GABAergic populations (subclass 06, GABA-V Samd5 Meis2, **Figure 2f**) occupied laminae VII-VIII, encompassing supertypes with V1 and V2b lineage annotations including Renshaw cells (026, GABA-V Ccbe1 Synpr)^9,31^, flexor-extensor coordination interneurons (023-024)^5,9^, and CSF-contacting neurons at the central canal (027, CSF-cNs Pkd1l2), which monitor the cerebrospinal fluid, sense spinal cord bending, and modulate locomotor rhythm^9,32^.

Glutamatergic subclasses showed an analogous dorsoventral stratification with distinct functional specializations at each level (**Figure 2g-m**). Dorsal *Ebf2*-expressing subclasses (08-12, **Figure 2g-k**) were consistently positioned in superficial to deep dorsal laminae I-IV, with supertypes mapping to multiple functional categories including nociceptive relay, itch processing, and mechanosensory circuits^8,23,33,34^ (**Figure S7**, **Supplemental Table 4**). Supertype 031 (Glut-D Nmur2 Car12), occupying superficial laminae I-II, corresponds to the GRP-related itch relay which is consistent with its Nmur2 expression and correspondence to the Russ et al. Excit-10 cluster associated with chemical itch sensation^9,28,35^. Supertypes 032 (Glut-D Tac1 Otof Lmo3), whose Tac1 expression marks substance P-expressing neurons driving sustained nociceptive behaviors, and 040 (Glut-D Nts Tac2 Calca), expressing CGRP (encoded by *Calca*) marking spinoparabrachial projection neurons also localized to superficial laminae I-II, the latter is consistent with its role as a nociceptive relay with ascending projection identity^36,37^. Excitatory populations including 039 (Glut-D Car8), which occupies the LTMR-recipient zone of laminae II-IV and processes static touch input under normal conditions^9,23,38^, and 043 (Glut-D Maf Mafa), which is activated by A-fiber light touch input in laminae III-IV^38,39^, both correspond to populations implicated in mechanical allodynia following inflammatory and neuropathic injury^33^, suggesting they serve as a circuit node linking normal mechanosensory processing to pathological pain states.

Intermediate glutamatergic populations (subclass 13, Glut-M Lmx1b Pou4f1, **Figure 2l**) occupied laminae IV-VI, a region associated with both sensorimotor integration and the origin of ascending projection pathways^5,40^. Within this subclass, supertype 045 (Glut-M Prox1 Pdyn), which co-expresses Tacr1 and dynorphin, corresponds to the Excit-25 cluster of Russ et al.^9^ and is annotated as a candidate ascending somatosensory relay neuron^9^. Supertype 046 (Glut-M Cck), expressing cholecystokinin, occupied equivalent intermediate laminae and represents a candidate pain amplification population, consistent with the known role of CCK-expressing neurons in facilitating nociception^33,41^. Supertype 044 (Glut-M Cep112 Sntg2) occupied the sensorimotor integration zone of laminae IV-VI, consistent with its annotation as a candidate proprioceptive relay interneuron positioned to integrate sensory and descending motor signals at the spinal cord intermediate zone^9,42^.

The most ventral glutamatergic populations (subclass 14, Glut-V Samd5 Meis2, **Figure 2m**) occupied laminae VII-VIII, corresponding to the ventral interneuron zone associated with locomotor circuit organization. Supertype 048 (Glut-V Htr4 Cdh9) corresponds to a V2a or V3 lineage candidate and represents excitatory interneurons of the locomotor Central Pattern Generator (CPG) involved in rhythm generation^9^, while supertype 049 (Glut-V Onecut2), an early-born ventral excitatory population with high Onecut2 expression, is annotated as a candidate long-range projection neuron in the ventral spinal cord^39,43^. Together the intermediate and ventral glutamatergic populations span a functional continuum from proprioceptive and nociceptive relay through ascending projection to locomotor CPG, mirroring the layered functional organization seen in the GABAergic compartment and reflecting the division of excitatory interneuron function along the dorsoventral axis.

Across all subclasses, the correspondence between spatial position and functional annotation was striking: supertypes assigned to pain gating consistently occupied superficial dorsal positions, touch relay and mechanosensory populations localized to the LTMR-recipient zone, premotor interneurons clustered in intermediate laminae, and CPG excitatory interneurons concentrated in the ventral horn.

Together, these results showed that neuronal supertypes in the spinal cord occupy highly stereotyped positions along the dorsoventral axis, raising the possibility that the local arrangement of molecularly defined cell types delineates spatial domains resembling the classical Rexed laminae, which were originally defined by cytoarchitecture and are associated with distinct sensory, motor, and autonomic functions. To examine this, we performed niche analysis, in which each cell is characterized not only by its own molecular identity but also by the composition of its immediate cellular neighborhood **(Figure 3A)** ^44^. By incorporating local cellular context, this approach extends the information provided by individual supertype distributions and tests whether laminar organization is also reflected in recurrent local combinations of neuronal types.

**Figure 3.**
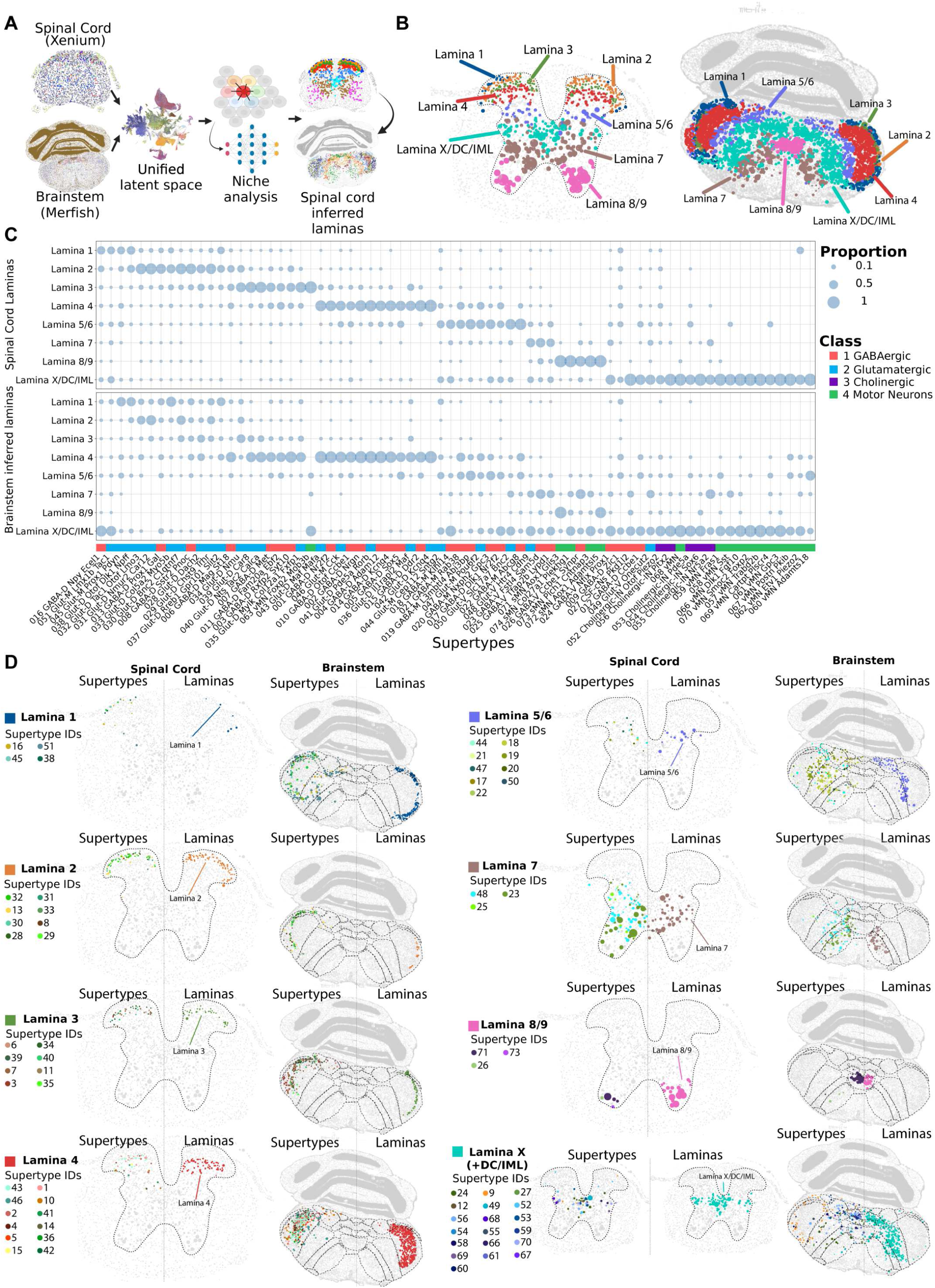
Shared spatial organization of cell types across brainstem and spinal cord. **(A)** Schematic of the analysis logic. **(B)** Representative section of the laminae of the spinal cord and inferred laminae organization of the brainstem. **(C)** Bubble plot showing distribution of each supertype across laminae in the spinal cord and inferred laminae in brainstem. Values are normalized per supertype. **(D)** Distribution of each lamina in the spinal cord and brainstem side by side with the defining supertypes for each lamina.

Niche analysis identified 55 distinct spatial niches in the spinal cord (**Figure S8A,B**). Clustering of these niches based on supertype composition revealed that they form distinct domains that appeared to be spatially reminiscent of the classical Rexed laminae (**Figure S8A**). To verify this and determine whether spinal laminar organization is encoded by the compositional arrangement of neuronal supertypes, we independently assigned each niche to a lamina using classical anatomical definitions together with known molecular markers^9,45^. The domains identified by unbiased clustering were largely consistent with the classical laminar organization described previously (**Figures 3B and S8C-F**), suggesting that the local compositional organization of cell types is sufficient to recapitulate the laminar structure of the spinal cord.

To relate these niche analysis-based spatial domains to established spinal cord anatomy, we annotated each niche-derived domain according to its correspondence with classical Rexed laminae. Domains corresponding to laminae I–IX showed a close match to canonical laminar definitions, spanning superficial nociceptive-processing regions of the dorsal horn, deeper low-threshold mechanoreceptive and sensorimotor integration zones, intermediate premotor and autonomic regions, and ventral motor-associated laminae. The lamina X-associated domain included lamina X proper around the central canal together with related central and accessory regions, including the intermediolateral column, lateral spinal nucleus, and dorsal commissural region of the sacral spinal cord. Together, these annotations established a molecular laminar framework that we use throughout the manuscript to relate neuronal subtype organization to anatomical position and functional specialization (**Figures 3C,D, S9A-D**).

Niche analysis identified 55 distinct spatial niches in the spinal cord **(Figure S8A, B)**. Clustering of these niches based on supertype composition revealed broader spatial domains that were reminiscent of the classical Rexed laminae **(Figure S8A-E)**. To relate these niche-derived domains to established spinal cord anatomy, we annotated each domain according to its correspondence with classical Rexed laminae using anatomical definitions together with known molecular markers. Domains corresponding to laminae I–IX closely matched canonical laminar definitions, spanning superficial nociceptive-processing regions of the dorsal horn, deeper low-threshold mechanoreceptive and sensorimotor integration zones, intermediate premotor and autonomic regions, and ventral motor-associated laminae. The lamina X-associated domain included lamina X proper around the central canal together with related central and accessory regions, including the intermediolateral column, lateral spinal nucleus, and dorsal commissural region of the sacral spinal cord **(Figure S8F)**. Consistent with this framework, most supertypes were enriched in distinct laminar domains **(Figures 3C).**

### Shared Organizational Principles between Spinal Cord and Brainstem

Strikingly, the same cell types were detected in spatially equivalent positions in brainstem sections shown alongside the spinal cord data (**Figure 2**), suggesting that a similar organizational logic may extend into the brainstem. Unlike the spinal cord, the brainstem has not traditionally been described as having a continuous laminar cytoarchitecture, and its organizational principles are less immediately apparent from classical histological examination alone. We therefore extended the niche analysis to combined spatial transcriptomic data from both structures to test whether conserved local cellular organization could be formally identified across the brainstem-spinal cord axis.

To address this, we leveraged our spatial dataset together with our scRNA-seq dataset to generate a shared latent space across spinal cord and brainstem samples, and then performed niche analysis on the combined data to derive a shared niche embedding across both structures **(Figure 3A)**. We next mapped individual niches onto the spinal cord lamina assignments described above and used these laminar domains as a common spatial framework for side-by-side comparison between the two structures. **(Figure 3B)** This analysis revealed that transcriptomically defined neuronal types largely co-occur in shared spatial domains across both regions **(Figure 3C,D; Figure S9, 10)**, supporting the idea that the spinal cord and caudal brainstem share a common organizational logic at the level of local cellular neighborhoods.

Examining individual laminar domains, spinal laminae I-II aligned with the superficial layers of the caudal trigeminal nucleus **(Figure 3D, Figure S10, Figure S11A,B),** consistent with the known role of both regions in processing nociceptive and thermal sensory information. Predicted ascending supertypes 045 Glut-M Prox1 Pdyn and 051 Glut-M Lmx1b Tac1, enriched in laminae I and V-VI of the spinal cord, and in the case of supertype 051 Glut-M Lmx1b Tac1, additionally in the lateral spinal nucleus, were also enriched in the superficial layer of the caudal trigeminal nucleus (SPVC). In addition, these cells were distributed along the boundary between the SPVC and the dorsal medullary reticular nucleus (MDRNd). This conserved spatial organization of ascending neuronal populations suggests that elements of the anterolateral system may follow a shared organizational logic across the spinal cord and caudal brainstem.

Along the rostro-caudal axis, however, this organization was not uniform. More rostrally, the superficial lamina I-like population progressively diminished within the trigeminal nucleus (SPVC), whereas the ventral ascending population persisted, with supertype 051 Glut-M Lmx1b Tac1 becoming increasingly prominent. Since laminae I and II are well-established recipients of nociceptive and somatosensory primary afferent input in the spinal cord we hypothesized that the diminished presence of laminae I/II-specific cell types may correlate with changes in primary afferent innervation. To test this prediction directly, we co-registered spatial transcriptomics-defined lamina assignments with immunohistochemical markers across sections spanning both the spinal cord and caudal brainstem. In both structures, spatial transcriptomics-assigned lamina I cells co-localized with CGRP immunoreactivity, marking peptidergic nociceptive afferent input zones, while lamina II cells co-localized with IB4 labeling (**Figure S11C**), marking non-peptidergic nociceptive afferent input zones. We note, however, that supertype 040 (Glut-D Nts Tac2 Calca), which is itself enriched in dorsal horn (however in deeper laminas) encodes *Calca/*CGRP, indicating that CGRP-immunoreactivity in the brainstem could reflects in part local neuronal expression in addition to primary afferent terminals^46^. Similarly, whether IB4 labels local medullary neurons in addition to afferent terminals in the brainstem remains to be determined. The co-localization patterns observed here are therefore consistent with, though not definitive proof of, primary afferent innervation of lamina I/II-equivalent brainstem regions. Interestingly, IB4 and CGRP labeling formed distinct layers in the spinal cord but showed greater overlap in more rostral brainstem sections, consistent with the higher degree of lamina I/II intermixing predicted by our analysis (**Figure S11C**).

Together, these observations suggest that the caudal brainstem contains both superficial nociceptive-like and deeper wide-dynamic-range-like ascending populations, similar to spinal cord, whereas more rostral regions are characterized predominantly by persistence of the ventral, wide-dynamic-range-like ascending component^47^.

Laminae III-IV, along with its representative supertypes, mapped to highly stereotyped deeper layers of the trigeminal nucleus in the brainstem, suggesting convergence in regions associated with tactile processing and LTMR-related sensory integration. Notably, the lamina IV-like domain appeared broader in the brainstem than in the spinal cord, raising the possibility that this territory is expanded in association with the increased demands of facial tactile processing, particularly in rodents, which rely extensively on the face and whiskers for environmental exploration. **(Figure 3D, Figure S10)**

The lamina V/VI-like domain, in turn, mapped near the border of deep spinal trigeminal nucleus, caudal and interpolar parts (SPVC and SPVI respectively) and MDRNd in caudal brainstem. This domain also extended into neighboring dorsal column nuclei, including the cuneate and external cuneate nuclei, consistent with the involvement of this territory in somatosensory integration across trigeminal and body-related sensory pathways. This region was enriched for a consistent set of supertypes shared with the spinal cord, including 018 GABA-M Klhl14, 019 GABA-M L3mbtl4 Abi3bp, 020 GABA-M Nxph2 Gpc3, and 022 GABA-M Abcg8, as well as for a wide-dynamic-range-like ascending population (051 Glut-M Lmx1b Tac1). This organization suggests that the corresponding brainstem territory may serve functions analogous to deep dorsal horn circuits, including convergence of sensory inputs, and relaying this information to the higher brain structures and integration with descending control systems. Laminae I-III-like identities were detected only in the SPVC region, whereas SPVI was represented predominantly by laminae IV-VI-like cell types, emphasizing a caudo-rostral shift toward mechanosensory integration in SPVI rather than nociceptive processing.

Given this predicted deep lamina-like identity, we next asked whether this brainstem territory also receives descending cortical input, a prominent feature of deep spinal laminae^48,49^. Since both the spinal cord and brainstem are known to receive cortical input^50,51^, we reasoned that comparing corticofugal projection patterns across these structures could provide an independent test of our predictions. To test this directly, we co-registered MERFISH assignments for all Rexed laminae with NeuN, corticofugal tract labeling from sensorimotor cortex, and IB4 immunoreactivity in spinal cord and brainstem sections **(Figure S11D)**.

As expected, in the spinal cord dorsal horn, corticospinal axons densely innervated lamina V/VI, consistent with its functional role as a major site of descending cortical modulation, sensory gating, and sensorimotor integration^48,49^. Corticospinal tract (CST) axons also innervated laminae III and IV, although to a lesser extent, and were not detected in laminae I/II (indicated by IB4-labeling). In addition, CST axons densely innervated lamina VII, consistent with its role in motor control and proprioceptive sensory modulation^52,53^. Consistently, this pattern extended into the caudal brainstem. Co-registration of anterograde tracing experiments with MERFISH spatial data demonstrated that CST axons densely mapped onto predicted lamina V-like regions, including the cuneate and external cuneate nuclei, consistent with the role of these regions in somatosensory integration and the role of cortical projections in tactile modulation^50^ (**Figure S11D**).

Lamina VII-like regions and their related supertypes mapped to ventral medullary reticular territories associated with reticulospinal control of posture and movement, including MDRNv, GRN, and MARN. These regions overlap major reticulospinal neuron-containing territories involved in locomotor regulation, postural control, and motor coordination^15,54^, consistent with the role of spinal lamina VII in premotor and motor-related processing. Similarly, we observed that descending cortical axons innervated lamina VII-like territories in both the spinal cord and brainstem, consistent with previous reports and supporting the idea that these regions share functional features with lamina VII of the spinal cord^51,53,54^.

Laminae VIII/IX, which are defined predominantly by skeletal motor neurons in the spinal cord, mapped to skeletal motor neuron populations in the hypoglossal nucleus (XII) which was primarily represented by alpha-motoneuron supertype 071 sMN Aox1 Glis3, consistent with the role of hypoglossal nucleus in controlling muscles of the tongue.

Finally, autonomic and viscerosensory spinal regions, including lamina X and the intermediolateral column (IML) mapped onto medullary autonomic centers such as the nucleus of the solitary tract, dorsal motor nucleus of the vagus (DMX), and nucleus ambiguous (Amb). This correspondence is consistent with the known role of these medullary nuclei in integrating visceral sensory information and coordinating autonomic motor output. In addition, this lamina X-like domain extended into the dorsal medullary reticular nucleus (MDRNd) and intermediate reticular nucleus (IRN), regions positioned between canonical sensory, autonomic, and motor brainstem centers. This broader distribution suggests that the conserved brainstem–spinal cord organization is not limited to somatic sensory and motor systems, but also encompasses circuits involved in visceral sensory processing, autonomic regulation, and integration between visceral and somatic functions **(Figure 3D; Figures S10, S11 A, B).**

### Molecular gradients organize cell types along the dorsoventral and mediolateral axes

The niche analysis demonstrated that cell type organization is conserved across the brainstem-spinal cord axis at the level of cellular neighborhoods. We next asked whether this spatial organization is reflected in the molecular profiles of individual cell types and whether specific marker genes encode positional information along the dorsoventral (D-V) and mediolateral (M-L) axes.

To quantify the spatial position of each supertype, we computed mean D-V and M-L distance scores from the Xenium and Merfish spatial data, placing each supertype in a two-dimensional spatial coordinate system independent of its molecular identity. Plotting supertypes in this D-V versus M-L space revealed a continuous and class-specific spatial organization (**Figure 4A**). GABAergic supertypes occupied a broad range of D-V positions, consistent with inhibitory interneurons being distributed throughout the dorsal-ventral extent of the spinal cord and brainstem. Glutamatergic supertypes showed a similarly broad distribution but with a tendency toward more dorsal positions, while motor neurons clustered at low D-V scores and intermediate M-L scores, reflecting their ventromedial localization. Cholinergic neurons occupied a restricted region of the spatial coordinate space consistent with their known medial to ventral distribution. Notably, this spatial organization was mostly consistent across brainstem and spinal cord sections (**Figure 4A**), with equivalent supertypes occupying equivalent spatial positions in both structures, reinforcing the conclusion that the organizational logic of the spinal cord extends continuously into the brainstem.

**Figure 4.**
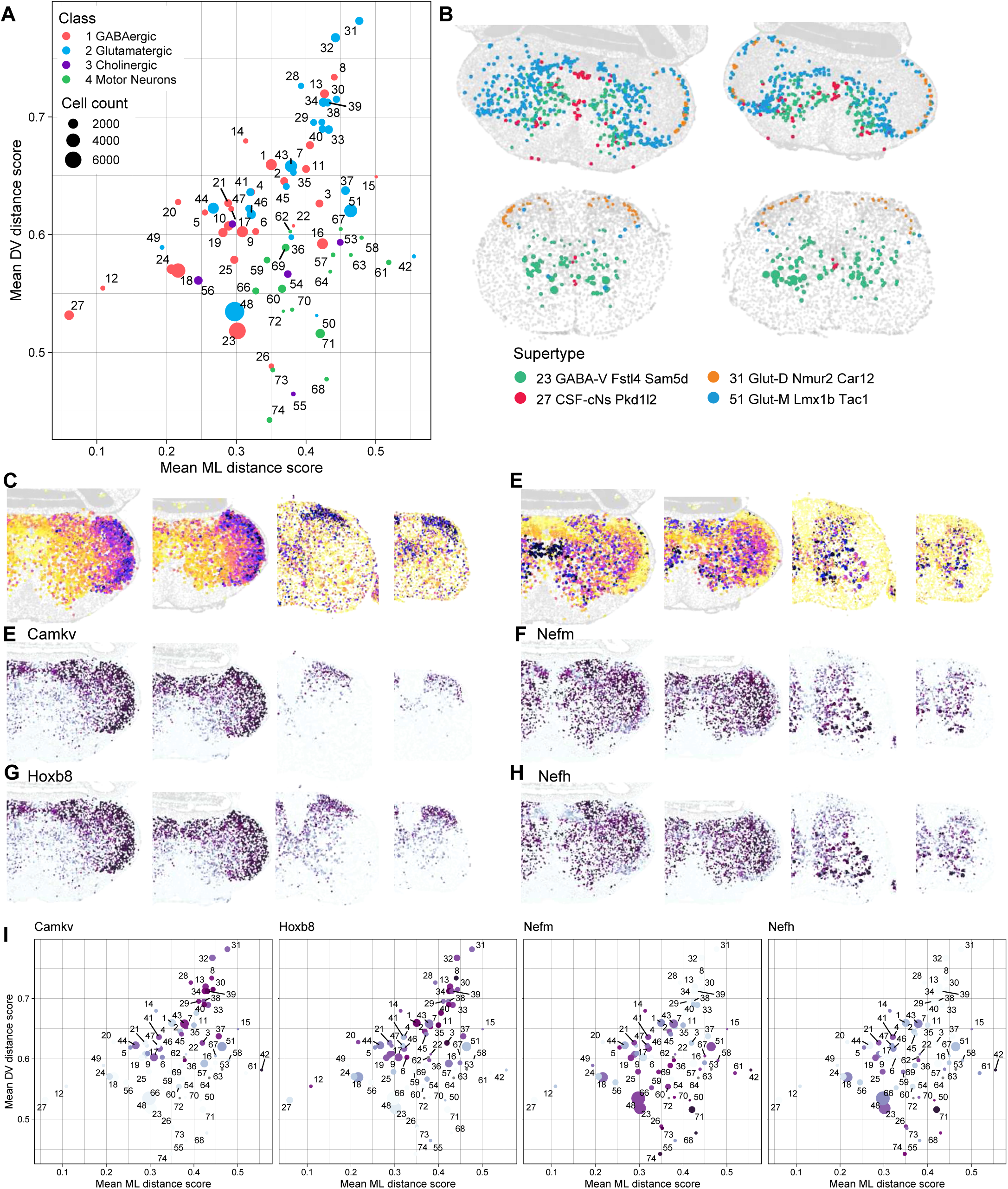
Spatial positioning scores and anatomical distribution of neuronal supertypes across the spinal cord. **(A)** Scatter plot of mean dorsoventral (DV) distance score versus mean mediolateral (ML) distance score for all neuronal supertypes. Each dot represents one supertype, colored by neurotransmitter class (pink: GABAergic; teal: Glutamatergic; blue: Cholinergic; red: Motor Neurons) and scaled by cell count (bubble size proportional to 2,000–6,000 cells). Supertype numbers are labeled adjacent to each dot. Supertypes occupying distinct spatial territories are dispersed across the plot, reflecting their anatomical segregation along the DV and ML axes. **(B)** Spatial transcriptomic maps showing the anatomical distribution of four representative supertypes projected onto coronal sections: 23 GABA-V Fst4 Sam5d (green), 27 CSF-cNs Pkd1l2 (pink), 31 Glut-D Nmur2 Car12 (orange), and 51 Glut-M Lmx1b Tac1 (blue). Each colored dot represents a single cell, overlaid on a gray reference tissue outline. **(C-D)** Spatial feature maps displaying gene module scores projected onto coronal tissue sections. Color gradients (purple–yellow) reflect module score intensity, with purple indicating higher scores. Panel (C) shows the DV (dorsoventral) NMF gene module score and panel (D) shows the VD (ventrodorsal) NMF gene module score across all profiled cells, capturing complementary spatial gradients of gene expression along the dorsoventral axis of the CNS. **(E-H)** Spatial feature maps showing the expression of *Camkv*, *Hoxb8*, *Nefm*, and *Nefh* projected onto coronal tissue sections. Color gradients reflect normalized expression levels across all profiled cells. **(I)** Scatter plots of mean DV versus mean ML distance score for all neuronal supertypes, reproduced from (A) and faceted by four landmark genes, *Camkv*, *Hoxb8*, *Nefm*, and *Nefh*. Each dot represents one supertype, colored by its mean expression level derived from single-nucleus RNA sequencing data, revealing how transcriptional identity of spatially positioned supertypes correlates with known regional and maturation markers along the dorsoventral and mediolateral axes.

To identify genes whose expression tracks the D-V spatial gradients independently of cell type identity, we performed spatially variable gene analysis on Xenium data using Moran’s I statistic, followed by Non-Negative Matrix Factorization (NMF) of cell-type-regressed expression residuals to identify co-varying gene programs representing continuous spatial gradients (see **Methods**). This approach yielded reproducible spatial gradient factors shared across datasets, each defined by a gene loading vector and a spatial score pattern (**Figure 4C,D**). Rather than assigning discrete spatial domains, NMF produced continuous, additive decompositions suited to the gradient structure of the data, with each factor representing an independent spatial program.

The spatial expression patterns of representative gradient genes were confirmed directly in Xenium tissue sections across brainstem and multiple spinal cord levels, demonstrating that these gradients are present in the tissue and reproducible across sections (**Figure 4E-H**). Factors with high D-V specificity included genes such as *Camkv* and *Hoxb8*, which showed strong dorsal enrichment across supertypes from multiple neuronal classes (Supp Figure Xe,g,i) whereas factors with V-D gradient specificity included genes like *Nefm* and *Nefh*, encoding medium and heavy neurofilament subunits, which were enriched in ventrally positioned supertypes (**Figure 4F,H,I**). Notably, these D-V gradient genes were expressed in a class-independent manner, both GABAergic and glutamatergic supertypes showed correlated expression with their D-V position, indicating that the D-V molecular gradient reflects a positional signal acting across cell type boundaries rather than a class-specific program (**Figure 4A,I**).

### Chromatin accessibility modules define conserved regulatory programs across brainstem and spinal cord neurons

The conserved spatial organization of neuronal types raises the question of what transcriptional regulatory programs underlie this conservation. Cell type identity in the central nervous system is established and maintained through precise transcriptional programs, in which combinations of transcription factors activate cell type-specific genes while repressing alternative fates. To address this, we leveraged our multiome dataset, which simultaneously profiles gene expression and chromatin accessibility in the same nuclei, alongside single-nucleus RNA-seq data that lack paired chromatin measurements. To integrate these modalities across all cells, we used MultiVI (see Methods)^55^, a probabilistic model that jointly embeds paired and unpaired cells into a shared latent space and imputes chromatin accessibility for expression-only cells (**Fig. S12A,B**). The resulting integrated representation captures both transcriptional and chromatin-level cell type structure: measured and imputed ATAC profiles were highly concordant across subclasses (**Fig. S12C,D**), and cell type identity could be recovered with high accuracy from accessibility alone, expression alone, or their combination (**Fig. S12E,F**), confirming that the multimodal integration faithfully preserves cell type information from both data modalities. By integrating single-cell transcriptomic profiles with these chromatin accessibility maps, we reconstructed cell type–specific cis-regulatory landscapes, enabling us to move beyond classification based on gene expression alone.

To systematically identify the chromatin regulatory programs underlying each neuronal cell type, we applied weighted gene co-expression network analysis to pseudobulk chromatin accessibility profiles computed across all neuronal subclass by region combinations. This allowed us to identify peak co-accessibility modules that each represent a coordinately regulated set of cis-regulatory elements (**Fig. 5A, Fig. S13**). Each module was characterized by a module membership score (kME) that quantifies how strongly each peak contributes to the module, and peak-to-gene correlation analysis linked module peaks to their putative target genes, revealing the downstream transcriptional programs regulated by each cis-regulatory module (**Fig. S13**). Crucially, the accessibility of these modules was highly cell-type-specific, each neuronal subclass showed a distinct pattern of module accessibility that was consistent across brainstem and spinal cord pseudobulks, demonstrating that the chromatin landscape faithfully encodes cell type identity independently of anatomical region (**Fig. 5A**).

**Figure 5.**
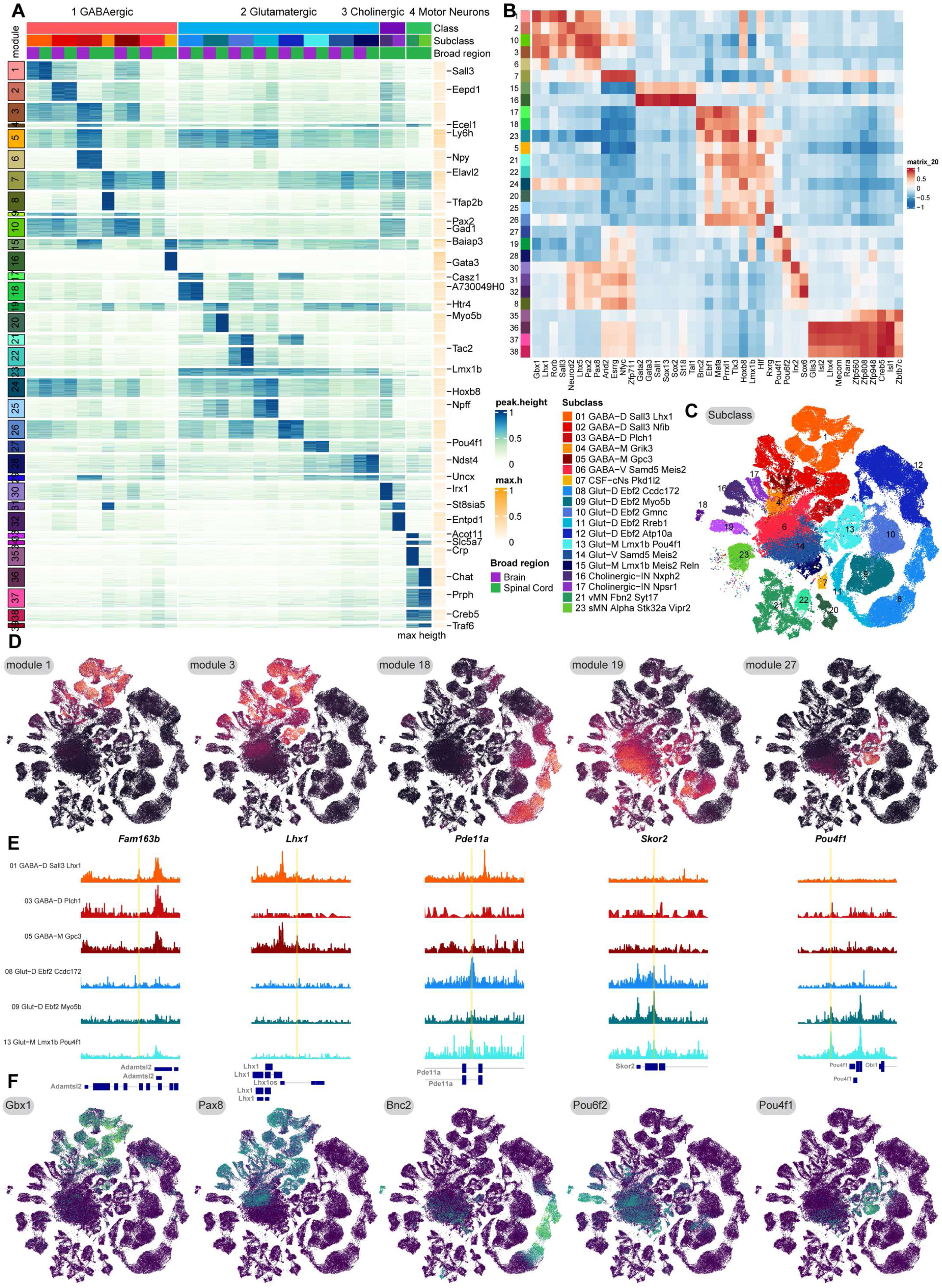
Chromatin accessibility modules reveal shared cell-type-specific regulatory programs in brainstem and spinal cord neurons. **(A)** Heatmap of chromatin accessibility across peak co-accessibility modules (rows) and neuronal subclass × region pseudobulks (columns). Peaks are ordered by module assignment and ranked by module membership score (kME). Columns are split by class and ordered by subclass. Accessibility values are row-normalized (0–1) per peak. Column annotations indicate cell class, subclass, and region. **(B)** Heatmap showing Pearson correlation between peak module accessibility scores and transcription factor motif enrichment across pseudobulks. Rows are peak modules from panel a and columns are TF motifs. Hierarchical clustering reveals groups of modules sharing common regulatory TFs. **(C)** UMAP of the MultiVI joint embedding for neurons, colored by subclass. **(D-F)** For each of 5 focal subclass × module combinations: (D) UMAP of neuronal cells, as in panel c, colored by chromatin module score, computed as the kME-weighted mean imputed accessibility across module peaks; (E) ATAC-seq signal tracks showing pseudobulk accessibility at the top peak-gene linked locus, with the focal peak highlighted in gold; (F) UMAP colored by expression of the module-associated transcription factor. Signal tracks are shown at a shared scale across all subclasses within each column.

To identify the transcription factors driving each chromatin module, we performed TF motif enrichment analysis on module peaks using JASPAR2020 vertebrate motifs, computing the mean motif accessibility score across module peaks relative to background (**Fig. 5B**). This analysis revealed clear TF-module associations consistent with known biology. Modules specific to GABAergic dorsal interneurons were enriched for *Lhx1*, *Pax2*, and *Gbx1* motifs, glutamatergic dorsal neuron modules showed enrichment for *Tlx3* and *Lmx1b* motifs, and motor neuron modules were enriched for *Creb5* and *Isl1* motifs. To validate these motif-based predictions, we confirmed that the corresponding TF genes were specifically expressed in the relevant neuronal subclasses using our single-cell RNA-seq data (**Fig. S14A**), directly linking chromatin accessibility patterns to the transcriptional regulators that establish them. This revealed a striking pattern of combinatorial TF expression that closely mirrors the module structure identified from chromatin accessibility. In GABAergic populations, *Gbx1*, *Lhx1*, *Sall3*, *Lhx5*, *Pax8*, and *Pax2* were enriched across dorsal GABAergic subclasses (01-03), while progressively more ventral subclasses showed enrichment for distinct TF combinations including *Meis2*, *Nfia*, *Nfix*, *Npas1*, *Foxj1*, *Gata3*, and *Pitx2*. Glutamatergic subclasses showed complementary enrichment for *Tlx3*, *Ebf2*, *Lmx1b*, *Onecut1*, *Onecut2*, and *Pou4f1*, with dorsal and ventral subclasses expressing largely non-overlapping TF combinations (**Fig. S14A**). Motor neuron subclasses were distinguished by *Isl1*, *Isl2*, *Sox2*, and *Phox2b* expression, with visceral and skeletal motor populations showing distinct profiles. The tight correspondence between motif enrichment in accessible chromatin and TF gene expression in the matching subclasses confirms that the identified modules reflect genuine cell-type-specific regulatory programs, and that the transcription factors predicted from chromatin data are actively expressed and likely to be functional regulators of those programs rather than passive binding site occupants.

To visualize these regulatory programs at single-cell resolution, we computed per-cell module scores from imputed chromatin accessibility values as the kME-weighted mean accessibility across module peaks and projected these onto the MultiVI joint embedding of all neurons (**Fig. 5C,D**). Focusing on six representative neuronal subclasses spanning the dorsal–ventral and medial–lateral axes, we found that these cell-type-specific chromatin modules are strongly conserved between brainstem and spinal cord. For instance, GABAergic dorsal interneurons (01 GABA-D Sall3 Lhx1) exhibited specific accessibility at Sall3-linked peaks with enrichment for Lhx1 motifs in both regions, while glutamatergic medial interneurons (13 Glut-M Lmx1b Pou4f1) showed consistent accessibility at *Pou4f1*-linked peaks with enrichment for *Pou4f1* motifs across brainstem and spinal cord (**Fig. 5E,F**). Together, these results demonstrate that neuronal cell type identity is encoded in conserved chromatin regulatory programs maintained along the rostral–caudal axis of the central nervous system.

### Hox genes encode positional identity along the rostral-caudal axis

While cell type identity programs are conserved across regions (**Fig. 5**), the same neurons must also encode their anatomical position along the rostral-caudal axis. Hox genes are well-established determinants of rostral-caudal positional identity during development and maintain region-specific expression in adult neurons^56^. Whether this positional code is reflected in cell-type-specific chromatin accessibility landscapes at single-cell resolution across the combined brainstem-spinal cord axis, however, has not been directly examined. To address this, we examined Hox-associated regulatory landscapes across the brainstem and spinal cord, asking whether positional identity is encoded alongside cell type at single-cell resolution. We applied the same peak co-accessibility module framework to identify modules whose top peaks by kME overlapped with the four *Hox* gene clusters in the mouse genome (*Hoxa*: chr6, *Hoxb*: chr11, *Hoxc*: chr15, *Hoxd*: chr2), identifying 15 Hox-associated chromatin modules (**Fig. 6A**).

**Figure 6.**
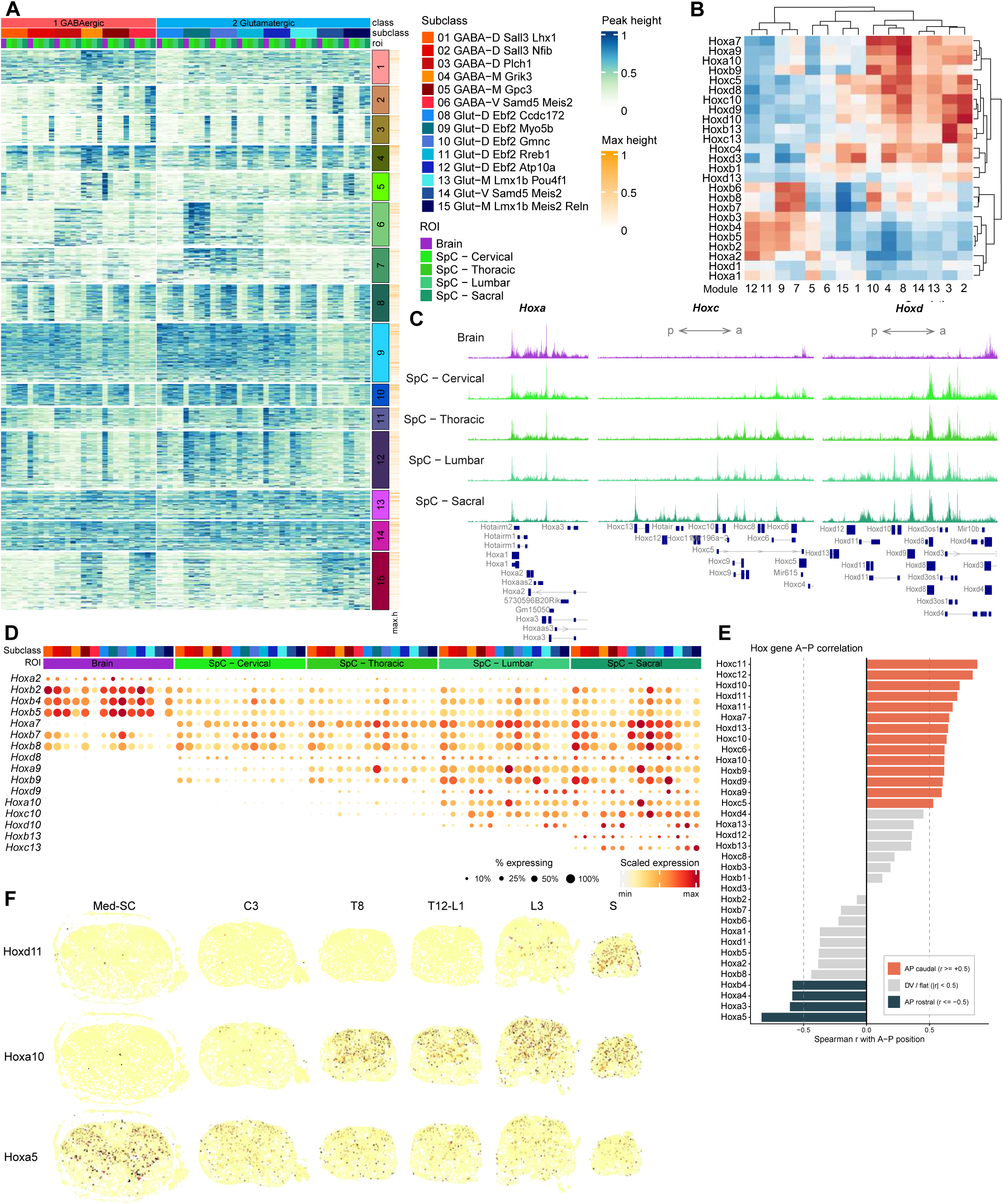
Hox genes encode positional identity along the rostral-caudal axis of the brainstem and spinal cord. **(A)** Heatmap of chromatin accessibility across Hox-associated peak modules (rows) and neuronal subclass × region pseudobulks (columns). Columns are ordered along the rostral-caudal axis (brain, cervical, thoracic, lumbar, sacral) and split by GABAergic and Glutamatergic class. Accessibility values are row-normalized (0–1) per peak. Column annotations indicate cell class, subclass, and region. **(B)** Heatmap showing Pearson correlation between Hox gene expression levels (rows) and peak module accessibility scores (columns) across pseudobulks. Both rows and columns are hierarchically clustered, revealing groups of Hox genes that are co-regulated with specific chromatin modules and separating rostral- from caudal-enriched Hox genes. **(C)** Dotplot showing Hox gene expression per neuronal subclass and region group. Dot color reflects row-normalized mean expression (min–max per gene) and dot size reflects the percentage of expressing cells. Genes are ordered by paralog group number (rostral to caudal). **(D)** ATAC-seq signal tracks showing mean pseudobulk accessibility averaged across all neuronal subclasses for three *Hox* gene cluster windows. The left, middle and right panels show the *Hoxa* cluster, the *Hoxc* cluster, and *Hoxd* cluster respectively. Each row represents one of five regions along the rostral-caudal axis (brain, cervical, thoracic, lumbar, sacral). Focal peaks are highlighted in gold. **(E)** Spatial distribution of *Hoxd11*, *Hoxa10*, and *Hoxa5* expression measured by Xenium spatial transcriptomics across six transverse spinal cord sections spanning the rostral-caudal axis (medulla-spinal cord border [Med-SC], cervical [C3], thoracic [T8], thoracic-lumbar [T12-L1], lumbar [L3], and sacral [S]). Each row shows one *Hox* gene across the six sections, revealing the spatial restriction of *Hox* gene expression to specific axial levels. *Hoxd11* expression is restricted to lumbar and sacral levels, *Hoxa10* is expressed from thoracic-lumbar to sacral, and *Hoxa5* expression extends from cervical to thoracic levels, directly confirming the rostral-caudal chromatin accessibility gradients identified in the ATAC-seq data at cellular and tissue resolution. **(F)** Spearman correlation between *Hox* gene expression and rostral-caudal position, computed from Xenium spatial transcriptomic data across the six sampled spinal cord levels. Genes are ordered by correlation coefficient. Red bars indicate genes with significant positive correlation with caudal position (Spearman r > 0.5, FDR < 0.05), grey bars indicate genes with significant negative correlation with caudal position (rostral-enriched, FDR < 0.05), and dark grey bars indicate non-significant correlations. Higher-numbered *Hox* paralog groups show the strongest positive correlation with caudal position, confirming the principle of *Hox* collinearity and directly linking the chromatin accessibility and gene expression gradients identified in panels A and B to the precise spatial distributions of *Hox* genes along the spinal cord axis.

Some of these modules showed a striking rostral-caudal gradient in chromatin accessibility across the five anatomical regions sampled - brainstem, cervical, thoracic, lumbar, and sacral spinal cord (**Fig. 6A**). To validate that these chromatin accessibility gradients are driven by the corresponding Hox transcription factors, we performed TF motif enrichment analysis on Hox module peaks (**Fig. 6B**). We confirmed that modules with high accessibility in brainstem and cervical neurons were enriched for peaks mapping to lower-numbered Hox paralog groups, i.e. *Hoxa2*-*Hoxa7* and *Hoxb2*-*Hoxb8*, while modules with high accessibility in lumbar and sacral neurons were enriched for peaks mapping to higher-numbered paralog groups, *Hoxa9*-*Hoxa10*, *Hoxb9*-*Hoxb13*, *Hoxc10*-*Hoxc13*, and *Hoxd9-Hoxd10*. ATAC-seq signal tracks at representative loci across all four Hox clusters confirmed these accessibility gradients at single-locus resolution, with peaks near lower-numbered Hox genes most accessible in brainstem and cervical pseudobulks, while peaks near higher-numbered Hox genes most accessible in lumbar and sacral pseudobulks (**Fig. 6C, Fig S15**).

To directly connect these chromatin-level findings to gene expression and spatial distributions, we examined Hox gene expression at both the transcriptomic and spatial levels. In our single-cell RNA-seq data, Hox gene expression showed a clear rostral-caudal gradient concordant with the chromatin accessibility patterns in which higher-numbered paralogs are expressed progressively more caudally along the body axis (**Fig. 6D**). To validate these findings at cellular and tissue resolution, we leveraged Xenium spatial transcriptomic data from transverse spinal cord sections spanning six axial levels, from medulla-spinal cord border, cervical, thoracic, lumbar, to sacral spinal cord. The spatial distribution of individual Hox genes precisely recapitulated the predicted gradients, *Hoxa5* expression was restricted to cervical and thoracic levels, *Hoxa10* extended from thoracic-lumbar to sacral, and *Hoxd11* was restricted to lumbar and sacral levels (**Fig. 6E**). Quantification of Hox gene expression across all sampled axial levels confirmed that higher-numbered Hox paralogs showed the strongest positive Spearman correlation with caudal position, directly confirming the principle of Hox collinearity in adult spinal cord neurons at spatial resolution (**Fig. 6F**). Importantly, the expression gradient was broadly shared across neuronal subclasses, demonstrating that the Hox positional code operates as a general mechanism across neuronal cell types rather than being restricted to specific subclasses.

Notably, not all *Hox* genes behave as purely positional markers. Examining *Hox* gene expression across subclasses and regions revealed that a subset, including *Hoxb3*, *Hoxb4*, *Hoxb6*, *Hoxb7*, and *Hoxb8,* showed expression primarily determined by subclass identity rather than anatomical region, while others including *Hoxc10*, *Hoxb13*, *Hoxc13*, *Hoxa10*, and *Hoxd10* varied primarily along the rostral-caudal axis across all cell types (**Fig. 6D, Fig. S45B**). To formally quantify this distinction, we decomposed the variance of each *Hox* gene’s expression into components explained by cell type identity and anatomical region respectively (**Fig. S14C**). This analysis revealed a clear separation of *Hox* genes into three groups: cell-type-specific *Hox* genes (*Hoxb3*, *Hoxb4*, *Hoxb6*, *Hoxb7*, *Hoxb8*) whose expression is primarily determined by subclass identity; region-specific *Hox* genes (*Hoxc10*, *Hoxb13*, *Hoxc13*, *Hoxa10*, *Hoxd10*, *Hoxa9*, *Hoxb9*, *Hoxa7*, *Hoxd8*) whose expression varies primarily along the rostral-caudal axis across all cell types; and a smaller mixed group (*Hoxb2*, *Hoxb5*, *Hoxd3*, *Hoxc4*) contributing to both axes. This variance decomposition provides a quantitative basis for the distinction between *Hox* genes integrated into cell-type identity programs and those encoding positional identity, demonstrating that these two modes of *Hox* function are encoded in largely non-overlapping sets of family members rather than being properties of all *Hox* genes simultaneously.

Together, these results reveal a multi-layered regulatory architecture in which conserved chromatin programs encode cell type identity across the entire neuraxis, while a superimposed *Hox* positional code continuously varies from brainstem to sacral spinal cord. The *Hox* family occupies a distinctive position in this architecture. Certain family members are incorporated into the cell-type-specific regulatory programs that maintain neuronal identity in the adult, while others function as positional markers whose chromatin accessibility and expression track rostral-caudal location. This dual organization provides a mechanistic framework for understanding how the same neuronal cell types can be maintained across anatomically distinct regions while retaining region-specific functional properties.

### Molecular taxonomy identifies candidate brainstem homologs of spinal motor neuron subtypes

A key prediction of a unified brainstem-spinal cord taxonomy is that molecularly defined cell types should identify functionally equivalent populations regardless of their anatomical location. Motor neurons provide a direct test of this prediction, as the brainstem contains well-characterized cranial motor nuclei whose functional equivalents in the spinal cord are well-established. To directly test this prediction, we focused specifically on cranial motor nuclei. These are the brainstem structures containing somatic and visceral motor neurons with well-established functional equivalents in the spinal cord. Because deep sampling of cranial motor nuclei requires dedicated brainstem datasets, we supplemented our atlas with motor neuron populations from the whole-brain single-cell atlas of Yao et al. (subclasses 316 and 317), which provides dense coverage of brainstem motor populations that complement the spinal cord-focused sampling of our primary dataset. We constructed a joint embedding of all motor neuron supertypes from both datasets and validated spatial assignments of brainstem motor populations using *Chat* and *Isl1* co-expression in our MERFISH data (**Figure S16A-C**). Skeletal alpha motor neurons, subclass 23 sMN Alpha Stk32a Vipr2, which in the spinal cord innervate extrafusal muscle fibers across fast-fatiguing and fatigue-resistant pools, mapped in the brainstem to the hypoglossal nucleus (XII), the facial motor nucleus (VII), and the motor nucleus of the trigeminal (V) (**Figure S16B**). These are precisely the brainstem nuclei containing somatic lower motor neurons controlling cranial musculature, the hypoglossal nucleus controlling tongue musculature, the facial nucleus controlling muscles of facial expression, and the trigeminal motor nucleus controlling muscles of mastication. The correspondence confirms that the molecular identity of alpha motor neurons is conserved across the neuraxis irrespective of their specific target musculature, and that the same transcriptional programs that define skeletal motor neuron identity in the spinal cord are deployed in cranial motor nuclei controlling functionally analogous but anatomically distinct muscle groups.

Gamma motor neuron supertypes, subclass 24 sMN Gamma Htr1d Npas1, which in the spinal cord innervate intrafusal muscle spindle fibers and regulate proprioceptive sensitivity, showed spatial correspondence with the midbrain reticular nucleus and inferior colliculus^12^. While these structures lie outside the strict brainstem-spinal cord axis that is the primary focus of this study, this mapping is consistent with the known expression of gamma motor neuron marker genes in reticular regions and suggests that the molecular programs defining spindle-regulating motor identity extend rostrally beyond the brainstem. Whether these populations represent true functional homologs of spinal gamma motor neurons or reflect shared developmental programs without equivalent peripheral connectivity remains an open question that warrants further investigation.

Visceral motor neuron supertypes were enriched in the dorsal motor nucleus of the vagus (DMX) and nucleus ambiguus (AMB) in our MERFISH spatial data (**Figure S16B**), both established sources of parasympathetic preganglionic output in the brainstem. Cells in these regions assigned to visceral motor neuron supertypes expressed Chat with low or absent Isl1 (**Figure S16C**), consistent with the molecular profile of autonomic preganglionic neurons. We note that the brainstem contains exclusively parasympathetic preganglionic neurons, whereas the spinal cord intermediolateral column contains primarily sympathetic preganglionic neurons, with parasympathetic outflow restricted to the sacral segments. The molecular correspondence observed between brainstem visceral motor populations and spinal visceral motor neuron supertypes therefore reflects shared cholinergic preganglionic identity rather than equivalence of autonomic output type.

Together, these findings establish that the molecular programs defining somatic and visceral motor neuron identity in the spinal cord are conserved in anatomically appropriate medullary populations, as validated by *in situ* marker co-expression in our MERFISH spatial dataset. Whether these molecularly defined brainstem populations are functionally equivalent to their spinal counterparts, innervating equivalent targets with equivalent physiological properties, remains to be established by connectivity and physiological studies.

### Molecular taxonomy identifies conserved cell types with defined projection identities across the brainstem-spinal cord axis

The molecular conservation of cell types across the brainstem and spinal cord predicts that neurons sharing supertype identity at different levels of the neuraxis should participate in equivalent circuit roles. To test this prediction and connect the molecular taxonomy to circuit function, we examined both the marker gene expression of spinal cord supertypes and their correspondence with retrogradely labeled populations from two independent tracing datasets.

The spinal cord contains molecularly defined populations whose marker gene expression predicts specific ascending projection identities consistent with known circuit anatomy. Supertype 045 (Glut-M Prox1 Pdyn) co-expresses *Prox1*, *Tacr1*, and *Pdyn*, consistent with a lamina I spinoparabrachial population relaying nociceptive information to the lateral parabrachial nucleus. Supertype 051 (Glut-M Lmx1b Tac1) occupies laminae V-VII and co-expresses *Lmx1b* and *Tac1* (**Fig. S17A**), consistent with wide-dynamic-range projection neurons of the anterolateral system ^9,37^. Notably, a subgroup within supertype 051 abundantly expresses *Phox2a*, a transcription factor that defines the developmental origin of the anterolateral system in mice and humans^57^, providing direct molecular evidence for ascending projection identity within this population. The presence of *Phox2a*-negative cells within the same supertype suggests molecular heterogeneity that may reflect a mixed population of projection and local interneurons at this taxonomic resolution. Supertype 048 (Glut-V Htr4 Cdh9) expresses *Shox2* and corresponds to the Excit-35 cluster of Russ et al.^9^ (**Fig. S6B, Fig. S17A**), which includes *Shox2*-expressing V2a interneurons. Within this population, a subset participates in spinocerebellar projections providing proprioceptive feedback to the cerebellum^19,58^ as confirmed by cerebellar retroSeq labeling in our dataset (CB, n=285). The broader population, however, is a key component of the locomotor CPG, the V2a excitatory interneurons that coordinate rhythmic motor output^59,60^, suggesting that this supertype encompasses neurons that both drive local spinal motor circuits and relay proprioceptive information to supraspinal centers. Together these three populations represent candidate ascending sensory and proprioceptive relay systems of the spinal cord, each identifiable by a distinct molecular signature in the adult tissue.

To identify which molecularly defined populations project from brain to spinal cord, we integrated the Winter et al.^15^ retrograde tracing dataset, in which brain neurons were retrogradely labeled from cervical (GFP) and lumbar (RFP) spinal cord injections and subsequently sequenced. Labeled cells were assigned to supertypes based on their transcriptional profiles in the shared latent space, directly linking molecular identity to projection target (**Fig. S17B**). The largest population was supertype 048 (Glut-V Htr4 Cdh9), with 1511 cervical-projecting and 824 lumbar-projecting labeled cells. The near-equal labeling from both cervical and lumbar injections indicates that this population provides broad descending excitatory drive across spinal levels, consistent with a reticulospinal identity^15^. Supertype 023 (GABA-V *Fstl4 Sam5d*) was the second most abundant, with 338 cervical- and 206 lumbar-projecting cells, representing the inhibitory counterpart. This is a glycinergic/GABAergic descending population providing broad inhibitory modulation at both spinal levels.

Cross-referencing supertype assignments with Winter et al.^15^ cluster identities revealed that the molecular identity of descending projection neurons is conserved with specific spinal cord interneuron supertypes, providing, for the first time, a direct link between the transcriptional identity of brainstem projection neurons and the spinal interneuron populations they most closely resemble (**Fig. S17C**). Supertype 009 (GABA-D Zic1) showed a Jaccard overlap of 0.967 with Winter et al.^15^ cluster 56 (MED-Lhx1/5-Glyc-Penk), a population that was identified as projecting predominantly to cervical spinal cord but whose relationship to spinal interneuron classes was previously unknown. The near-complete molecular correspondence established here identifies this descending population as the brainstem homolog of a specific enkephalin-expressing glycinergic spinal interneuron. This is consistent with a role in descending inhibitory control of upper limb sensorimotor circuits and nociceptive sensory processing. Similarly, supertype 044 (Glut-M Cep112 Sntg2) showed similarly high overlap (Jaccard = 0.844) with Winter et al.^15^ cluster 46 (MED-Lmx1b-Nfib), a glutamatergic medullary population projecting to cervical cord, representing a distinct descending excitatory system targeting intermediate zone sensorimotor circuits. Together these correspondences demonstrate that the descending control system is not composed of cell types unique to the brainstem but rather deploys the same molecular programs as the spinal cord interneurons it innervates, a principle of circuit organization that could not be identified from either dataset alone.

To characterize the full projection breadth of these populations, we integrated an unpublished medullary retroSeq dataset in which cells were isolated from medulla following retrograde injections at multiple target sites spanning spinal cord and brain. Labeled cells were again assigned to supertypes in the shared latent space (**Fig. S17D**). Supertype 048 contained retrogradely labeled cells from spinal cord (SC, n=812), medulla (MY, n=840), cerebellum (CB, n=285), midbrain (MB, n=83), and thalamus (TH, n=47). Supertype 023 showed an equally broad profile: SC (n=651), MY (n=720), CB (n=220), MB (n=33), and thalamus (n=11). These projection profiles reveal that supertypes 048 and 023 contain subpopulations with diverse long-range projection targets, collectively spanning spinal cord, cerebellum, thalamus, and midbrain circuits. Whether individual neurons within these supertypes project to multiple targets simultaneously, as would be expected of canonical reticulospinal neurons, remains to be determined by single-cell morphological reconstruction^61^.

Supertype 024 (GABA-V Nfix Prox1) showed a distinct profile dominated by cerebellar (CB, n=70) and medullary (MY, n=171) labeling with modest spinal cord projection (SC, n=32), consistent with a population participating primarily in cerebellar-brainstem circuits. The V0c partition cell supertypes 052 (Cholinergic-IN Otof Piezo2) and 054 (Cholinergic-IN Sox6) showed labeling exclusively from medullary injections with no spinal cord projection, suggesting these populations function as local modulatory neurons at the brainstem level rather than long-range projection cells.

Together, these findings demonstrate that conserved cell type programs are deployed at multiple levels of the neuraxis in circuit-appropriate roles. The same molecular identity that defines a spinal CPG excitatory interneuron (048) also defines a medullary reticulospinal neuron providing descending drive to that same circuit. Equally, the molecular signatures that define spinal ascending relay populations (045, 051) identify medullary relay neurons in the ascending pain and sensorimotor pathways. The molecular taxonomy therefore not only classifies cell types but predicts their connectivity, providing a unified framework for interpreting circuit organization across the full brainstem-spinal cord axis (**Fig. 7A**).

**Figure 7.**
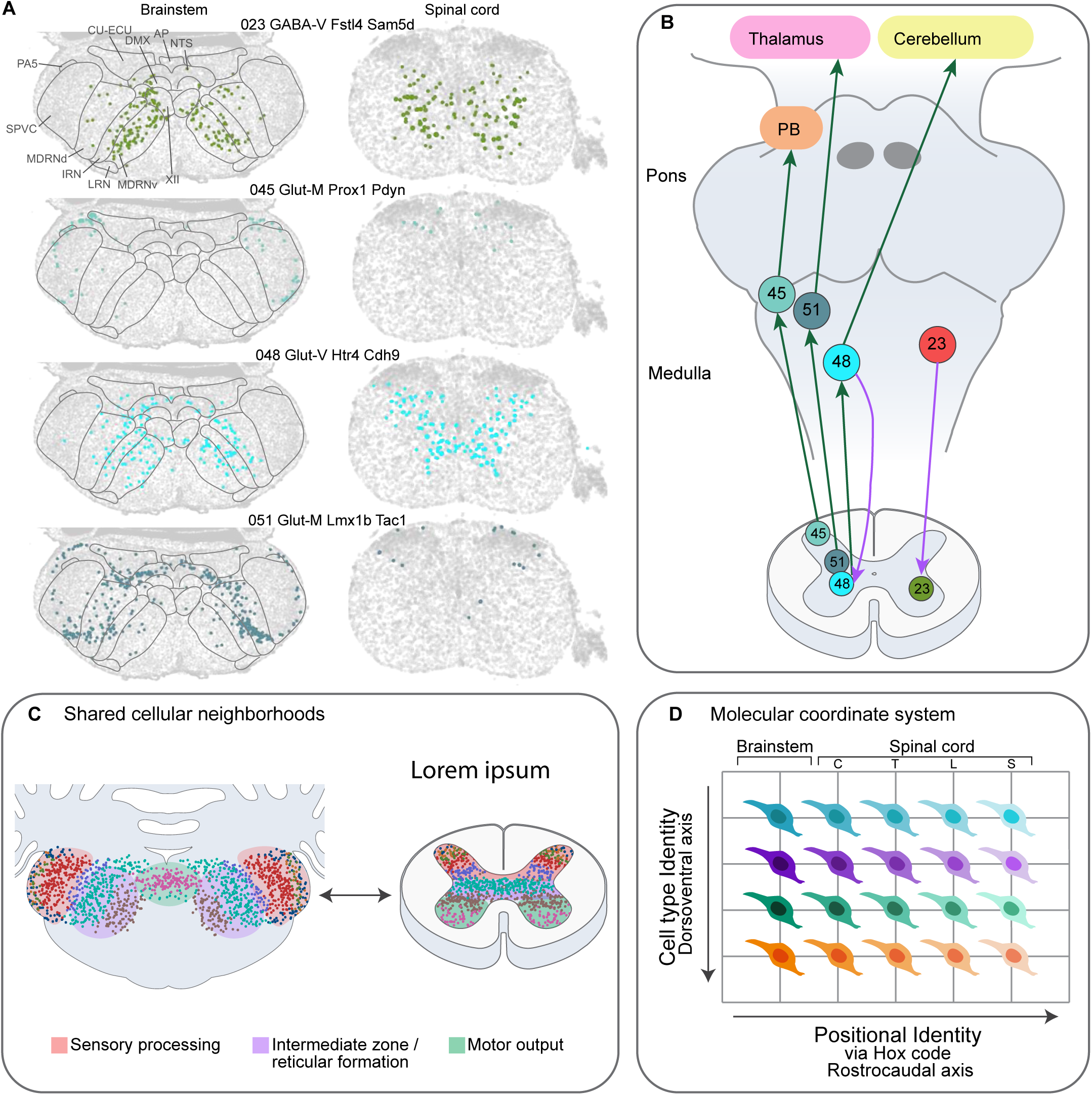
Summary of conserved cell type organization and connectivity across the brainstem-spinal cord axis. **(A)** Spatial distribution of molecularly defined ascending and descending projection supertypes across paired brainstem (left) and spinal cord (right) sections. Each row shows cells assigned to a single supertype overlaid on MERFISH (brainstem) and Xenium (spinal cord) spatial transcriptomic sections. **(B)** Schematic of ascending and descending projection pathways identified by molecular taxonomy. Ascending supertypes 045 (spinoparabrachial candidate population, *Tacr1/Pdyn*), 051 (anterolateral system, *Lmx1b*/*Phox2a*), and 048 (spinocerebellar/CPG, *Shox2*) are shown in the spinal cord (bottom) and their relay populations in the medulla, with projections to parabrachial nucleus (PB), thalamus, and cerebellum respectively (green arrows). Descending supertype 023 (glycinergic/GABAergic reticulospinal) projects from medulla to spinal cord (purple arrow). Numbers indicate supertype identifiers. The same supertypes are present at both levels of the neuraxis, distinguished by their HOX positional code. **(C)** Shared cellular neighborhoods between brainstem (left, medullary cross-section) and spinal cord (right). Colored dots represent cell types occupying equivalent spatial positions in both structures, with functional zones indicated: sensory processing (pink), intermediate zone/reticular formation (purple), and motor output (green). **(D)** Molecular coordinate system governing neuronal identity across the neuraxis. Cell type identity (dorsoventral axis, rows) and positional identity via HOX code (rostral-caudal axis, columns) operate as largely orthogonal axes of specification. Columns span brainstem through cervical (C), thoracic (T), lumbar (L), and sacral (S) spinal cord levels. Each cell illustration represents a molecularly defined population sharing transcriptional identity across positions but distinguished by its HOX positional code.

## DISCUSSION

Here we present a comprehensive molecular atlas of the mouse spinal cord and related regions in the brainstem, establishing a unified neuronal taxonomy that spans the rostral-caudal extent of both structures. By integrating single-nucleus RNA sequencing with spatial transcriptomics and chromatin accessibility profiling, we show that the brainstem and spinal cord share a common organizational logic at multiple levels: the spatial arrangement of cell types, the molecular gradients that underlie that arrangement, and the regulatory programs that define cell type identity. These findings reframe the relationship between the brainstem and spinal cord from two distinct anatomical structures with separate research traditions to a single integrated sensorimotor axis organized according to conserved cellular and molecular principles.

### A unified molecular reference across the brainstem-spinal cord axis

We generated a new single-nucleus multiome dataset profiling chromatin accessibility and gene expression from the adult mouse spinal cord, which we integrated with previously published single-cell datasets^9,12,13,16–18^ to create a unified reference atlas spanning the spinal cord and brainstem. Previous single-cell atlases have characterized either structure in isolation, producing classification schemes that are difficult to compare across studies and that obscure potential homologies between brainstem and spinal cord populations. The integrated taxonomy reveals that the transcriptional diversity of neurons in both structures can be organized into a coherent hierarchy of classes, subclasses, and supertypes grounded in biology rather than imposed by anatomical boundaries. This enables the direct cross-comparison of molecular signatures, improving annotation consistency, and facilitating the identification of conserved and region-specific gene regulatory programs.

The correspondence between molecularly defined supertypes and previously characterized functional populations, including pain-gating interneurons, locomotor CPG components, and autonomic preganglionic neurons, demonstrates that the taxonomy captures biologically coherent divisions with direct functional correlates. This alignment provides a molecular definition for populations that have historically been characterized through physiology, connectivity, or developmental lineage, establishing a bridge between the molecular and functional frameworks that have previously been developed independently for these circuits. Several cell types previously described as region-specific are in fact shared across the brainstem-spinal cord axis^11,14^. This suggests that functional studies in the spinal cord may be more directly relevant to brainstem circuits than previously appreciated and raises the possibility that findings about circuit dysfunction in spinal cord injury or brainstem disorders are more broadly applicable across the axis than previously recognized.

The spinal cord and brainstem, while sharing a common organizational logic, exhibit profound functional specialization along the rostrocaudal axis. The brainstem coordinates cranial nerve functions including facial movement, swallowing, and balance, cervical spinal cord segments control upper limb movements, thoracic segments innervate the trunk for respiration, and lumbar segments coordinate lower limb function. By sampling across the full rostrocaudal extent of both structures, this atlas addresses a longstanding question: whether regional functional specialization reflects distinct cell types in different segments or the same cell types with regionally tuned molecular properties. The answer, as discussed below, is largely the latter.

### Conservation of laminar organizational logic across the neuraxis

One of the most striking findings of this study is that the laminar organizational logic of the spinal cord is mostly conserved in the brainstem (**Fig. 7B**). The niche analysis shows that cell types occupying equivalent spatial positions in the spinal cord co-occur in shared cellular neighborhoods in the brainstem, allowing the Rexed laminar framework to be extended into a structure that lacks a continuous laminar cytoarchitecture. The correspondence between spinal laminae and specific brainstem nuclei is anatomically precise and functionally coherent. Superficial nociceptive laminae align with the superficial trigeminal nucleus (SPVC), intermediate sensorimotor laminae align with reticular structures (including GRN, MDRN, IRN, PARN, and PGRN) involved in motor coordination, and autonomic laminae align with medullary autonomic centers including the nucleus tractus solitarius (NTS) and dorsal motor nucleus of the vagus (DMX)^37,62^.

This conservation is not simply a reflection of shared developmental origin, since the brainstem and spinal cord derive from distinct rhombomeric and spinal progenitor domains respectively^63,64^. However, both structures share a continuous rostral-caudal patterning system, including the HOX code that extends across the brainstem-spinal cord boundary, suggesting that the laminar organizational logic may reflect a deeply conserved feature of the vertebrate sensorimotor axis that is elaborated differently in different regions rather than implemented independently. Whether this reflects duplication and divergence of a common ancestral progenitor pool or convergent deployment of shared transcription factor programs in distinct developmental contexts remains an open question, but the molecular correspondence identified here provides a framework for addressing it. The observation that the lamina IV-like domain appears broader in the brainstem than in the spinal cord, potentially reflecting the expanded facial tactile processing demands of a whisker-dependent rodent, suggests that while the organizational logic is conserved, the relative weighting of different functional domains can be modified in response to peripheral specialization. This raises the interesting question of whether the laminar organizational logic is similarly conserved across species with different sensory specializations, which could be addressed by applying the same niche framework to datasets from other vertebrates.

Motor neurons provide a clear demonstration of conservation across the brainstem-spinal cord boundary. Somatic motor neuron classes show strong molecular correspondence between spinal motor neurons and cranial motor nuclei, including the facial, abducens, and hypoglossal nuclei, indicating that core transcriptional programs defining somatic motor identity are conserved across axial levels^65,66^. Visceral motor neuron populations likewise exhibit conserved molecular organization across brainstem and spinal autonomic systems, with parasympathetic preganglionic neurons in the dorsal motor nucleus of the vagus and sacral spinal cord, and sympathetic preganglionic neurons in the thoracolumbar spinal cord reflecting a shared autonomic motor framework^65,67^.These correspondences have practical implications for motor neuron disease. Conditions such as amyotrophic lateral sclerosis, spinal muscular atrophy, and bulbar palsy affect motor neurons across the neuraxis, and the molecular basis of differential vulnerability remains poorly understood. The shared molecular taxonomy of spinal and cranial motor neurons provides a framework for investigating which programs are associated with vulnerability or resilience, and whether therapeutic strategies developed for spinal motor neurons extend to cranial motor populations.

### A molecular coordinate system for positional identity

The identification of a continuous molecular gradient along the dorsoventral axis adds a mechanistic dimension to the spatial organization observed in the niche analysis. The D-V gradient defined by opposing expression of *Hoxb8* dorsally and neurofilament coding genes ventrally operates across neuronal classes. This suggests that the gradient reflects a positional coordinate system that is independent of cell type identity and acts as a spatial reference frame within which cell type diversity is organized. Identifying these gradients required spatially variable gene analysis followed by NMF decomposition, an approach well suited to this problem because it produces continuous additive decompositions rather than discrete cluster assignments, capturing the graded nature of positional identity where boundaries between functional domains are gradients rather than sharp transitions.

These molecular gradients find their regulatory counterpart in the chromatin accessibility data. Cell-type-specific patterns of chromatin accessibility, linked to gene expression through peak-to-gene analysis, point to the transcription factors and regulatory elements that actively maintain neuronal identity in the adult tissue, as opposed to those whose expression simply reflects developmental history without ongoing regulatory activity. *Hox* transcription factors occupy a distinctive position in this landscape. Unlike transcription factors that vary primarily by cell type or primarily by region, individual *Hox* genes show varying degrees of cell-type-specificity and regional restriction, with some showing strong subclass-specific accessibility and others tracking rostral-caudal position across cell types. This pattern suggests that the *Hox* family does not encode a single axis of identity but instead participates in the regulatory definition of both what a neuron is and where it is. So, a dual involvement that connects the positional gradients identified at the expression level to the cell-type-specific regulatory programs that maintain neuronal identity in the adult.

That *Hox* genes remain active in mature neurons is itself notable. Their persistence in adult chromatin accessibility programs is consistent with a terminal selector role, in which transcription factors first deployed during development must be continuously expressed to sustain the differentiated state of the cells they specify^68^. This has been shown most clearly in C. elegans, where terminal selectors must be continuously expressed to maintain neurotransmitter identity and other features of mature neurons^69^. The involvement of *Hox* genes in both cell-type-specific and positional programs suggests they serve a dual function in the adult nervous system. On one hand, they contribute to the stable maintenance of cell type identity, and on the other, they encode segment-specific positional information, with different *Hox* genes appearing to be specialized for each role^64,70^.

### Cell type and positional identity as partially separable axes

Taken together, the analyses presented here support a framework in which neuronal identity in the spinal cord and brainstem exists along two partially separable axes, a cell type axis defined by developmental specification programs and core functional properties, and a positional axis defined by rostrocaudal patterning genes and region-specific adaptations (**Fig. 6C**). Most cell types represent conserved functional units deployed along the rostrocaudal axis, with regional specialization arising through tuning of molecular programs rather than wholesale invention of new cell types. Positional information, encoded through *Hox* genes and additional patterning transcription factors, is layered atop core cell type programs, while region-specific expression of connectivity and neuromodulatory genes refines local circuit properties. This architecture reconciles an apparent contradiction between the conservation of cell types across regions and the clear functional specialization of spinal segments and brainstem nuclei. The same basic cellular components are deployed throughout the neuraxis, but their molecular properties and circuit integration are customized for regional function. This view is consistent with recent work in embryonic motor neurons showing that while adult motor neurons use diverse neuropeptide codes, the transcription factor–neuropeptide combinations observed in the embryo are largely distinct from those in adult subtypes^39,71^, suggesting that positional tuning of cell type programs is a dynamic process that continues beyond initial specification.

The relationship between molecular identity and circuit function at the level of individual synaptic connections is not addressed by the current dataset. Understanding how the spatial organization characterized here gives rise to the computational properties of sensorimotor circuits will require integration with connectomic, electrophysiological, and behavioral data. The framework also has implications for therapeutic approaches to spinal cord and brainstem disorders: cell replacement or reprogramming strategies must consider not only cell type identity but also positional identity to achieve appropriate circuit integration, and the shared molecular taxonomy of spinal and brainstem populations suggests that genetic mutations affecting either cell type specification or regional patterning may have consequences that extend across the full neuraxis.

The spinal cord and brainstem have long been viewed as central to the execution and regulation of nervous system function, translating intention into movement, relaying sensory information to the brain, and sustaining the autonomic processes essential for life. The molecular atlas presented here suggests that these structures are organized according to a logic that is more streamlined and conserved than their functional diversity might imply, with a relatively small set of cell types arranged along conserved spatial axes and modulated by positional programs to meet the specific demands of each segment. Understanding this organizational logic, and how it is disrupted in injury and disease, is now increasingly within reach.

## METHODS

### Mouse breeding and husbandry

All experimental procedures related to the use of mice were approved by the Institutional Animal Care and Use Committee of the Allen Institute for Brain Science, in accordance with NIH guidelines. Mice were housed in rooms with temperature (21–22 °C) and humidity (40–51%) control at no more than five adult animals of the same sex per cage. Mice were provided with food and water ad libitum and were maintained on a regular 14:10 h light/dark cycle. Mice were maintained on the C57BL/6 J (RRID: IMSR_JAX:000664) background. We excluded any mice with anophthalmia or microphthalmia. All donor animals used for data generation are listed in **Supplementary Table 1**. No statistical methods were used to predetermine sample size. In total we used 16 C57Bl6/j wild type donors to collect snMultiome data from 184,657 nuclei across four spinal cord segments. To enrich motor neurons, we collected nuclei from RCL-Sun1sfGFP-neo;Chat-IRES-Cre mice and performed fluorescence-positive nuclei isolation by fluorescence-activated cell sorting (FACS). Nuclei from some donors were pooled to meet the minimum nuclei number for loading the 10x chip (**Supplementary Table 1**).

### Single-nucleus isolation

Mice were anaesthetized with 2.5–3% isoflurane and transcardially perfused with cold, pH 7.4 HEPES buffer containing 110 mM NaCl, 10 mM HEPES, 25 mM glucose, 75 mM sucrose, 7.5 mM MgCl2, and 2.5 mM KCl to remove blood from brain ^72^. Following perfusion, the brain was dissected quickly, frozen for 2 min in liquid nitrogen vapor and then moved to −80 °C for long term storage following a freezing protocol developed at AIBS ^73^.

Nuclei were isolated using the RAISINs method ^74^ with a few modifications as described in a nuclei isolation protocol developed at AIBS ^75^. In short, excised tissue dissectates were transferred to a 12-well plate containing CST extraction buffer. Mechanical dissociation was performed by chopping the dissectate using spring scissors in ice-cold CST buffer for 10 min. The entire volume of the well was then transferred to a 50-ml conical tube while passing through a 100-μm filter and the walls of the tube were washed using ST buffer. Next the suspension was gently transferred to a 15-ml conical tube and centrifuged in a swinging-bucket centrifuge for 5 min at 500 rcf and 4 °C. Following centrifugation, the majority of supernatant was discarded, pellets were resuspended in 100 μl 0.1× lysis buffer and incubated for 2 min on ice. Following addition of 1 ml wash buffer, samples were gently filtered using a 20-μm filter and centrifuged as before. After centrifugation most of the supernatant was discarded, pellets were resuspended in 10 μl chilled nuclei buffer and nuclei were counted to determine the concentration. Nuclei were diluted to a concentration targeting 5,000 nuclei per μl.

### cDNA amplification and library construction

For 10x Multiome processing, we used the Chromium Next GEM Single Cell Multiome ATAC + Gene Expression Reagent Bundle (1000283, 10x Genomics). We followed the manufacturer’s instructions for transposition, nucleus capture, barcoding, reverse transcription, cDNA amplification and library construction^76^. For the snMultiome libraries, we loaded 10,156 ± 4,208 nuclei per port across 19 libraries (**Supplementary Table 1**). For snRNA-seq we targeted a sequencing depth of 120,000 reads per nucleus. For snATAC-seq we targeted a sequencing depth of 85,000 reads per nucleus

### Sequencing data processing and QC

To remove low-quality nuclei, we developed a stringent QC process. Nuclei were first classified into broad cell classes after mapping to our established Allen Brain Cell Atlas for the whole mouse brain (ABC-WMB Atlas)^12^ and nuclei quality was assessed based on gene detection and doublet score. Doublets were identified using a modified version of the DoubletFinder algorithm (available in scrattch.hicat)^77^ and removed when doublet score was > 0.3. For 10x Multiome snATAC-seq data, we used the default criteria implemented in ArchR (RRID:SCR_020982)^78^: number of unique nuclear fragments (nFrags > 1000) and signal-to-background ratio (TSS > 3). Only nuclei having passed both snRNA-seq and snATAC-seq QC criteria (total 109,373 nuclei) were included in the downstream analysis.

### External datasets

We combined our snMultiome spinal cord dataset with 12 published datasets covering the brainstem and spinal cord (**Supplementary Table 2**). Raw counts, and if available cell type and meta data annotations, for each of the datasets were downloaded from their respective sources. We applied minimal QC thresholds to the external datasets (**Supplementary Table 2**), resulting in a total of 224,222 remaining nuclei for integration with our own dataset. The combined datasets vary greatly in number of genes and UMI’s detected (**Extended Data Fig. 2**).

### scVI integration, clustering, and label transfer

Raw counts were log-normalized and subsequently used to select 4,000 highly variable genes. For dimensionality reduction and batch correction, the raw counts of the highly variable genes were used to train the scvi-tools (v0.17.1, RRID:SCR_026673) scVI variational autoencoder (parameters: batch_key = ‘dataset’+‘platform’, n_latent = 32, gene_likelihood = ‘nb’, n_layers = 3)^79^. The latent dimensions were used to generate a KNN graph for UMAP generation and subclustering. We performed label transfer from the ABC-WMB Atlas using MapMyCells^80^. Clustering was conducted in the integrated space to generate coarse-level clusters, which were subsequently annotated based on the transferred labels. Clusters in which the majority of cells originated from brain tissue were excluded from downstream analyses. We then performed a second round of scVI integration separately for glutamatergic, GABAergic, cholinergic, motor neuron, and non-neuronal populations. To ensure comparable granularity across populations, we used the same clustering resolution parameters described above. Within each integrated class dataset, we applied iterative clustering using scrattch.bigcat (v0.0.5, https://github.com/AllenInstitute/scrattch.bigcat)^12^. Finally, we conducted a final round of cluster merging within each of the aforementioned populations to define the final cell types.

### UMAP projection

The scVI or MultiVI integrated latent spaces were used as input to create 2D and 3D UMAPs (v0.5.6, RRID:SCR_018217)^81^, using parameters nn.neighbors = 25 and md = 0.4.

### Constellation plot

To generate the constellation plot, each transcriptomic supertype was represented by a node (circle), whose surface area reflected the number of cells within the supertype in log scale. The position of nodes was based on the centroid positions of the corresponding supertypes in UMAP coordinates. The relationships between nodes were indicated by edges that were calculated as follows. For each cell, 15 nearest neighbors in reduced dimension space were determined and summarized by supertypes. For each supertype, we then calculated the fraction of nearest neighbors that were assigned to other supertypes. The edges connected two nodes in which at least one of the nodes had > 5% of nearest neighbors in the connecting node. The width of the edge at the node reflected the fraction of nearest neighbors that were assigned to the connecting node and was scaled to node size. For all nodes in the plot, we then determined the maximum fraction of “outside” neighbors and set this as edge width = 100% of node width. The function for creating these plots, plot_constellation, is included in scrattch.bigcat (v0.0.5, https://github.com/AllenInstitute/scrattch.bigcat)^12^.

### Xenium data generation

Mice were deeply anesthetized with a ketamine/xylazine solution and transcardially perfused with ice-cold sterile saline, followed by 10% neutral buffered formalin (NBF) for at least 10 min. To improve fixative penetration, the dorsal spinal column was removed, and the spinal cords (with the remaining spinal column) were incubated in NBF for 18–20 h at 4 °C on a shaker. Samples were then washed three times in 70% ethanol. After spinal cord dissection, 2 mm-log spinal cord chunks were embedded in HistoGel (Epredia) and processed for paraffin embedding using a tissue processor. Samples were dehydrated through increasing concentrations of ethanol (e.g., 80%, 95%, and multiple changes of 100%; 30–90 min per step). Tissues were then cleared in a xylene substitute (Formula 83; 3 × 1 h) and infiltrated with molten paraffin wax (3 × 1 h at ∼60°C) prior to embedding. Sections were mounted onto Xenium slides according to the manufacturer’s protocol (CG000578). Briefly, samples were exposed, hydrated in ice-cold RNAse-free water, sectioned in 39C water bath and finally placed on Xenium slides. The slides were dried at room temperature, then incubated at 42C for 3 hours. Then imaged using standard manufacturer’s protocol (CG000580, CG000760) using Xenium Prime chemistry with 5K mouse pan tissue panel and custom genes **(full panel in Supplemetary table 5)** with advanced segmentation kit. In order to maximize the capture of all the cell types, final dataset was resegmented using a custom-developed CellPose-based model based on 18S and DAPI channels of advanced segmentation stain.

### Integration of spatial transcriptomics and single-nucleus RNA-seq data

To assign cell type identities to spatially resolved cells, Xenium and MERFISH *in situ* transcriptomics data were integrated with the single-nucleus RNA-seq reference atlas using scVI^79^. Spatial transcriptomics cells and reference single-nucleus profiles were concatenated into a joint AnnData object. Genes present in the reference but absent from the Xenium gene panel were padded with zeros to produce a common gene space. A scVI model was trained on the combined dataset with dataset identity as the batch covariate, using a medium-sized architecture (128 hidden units, 2 layers, 32 latent dimensions) with gene-batch dispersion, a maximum of 2048 training epochs, and early stopping. A 32-dimensional latent representation was extracted for all cells. Two-dimensional and three-dimensional UMAP embeddings were computed from the scVI latent space using 10 nearest neighbors.

### Cell type label transfer by Random Forest classification

Cell type labels were transferred from the reference single-nucleus dataset to spatial transcriptomics cells using Random Forest classifiers trained on scVI latent representations^77^. Prior to classifier training, reference cell metadata were updated to reflect the final integrated taxonomy and cells assigned to non-spinal and non-medullary classes were excluded. Label transfer followed a two-step hierarchical strategy. First, a subclass-level classifier was applied to all query cells. Next, cluster-level classification was performed within each predicted subclass, restricting the label space to clusters within that subclass. Reference cells were downsampled per label prior to training to mitigate class imbalance (up to 1000 and 200 cells per type at the subclass and cluster levels respectively). Each Random Forest comprised 25 trees. For each cell, the predicted label and its associated probability score, defined as the maximum vote fraction across trees, were retained for downstream analysis.

### Niche analysis

Prior to niche analysis, the spatial dataset have been filtered to remove cells with low mapping probability or having mixed transcriptional signature due to off-target or low-sensitivity transcript detections. To define spatial niches in the spinal cord, we applied the scNiche^44^ tool to our Xenium spatial transcriptomics dataset. Data from different sample batches were first integrated using Harmony, and the resulting integrated latent space was used as input for the scNiche pipeline. The analysis was restricted to neuronal cells, with each tissue section treated as an individual sample (31 sections total, including both coronal and longitudinal sections). For niche construction, 30 nearest neighbors were considered for each cell. Model training was performed in 10 batches for 100 epochs. Spatial niches were then defined using Leiden clustering at a resolution of 2 on a 20-neighbor cosine kNN graph.

### Spatial domain clustering

To group Xenium neuronal niches into broader spatial domains, we compared niches based on their neuronal cell type composition. For each niche cluster, we quantified its composition across neuronal supertypes, generating a niche-by-supertype count table. To reduce noise, we retained only supertypes represented by at least 25 cells. Counts were then converted to within-niche proportions, such that each niche was represented by its relative supertype composition profile.

We next calculated pairwise similarity between niches using cosine distance on these proportional composition profiles and performed hierarchical clustering with average linkage. For visualization, the resulting dendrogram was reordered using optimal leaf ordering. Broader composition-based spatial domain groups were then defined by cutting the hierarchical tree at a distance threshold of 0.4. In this way, niches with similar neuronal supertype composition were grouped into shared spatial domains.

### Mapping of spinal cord lamina organization onto the brainstem

To investigate whether shared organizational principles exist between the spinal cord and brainstem, we extended the niche analysis to generate a shared niche framework across both regions. To do this, we first generated a shared latent space between the spatial datasets. In short, MERFISH and Xenium spatial transcriptomics datasets were integrated using KNN anchor-based alignment. Each dataset had been independently integrated with the common RNA-seq reference using scVI, but the resulting latent spaces were not directly comparable. We created a unified reference by averaging the latent representations of 114,122 shared RNA-seq cells, then mapped each spatial cell to this reference using k=50 nearest neighbors with inverse distance weighting. This approach enabled flexible integration while preserving local transcriptomic relationships.

Using this unified latent space, we reran niche analysis on the combined spinal cord and brainstem dataset and processed the resulting niches using the same procedure described above for the spinal cord. To map the newly defined niches onto previously assigned spinal cord laminae, we assigned each brainstem cell the most prevalent lamina identity among its 100 nearest neighbors in the latent niche space. Each niche was then assigned to a lamina based on the dominant lamina identity of the cells belonging to that niche.

### Spatial Axis Scoring

To characterize the anatomical distribution of each cell type, we assigned normalized spatial scores along tree axes, dorsoventral (D-V), mediolateral (M-L), and rostrocaudal (R-C), to each cell in spatially resolved transcriptomic datasets. Scores were computed within individual tissue sections, ensuring comparability across sections, spinal levels, and datasets regardless of differences in tissue size or coordinate systems. A coordinate scale check was performed for each dataset by computing the median nearest-neighbor distance across a random subsample of 2,000 cells, verifying spatial resolution consistency prior to scoring. The D-V score was computed by linearly normalizing each cell’s y-coordinate within its section. A score of 0 indicates the ventral extreme of the section and 1 indicates the dorsal extreme.

The M-L score was derived from each cell’s absolute distance from the estimated anatomical midline of the section. Rather than using the simple x-range midpoint, the midline was estimated computationally by identifying the x-coordinate that minimized the asymmetry between the left and right hemispheres when mirrored onto each other. Specifically, a kernel density estimate (Gaussian KDE, Scott’s bandwidth) was constructed from the x-coordinates of all cells in the section, and a candidate midline was selected from the inner 60th percentile of the x range. For each candidate, the right-side density was reflected onto the left, and the mean squared difference between the two mirrored density profiles was computed. The midline minimizing this asymmetry score was selected. A score of 0 indicates the medial center and 1 indicates the lateral edge.

### Analysis of spatial gene expression gradients

To identify genes with spatial expression patterns independent of cell type identity, we performed spatially variable gene (SVG) analysis on Xenium single-cell spatial transcriptomic data using Moran’s I statistic as implemented in Squidpy^82^. Prior to analysis, cells were filtered to retain those with a minimum of 5 total transcript counts, and genes were retained if detected in at least 5 cells per section, to remove low-quality observations whose near-zero expression produces undefined dispersion statistics. Expression values were taken from log-normalized. A spatial k-nearest neighbor graph (k=10) was constructed for each section independently using Euclidean distances between cell centroid coordinates (coord_type=“generic”), and Moran’s I was computed across all retained genes in parallel. Sections were analyzed independently to avoid spatial coordinate system conflicts between sections and to enable assessment of cross-section reproducibility. Genes were ranked by their Moran’s I on cell-type-regressed residuals and retained as candidate gradient genes if they ranked within the top 500 SVGs in at least 50% of sections. A cross-section mean Moran’s I was computed for each gene as a summary statistic of gradient strength.

To identify co-varying gene programs that define continuous spatial gradients across the spinal cord, we applied Non-Negative Matrix Factorization (NMF) to spatially-binned, cell-type-regressed expression data. NMF was chosen over clustering-based approaches because it produces continuous, additive decompositions suited to gradient structure: each factor represents a spatial program with a per-bin score and a gene loading vector, rather than assigning discrete cluster memberships. For each section independently, a uniform grid was defined along both spatial axes with a target of 20 cells per bin and a minimum of 10 bins per section. Within each section, cells were assigned to bins by their x- and y-coordinates, and the mean residual expression across cells in each bin was computed to yield a bin-by-gene expression matrix. Bins containing fewer than 5 cells were discarded. The resulting bin matrix was concatenated across all sections and datasets prior to NMF.

NMF was performed using scikit-learn’s implementation^83^ with n_components=15 factors, non-negative double singular value decomposition initialization (init=“nndsvda”) for deterministic and biologically-motivated initialization, a maximum of 1,000 iterations, and a small L1 regularization term (l1_ratio=0.1) on gene loadings to encourage sparser, more interpretable gene modules. Factorization yielded a bin-by-factor score matrix (W) and a factor-by-gene loading matrix (H). Each factor was interpreted as a spatial gene expression program (gradient factor), and the top genes per factor were identified by their NMF loading values in H. Bin-level factor scores and gene loadings were saved as outputs for downstream analysis and cross-dataset alignment. To enable single-cell resolution downstream analyses, NMF factor scores were computed independently for each cell from its own gene expression profile rather than by inheriting the score of its spatial bin. NMF was run independently for each Xenium batch. To identify gradient factors shared across datasets, the gene loading matrices (H) from both runs were aligned by computing pairwise cosine similarities between all factor loading vectors, restricted to genes present in both datasets. Factor pairs with cosine similarity ≥ 0.5 were considered matched across datasets, indicating a reproducible spatial gene expression program.

### Multiome peak calling

To call chromatin accessibility peaks in the snATAC-seq data, we first categorized snMultiome data according to supertype label and roi. We kept only the group category with more than 50 cells. We generated pseudo-bulk replicates using the ArchR function addGroupCoverages. We created a reproducible merged peak set using function addReproduciblePeakSet. Finally, we built the peak by cell matrix, which contains insertion counts within the merged peak set using function “addPeakMatrix”.

### Identification of differentially expression genes and differentially accessible peaks

To characterize transcriptomic and chromatin accessibility variation across cell populations, we analyzed paired gene expression and ATAC-seq data profiled by snMultiome from following rois: Thor, Cerv, Sac, Lum, and CB/MY. Gene expression data were log-CPM normalized prior to analysis, while chromatin accessibility data were retained as raw counts. To mitigate the influence of highly abundant cell clusters on differential analysis, we applied cluster-level downsampling capping each cluster at 500 cells, yielding a final dataset of 48,500 cells with 17,274 genes and 847,384 peaks with paired modalities. Differential expression (DE) analysis was performed using a fast limma-based model (fast_limma), while differential accessibility (DA) analysis was conducted using a chi-squared test due to the binary distributional properties of accessibility data. Both DE and DA analyses were carried out using the de_all_pairs and de_selected_pairs functions from the scrattch.bigcat package. Pairwise comparisons were performed across subclass, supertype, cluster, and subclass-by-ROI(subclass across different regions of interest). This multi-resolution design enables detection of both broad transcriptomic and epigenomic differences between major cell classes and finer-grained variation at the region-specific level between spinal cord and brain.

### MultiVI integration

We performed integrative analysis of multiomic and transcriptomic datasets using the MultiVI model implemented in scvi-tools (version 1.1.2). snMultiome data, for which both gene expression and chromatin accessibility modalities were available (N = 154,893), were integrated with RNA-seq–only cells (N = 224,996), combining multiple previously published transcriptomic datasets described above. These two “datasets” were jointly modeled to learn a shared latent representation across modalities and data sources (parameters: batch_key = ‘dataset’+‘platform_recode’+ ‘load_name’, n_hidden = 283, n_latent = 16, n_layers_encoder = 2, n_layers_decoder = 2).

As input to MultiVI, we constructed a cell-by-feature matrix including both gene expression features (N = 4,449 differentially expressed (DE) genes) and chromatin accessibility peaks (N = 80,439 differential accessible (DA) peaks). Features were selected based on DE genes and DA peaks identified from pairwise comparisons across cell types at multiple annotation levels (class, subclass, supertype, and cluster), as described above. Sex-linked genes (Xist, Tsix, Ddx3y, Eif2s3y, Uty) were removed prior to integration.

Given the pronounced transcriptional and epigenomic differences between neuronal and non-neuronal populations, we performed separate MultiVI integrations for neuronal and non-neuronal cells. To reduce computational burden, a representative subset of cells from each group was used for model training, and the trained model was subsequently used to project the full dataset into the learned latent space.

After training, joint, expression-specific, and accessibility-specific latent representations were extracted from the model and used to generate UMAP visualizations. Unless otherwise specified, downstream analyses and visualizations were based on joint latent space. MultiVI was further used to infer imputed chromatin accessibility values. To validate imputed chromatin accessibility, we compared measured and MultiVI-imputed accessibility profiles across all DA peaks (N = 80,439) for paired multiome cells. Measured ATAC-seq counts were normalized to count per 1e4 (10000), then log1p-transformed for comparison. For each condition (measured and imputed), a subclass-by-peak mean accessibility matrix was constructed by averaging values across cells within each subclass. Both matrices were then Z-score normalized per peak across subclasses.

### Cell type classification using Random Forest on MultiVI latent representations

To evaluate the quality of the MultiVI latent representations, we trained Random Forest classifiers (n_estimators = 100) to predict subclass labels and subclass–region combinations from each of the three latent spaces independently: accessibility, expression, and joint. Classification was performed separately for neuronal and non-neuronal cells. We used a held-out evaluation strategy in which training cells (those used during MultiVI model training) were separated from test cells (all remaining projected cells), ensuring the classifier was always tested on cells the model had not seen during training. For the accessibility-specific latent space, only paired multiome cells were included in both training and test sets, as expression-only cells lack chromatin accessibility measurements. Cell type labels with fewer than 50 cells in either the training or test partition were excluded from that evaluation. Confusion matrices were computed on the test set and row-normalized to represent the proportion of true instances of a given subclass predicted as each subclass. Classifications were performed additionally at subclass–region combinations (subclass × broad region of origin: brain versus spinal cord), to assess whether the latent representations resolve region-specific cell type identities.

### Identification of Co-accessible Peak Modules

Differentially accessible peaks were selected as candidates for module discovery (|log fold-change| > 2, rank < 500, –log_10_(p) > 5 across subclass–region comparisons). Cluster-level mean ATAC accessibility profiles were computed for each subclass–brain-region combination (normalized to counts per 30,000) and used as input for unsupervised peak clustering. Each peak’s accessibility profile was scaled to its per-peak maximum prior to graph construction. An iterative peak module algorithm was applied: a KNN graph (k = 20, Annoy cosine distance) was built over peak profiles, and Jaccard-weighted community detection was used to define initial modules. Modules above a size threshold (200 peaks) were recursively split by repeating the procedure on the submatrix. After splitting, adjacent modules were merged if their cosine similarity exceeded 0.55 and their maximum pairwise accessibility difference was below 0.45, or if either module contained fewer than 30 peaks. Module quality was assessed by mean within-module kME (eigengene correlation) and mean number of cell types with above-threshold accessibility.

Modules were retained if they contained more than 30 peaks, had a mean kME greater than 0.6, and were selectively accessible across fewer than 20 cell types on average. Within each module, peaks were ranked by kME and the top 500 by rank were used for visualization and downstream analyses. Module membership (kME) for each peak was computed as the Pearson correlation between the peak’s accessibility profile and the corresponding module eigengene (mean accessibility across module members).

For each peak in a module, candidate linked genes were identified from the peak–gene distance map (within 250 kb of the TSS). Cluster-level mean ATAC and RNA profiles were each min-max scaled per feature (using the 10th percentile of row maxima as the normalization floor). Pearson correlation between each peak’s accessibility profile and each candidate gene’s expression profile was computed across cell types. Peak–gene pairs with correlation above 0.5 were classified as linked. For each linked pair, the RNA expression profile was additionally correlated against all module eigengenes; a pair was designated as module-concordant if the gene’s best-matching module was the peak’s own module, or if the peak’s module correlation was within 0.05 of the best. The best-correlated gene for each peak (maximizing the product of kME and peak–gene correlation) was used to label peaks for visualization.

For each peak in a module, candidate linked genes were identified from the peak–gene distance map (within 250 kb of the TSS). Cluster-level mean ATAC and RNA profiles were each min-max scaled per feature (using the 10th percentile of row maxima as the normalization floor). Pearson correlation between each peak’s accessibility profile and each candidate gene’s expression profile was computed across cell types. Peak–gene pairs with correlation above 0.5 were classified as linked. For each linked pair, the RNA expression profile was additionally correlated against all module eigengenes; a pair was designated as module-concordant if the gene’s best-matching module was the peak’s own module, or if the peak’s module correlation was within 0.05 of the best. The best-correlated gene for each peak (maximizing the product of kME and peak–gene correlation) was used to label peaks for visualization.

### Anterograde tracing and immunohistochemistry

For anterograde tracing experiments from sensorimotor cortex, 8-week-old C57BL/6 mice were used. AAV2/8-CAG-hSyn-ChR2-mCherry was injected into the sensorimotor cortex at five anterior–posterior coordinates: A/P −1, −0.5, 0, 0.5, and 1 mm; M/L 1 mm; D/V −0.7 mm from the dura. Virus was injected at 5 × 10¹² vg/ml, with 200 nl delivered per site. Animals were collected 4 weeks after virus injection.

Tissue was post-fixed in 4% PFA overnight at 4°C. Brain and spinal cord tissues were dissected and dehydrated in 30% sucrose in PBS at 4°C for 2 days. Tissues were then embedded, frozen, and sectioned at 50 µm on a cryostat as free-floating sections while preserving section order. For staining, free-floating sections were rinsed three times in 1% Triton X-100 in PBS for 10 min each, then incubated in blocking solution containing 10% normal donkey serum and 0.5% Triton X-100 in PBS for 1 h at room temperature. Sections were then incubated overnight at 4°C in primary antibody solution containing 5% normal donkey serum and 0.5% Triton X-100 in PBS. The following day, sections were rinsed three times in 0.5% Triton X-100 in PBS for 10 min each, then incubated with secondary antibodies and Alexa Fluor 647-conjugated isolectin Griffonia simplicifolia IB4 diluted in PBS for 2 h at room temperature. Secondary antibody solutions were filtered through a 0.22 µm syringe filter before use. Sections were then washed three times in PBS and mounted with Fluoromount-G mounting medium for imaging. The following primary antibodies were used: rat anti-NeuN, 1:1000, Abcam, #ab279297; rabbit anti-RFP, 1:1000, Abcam, #ab34771; and rabbit anti-CGRP, 1:1000, Abcam, #ab81887. Alexa Fluor 647-conjugated isolectin Griffonia simplicifolia IB4 was used at 1:1000, Invitrogen, #I32450. The following secondary antibodies were used at 1:1000 dilution: Alexa Fluor 405-conjugated donkey anti-rat, Jackson ImmunoResearch, #712-475-150; Alexa Fluor 488-conjugated donkey anti-rabbit, Jackson ImmunoResearch, #711-545-152; and Alexa Fluor 555-conjugated donkey anti-rabbit, Jackson ImmunoResearch, #711-565-152.

## RESOURCE AVAILABILITY

### Additional Information

Correspondence and requests for further information and resources should be addressed to: C.T.J.vV.

### Data Availability

The newly generated AIBS spinal cord single nucleus Multiome data is available at NeMO (https://assets.nemoarchive.org/col-nnaxqv4). The newly generated spinal cord Xenium data are available in the Gene Expression Omnibus (GEO) under accession number GSE332614.

The scRNA-seq and MERFISH datasets for this study are part of the Allen whole mouse brain (WMB) cell type atlas and are accessible through Neuroscience Multi-omic Data Archive (NeMO, RRID:SCR_016152) (https://assets.nemoarchive.org/dat-zqurqvh) and Brain Image Library (BIL, RRID:SCR_017272) (https://doi.org/10.35077/g.610).

The following published datasets were used; Yao et al., 2023 ^12^ (https://assets.nemoarchive.org/dat-qg7n1b0, GSE246717); Langlieb et al., 2023^13^ (https://assets.nemoarchive.org/dat-aa0jwmj); Blum et al., 2021^17^ (GSE161621); Alkaslasi et al., 2021^18^ (GSE167597); Russ et al., 2021^9^ (GSE158380); Skinnider et al., 2024^16^ (GSE234774; GSE230765).

### Code Availability

The R package scrattch.bigcat, is available via github https://github.com/AllenInstitute/scrattch.bigcat. Notebooks with examples of data analysis code used in this manuscript are available via github https://alleninstitute.github.io/scrattch.example/.

## ACKNOWLEDGEMENTS

We are grateful to the Transgenic Colony Management, Lab Animal Services, Molecular Biology and Histology teams at the Allen Institute for technical support. We thank Jayaram Chandrashekar, at the Allen Institute for Neural Dynamics generously sharing unpublished medullary retrograde sequencing data that contributed to the connectivity analyses presented here. The research was funded by the U19MH114830 grant from National Institute of Mental Health to H.Z., under the BRAIN Initiative of National Institutes of Health (NIH). NIH grant 1R01NS142815 and also grants from Dr. Miriam and Sheldon G. Adelson Medical Research Foundation and Wings for Life Research Foundation to Z.H. The content is solely the responsibility of the authors and does not necessarily represent the official views of NIH and its subsidiary institutes. S.Y. was supported by the SCIRTS Postdoctoral Fellowship from the Craig H. Neilsen Foundation. This work was also supported by Harvard Medical School Rodent Histopathology core and Boston Children’s Hospital Viral Core, which is supported by NIH5P30EY012196 grant. This work was also supported by the Allen Institute for Brain Science. The authors thank the Allen Institute founders, Paul G. Allen and Jody Allen, for their vision, encouragement, and support.

## AUTHOR CONTRIBUTIONS

Conceptualization: H.Z., Z.Y., C.T.J.v.V.

Data analysis lead and coordination: Z.Y., C.T.J.v.V.

Data generation scRNA-seq: T.C., M.C., K.A.F, J.Gl., J.Gu., C.R.H, W.H., M.J., R.MC., N.P., E.P., N.S., E.D.T., A.T., N.D., K.A.S., H.Z., Z.Y., C.T.J.v.V.

Data processing and analysis (scRNA-seq): Y.G., J.X., C.L., M.T.S., N.J.J., J.Go., M.H., T.E.B., K.A.S., H.Z., Z.Y., C.T.J.v.V.

Data generation (spatial transcriptomics): E.K., K.J., Y.K., S.A.M., A.R., R. Y., S. Y., Z.H., H.Z.

Data processing and analysis (spatial transcriptomics): E.K., K.J., D.S., H.Z., Z.Y., C.T.J.v.V.

Management and supervision: N.D., T.E.B., K.A.S., Z.H., H.Z., Z.Y., C.T.J.v.V.

Manuscript writing and figure generation: E.K., Z.H., H.Z., Z.Y., C.T.J.v.V.

Manuscript review and editing: Y.G., E.K., J.X., Z.H., H.Z., Z.Y., C.T.J.v.V.

## DECLARATION OF GENERATIVE AI AND AI-ASSISTED TECHNOLOGIES

During the preparation of this work, the authors used Claude (Anthropic) in order to assist with the debugging of code, and ChatGPT (OpenAI) in order to improve the clarity and readability of the written text. After using these tools or services, the authors reviewed and edited the content as needed and take full responsibility for the content of the publication.

---

**Supplementary Figure 1.**
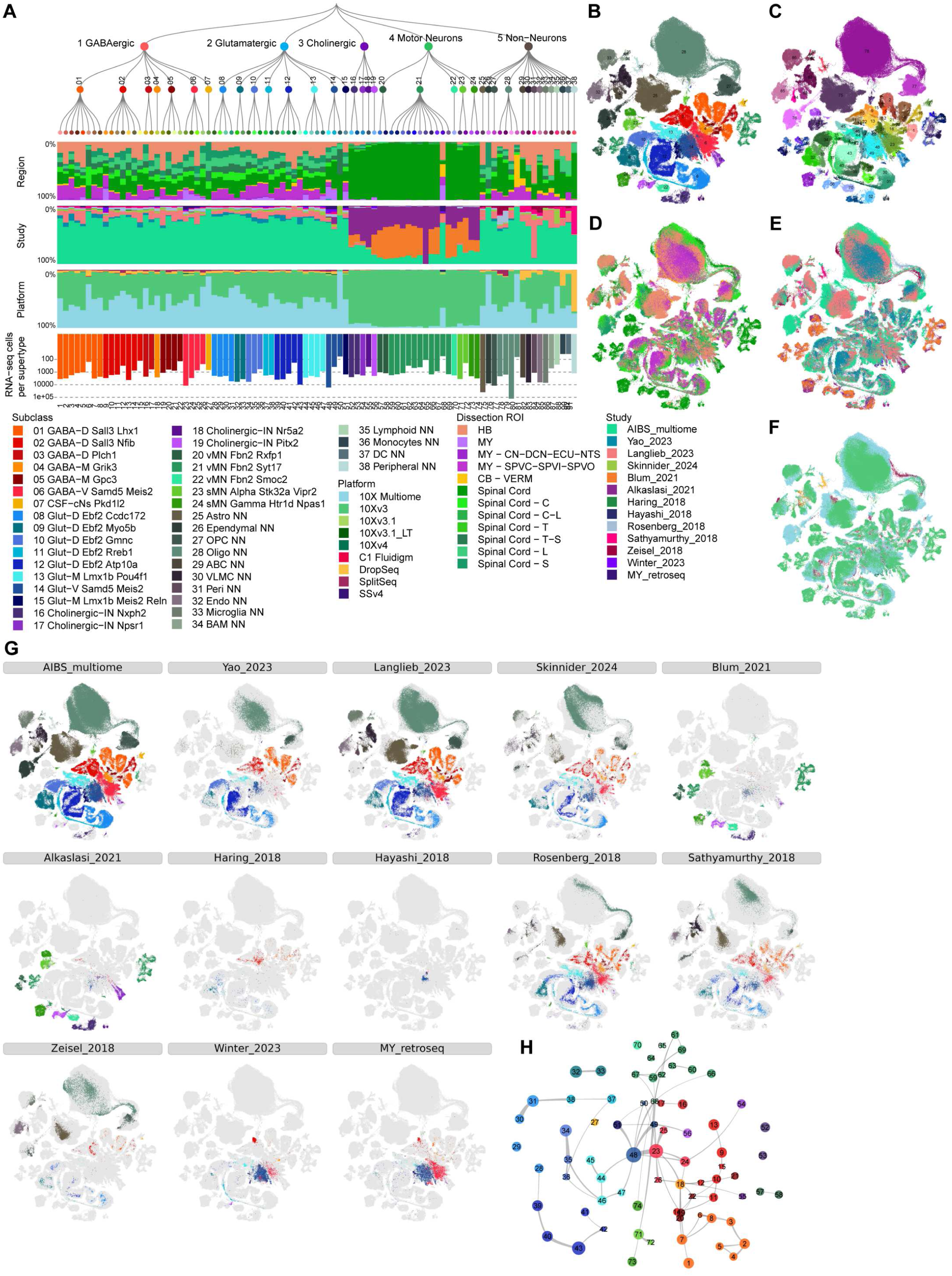
Cross-dataset integration and composition of the spinal cord neuronal atlas. **(A)** Hierarchical taxonomy of neuronal and non-neuronal supertypes shown alongside stacked bar plots summarizing atlas composition. From top to bottom: proportional contribution of CNS region (brain vs. spinal cord), contributing study, sequencing platform, and absolute number of single-nucleus RNA sequencing cells per supertype. Supertypes are ordered along the x-axis according to the dendrogram. **(B)** UMAP embeddings of all atlas cells faceted by contributing dataset, including AIBS_multiome, Yao_2023^12^, Langlieb_2023^13^, Skinnider_2024^16^, Blum_2021^17^, Alkaslasi_2021^18^, Haring_2018^8^, Hayashi_2018^19^, Rosenberg_2018^20^, Sathyamurthy_2018^21^, Zeisel_2018^11^, Winter_2023^15^, and MY_retroseq (unpublished). Each panel highlights cells from the indicated study (colored) against the full atlas in gray. **(C)** Constellation plot of the global relatedness between supertypes. Each supertype is represented by a disk, labeled by the supertype ID, and positioned at the supertype centroid in UMAP coordinates (Figure 1D). The size of the disk corresponds to the number of cells within each supertype, and the edge weights correspond to the fraction of shared neighbors (see **Methods**) between supertypes. Bubbles drawn around supertypes outline the subclass.

**Supplementary Figure 2.**
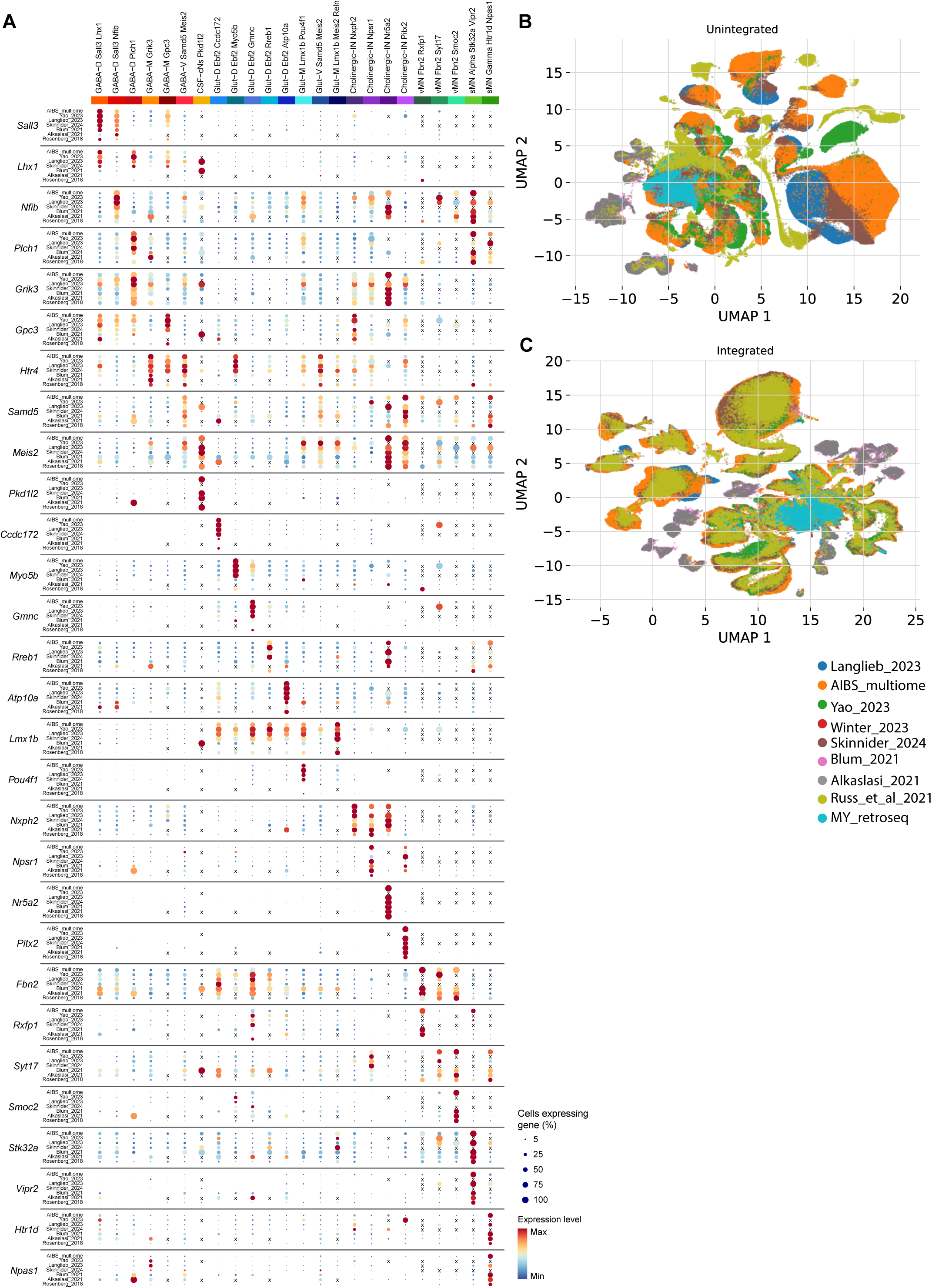
Marker gene expression across supertypes and cross-dataset integration of brain stem and spinal cord. **(A)** Dot plot showing the expression of curated marker genes (rows) across transcriptomically defined supertypes (columns) split across the different studies of origin (rows). Supertype identities are indicated along the top, grouped by major cell class (color-coded bar). Dot size reflects the percentage of cells within a cluster expressing the gene (5–100%), and dot color indicates the mean expression level (blue = minimum, red = maximum). **(B)** UMAP embedding of cells prior to dataset integration, colored by dataset of origin. Distinct clustering of cells by dataset (batch separation) indicates the presence of dataset-specific technical variation. **(C)** UMAP embedding of the same cells following cross-dataset integration, colored by dataset of origin. Improved intermingling of cells from different datasets reflects successful batch correction and alignment of shared transcriptomic structure across datasets.

**Supplementary Figure 3.**
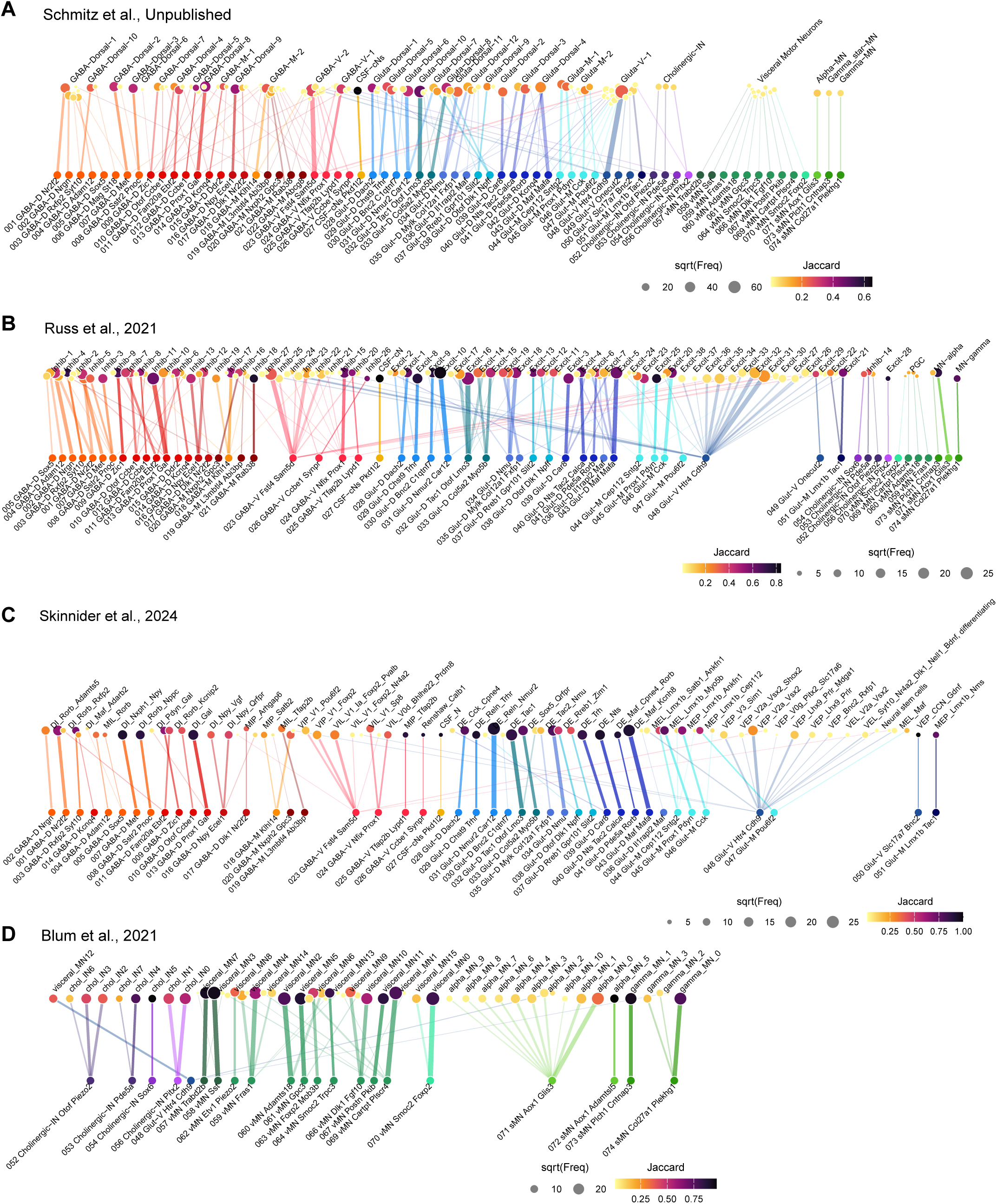
Cross-dataset supertype mapping of spinal cord datasets. (A-D) Alluvial-style correspondence plots comparing neuronal supertypes defined in the present atlas (bottom nodes) to cell type annotations from four independent published or unpublished datasets: (A) Schmitz et al., 2026^10^, (B) Russ et al., 2021^9^, (C) Skinnider et al., 2024^16^, and (D) Blum et al., 2021^17^ (top nodes). Each node represents a cell type, with node size scaled proportionally to the square root of cell frequency (sqrt(Freq)). Edges connect corresponding cell types across datasets, with edge color reflecting the Jaccard similarity index between the two matched populations.

**Supplementary Figure 4.**
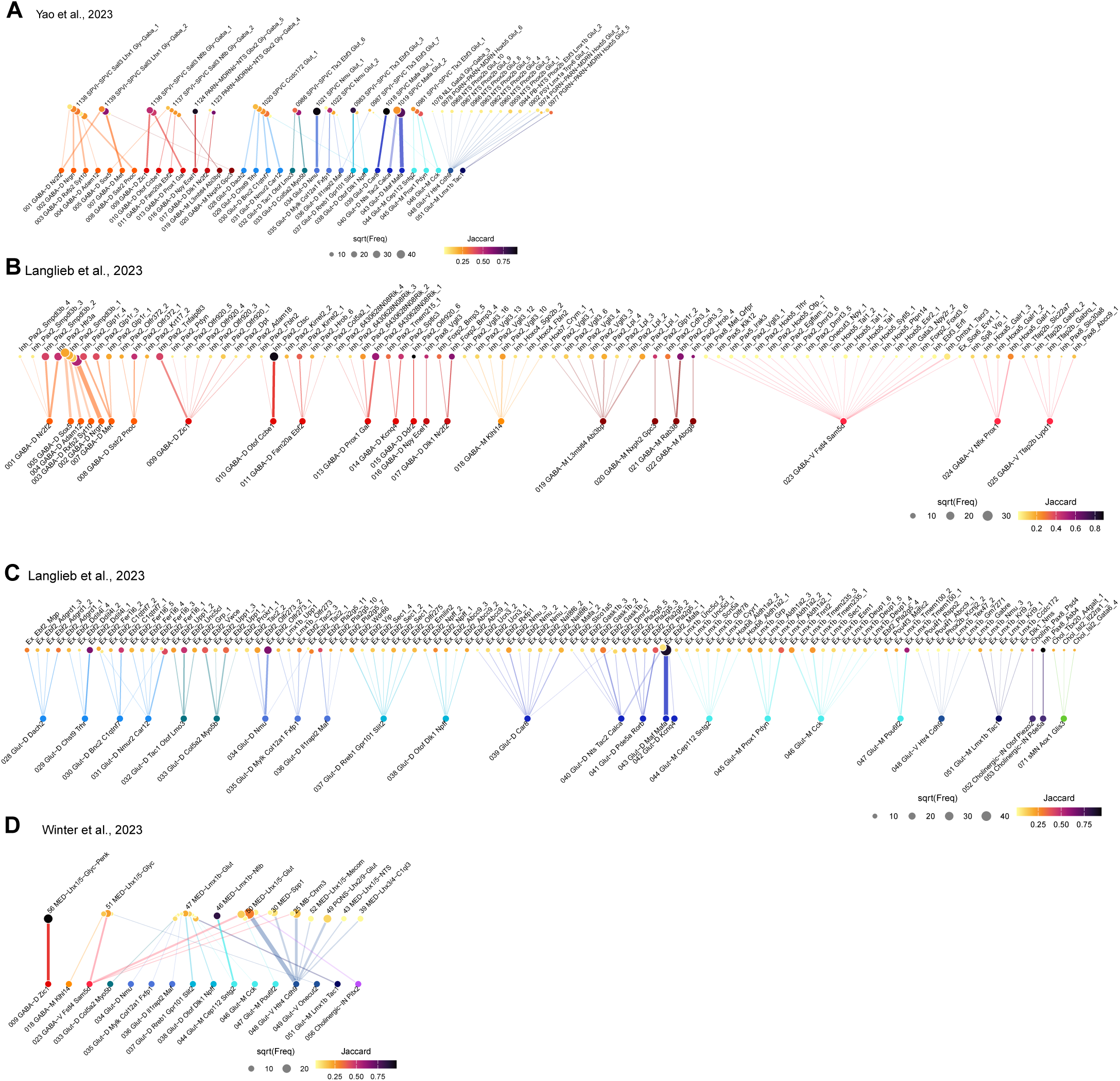
Cross-dataset supertype mapping brain datasets. (A–D) Alluvial-style correspondence plots comparing neuronal supertypes defined in the present atlas (bottom nodes) to cell type annotations from four additional datasets: (A) Yao et al., 2023^12^, (B-C) Langlieb et al., 2023^13^ (shown in two panels reflecting distinct subsets of the dataset), and (D) Winter et al., 2023^15^ (top nodes). Each node represents a cell type, with node size scaled proportionally to the square root of cell frequency (sqrt(Freq)). Edges connect corresponding cell types across datasets, with edge color reflecting the Jaccard similarity index between matched populations.

**Supplementary Figure 5.**
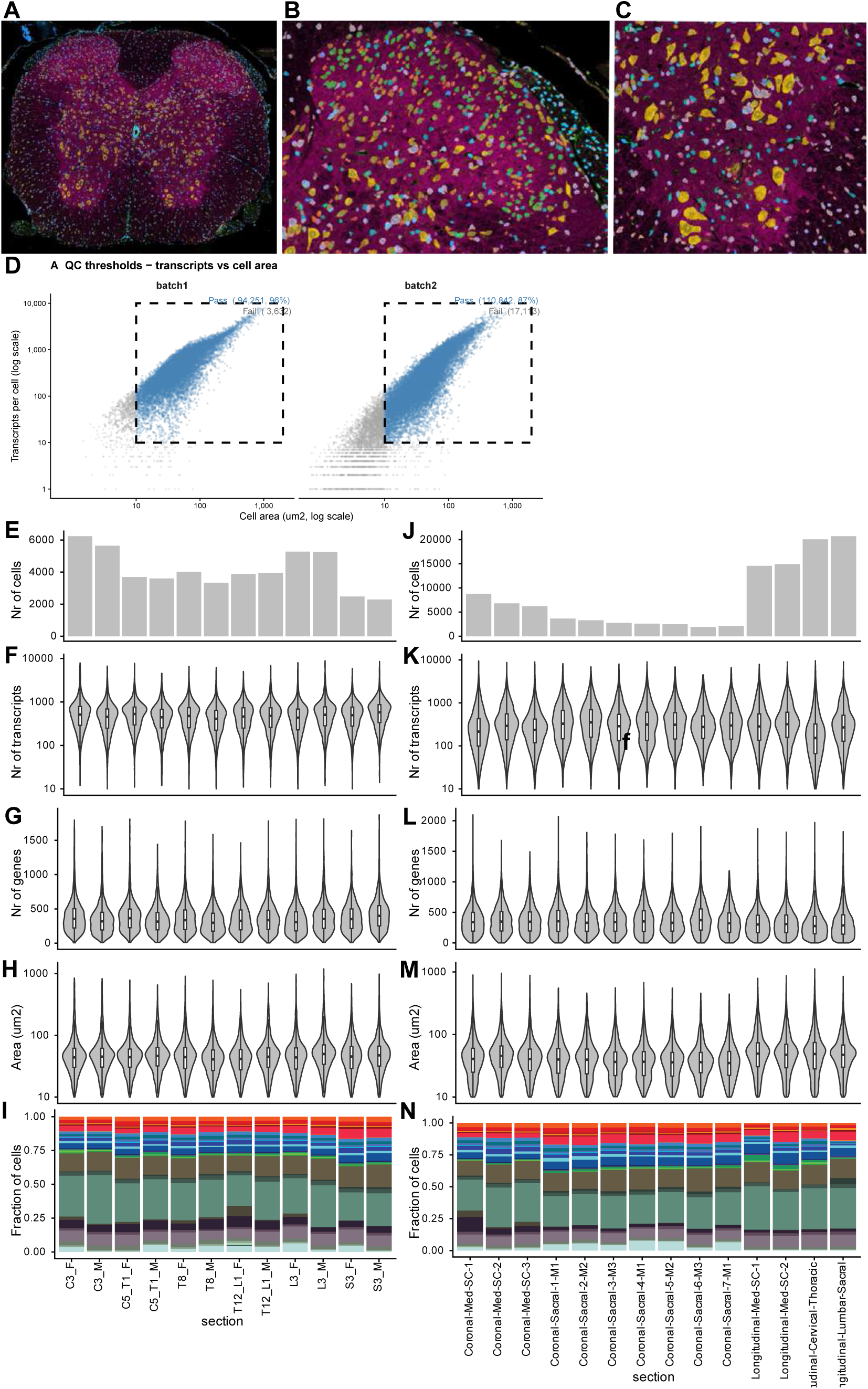
Quality control metrics for Xenium spatial transcriptomic data. (A–C) Violin plots summarizing per-cell quality control metrics across individual tissue sections and for the total dataset. From top to bottom, the three rows display the distribution of: number of transcripts per cell (Nr of transcripts) (A), number of genes detected per cell (Nr of genes) (B), and cell area (µm²) (C). Whiskers and embedded box plots indicate median and interquartile range. **(D)** Representative fluorescence images from Xenium spatial transcriptomic profiling of spinal cord tissue. The low-magnification overview (bottom left) shows the full cross-section with spatially resolved cells colored by segmentation mask. High-magnification insets (top and bottom right) illustrate the cellular resolution of the Xenium platform, with individual cells pseudo-colored by cell type identity, demonstrating the dense and spatially organized cellular architecture captured by the assay.

**Supplementary Figure 6.**
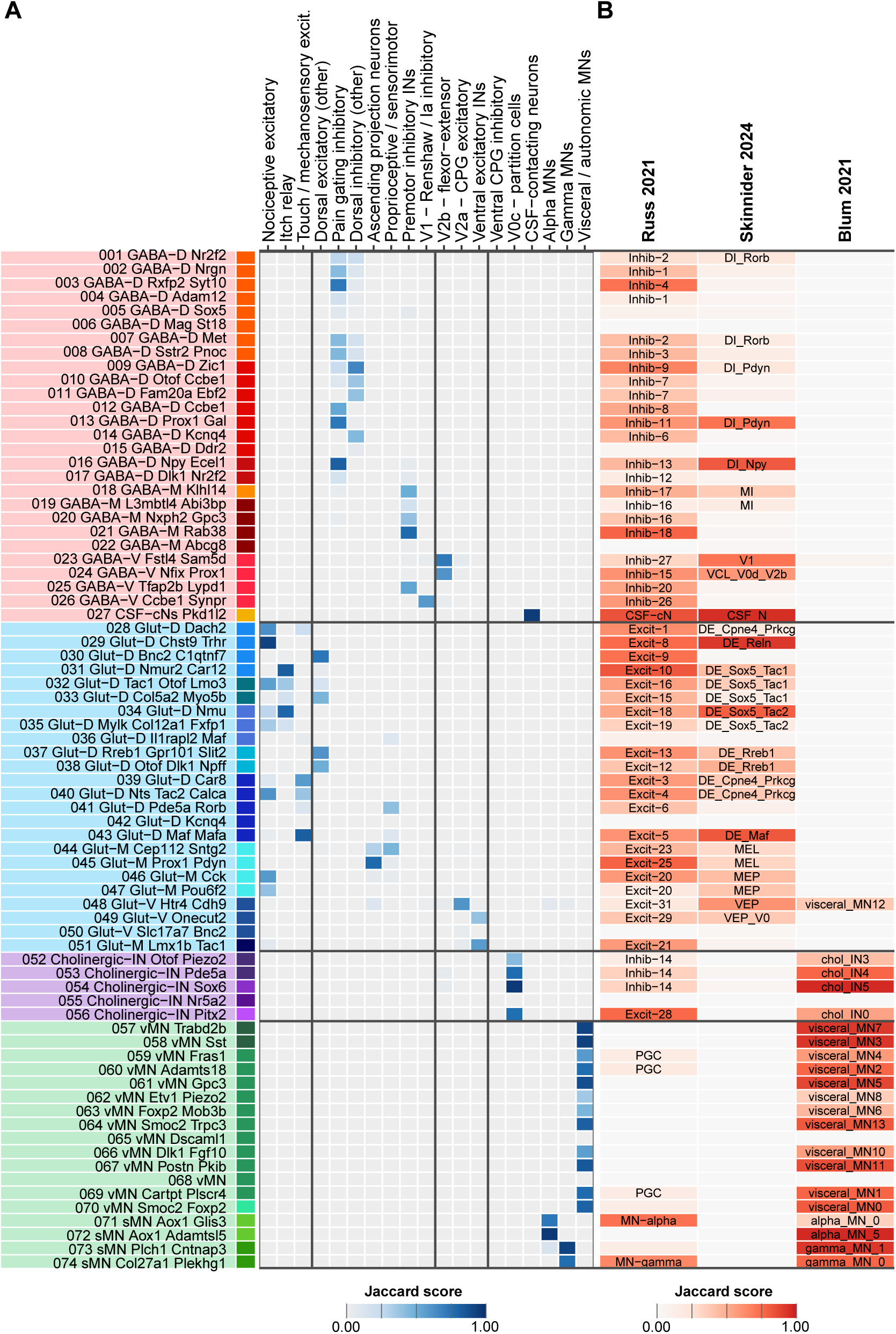
Functional annotation and cross-dataset correspondence of neuronal supertypes. **(A)** Functional group assignments for all neuronal supertypes. Each row represents a supertype and each column a functional category derived from correspondence with previously characterized populations and marker gene identity (see **Supplementary Table 4**). Colored tiles indicate assignment of a supertype to the indicated functional group. Functional categories are organized from dorsal sensory (left) to ventral motor (right), reflecting the dorsoventral organization of the spinal cord. Supertypes are ordered by class and subclass as in Figure 1. **(B)** Jaccard similarity scores between supertypes defined in this study and neuronal clusters from Russ et al. 2021^9^ (left), Skinnider et al. 2024^16^ (middle), and Blum et al. 2021^17^ (right). Each cell represents the Jaccard index between the set of cells assigned to a supertype in this atlas and a cluster in the reference dataset, computed on the basis of shared cell identities following label transfer. Higher scores indicate greater overlap between matched populations. Only the best-matching reference cluster per supertype is shown. Color scale ranges from 0 (no overlap) to 1 (complete overlap). This analysis demonstrates that the majority of supertypes defined here correspond to previously characterized populations across independent datasets, validating the taxonomy and enabling functional annotation through cross-dataset correspondence.

**Supplementary Figure 7.**
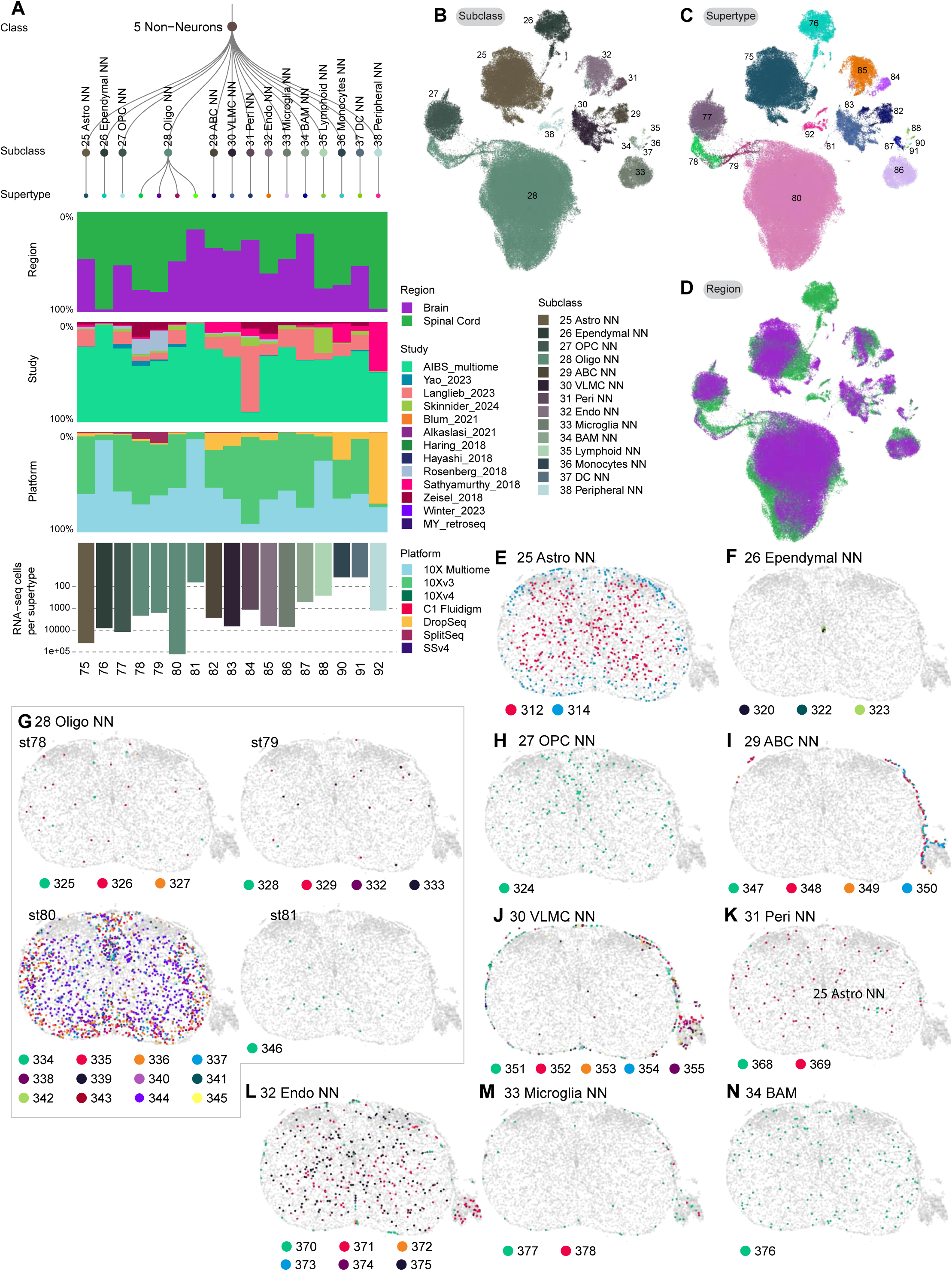
Non-neuronal cell types across the spinal cord. **(A)** UMAP projection of non-neuronal cells colored by subclass identity (subclasses 25–38). Clusters correspond to major glial, vascular, and immune cell classes, including Astrocytes (25), Ependymal (26), OPCs (27), Oligodendrocytes (28), ABCs (29), VLMCs (30), Pericytes (31), Endothelial cells (32), Microglia (33), BAMs (34), Lymphoid (35), Monocytes (36), Dendritic cells (37), and Peripheral glia (38). Supertype ID’s are annotated directly on the embedding. **(B)** The same UMAP as in (A), with cells colored by refined supertype identity (supertypes 075–092). **(C)** The same UMAP colored by tissue of origin, distinguishing cells derived from brain (purple) versus spinal cord (green). The overlapping distribution of brain and spinal cord cells across clusters indicates shared transcriptomic identity of non-neuronal cell types across CNS regions.

**Supplementary Figure 8.**
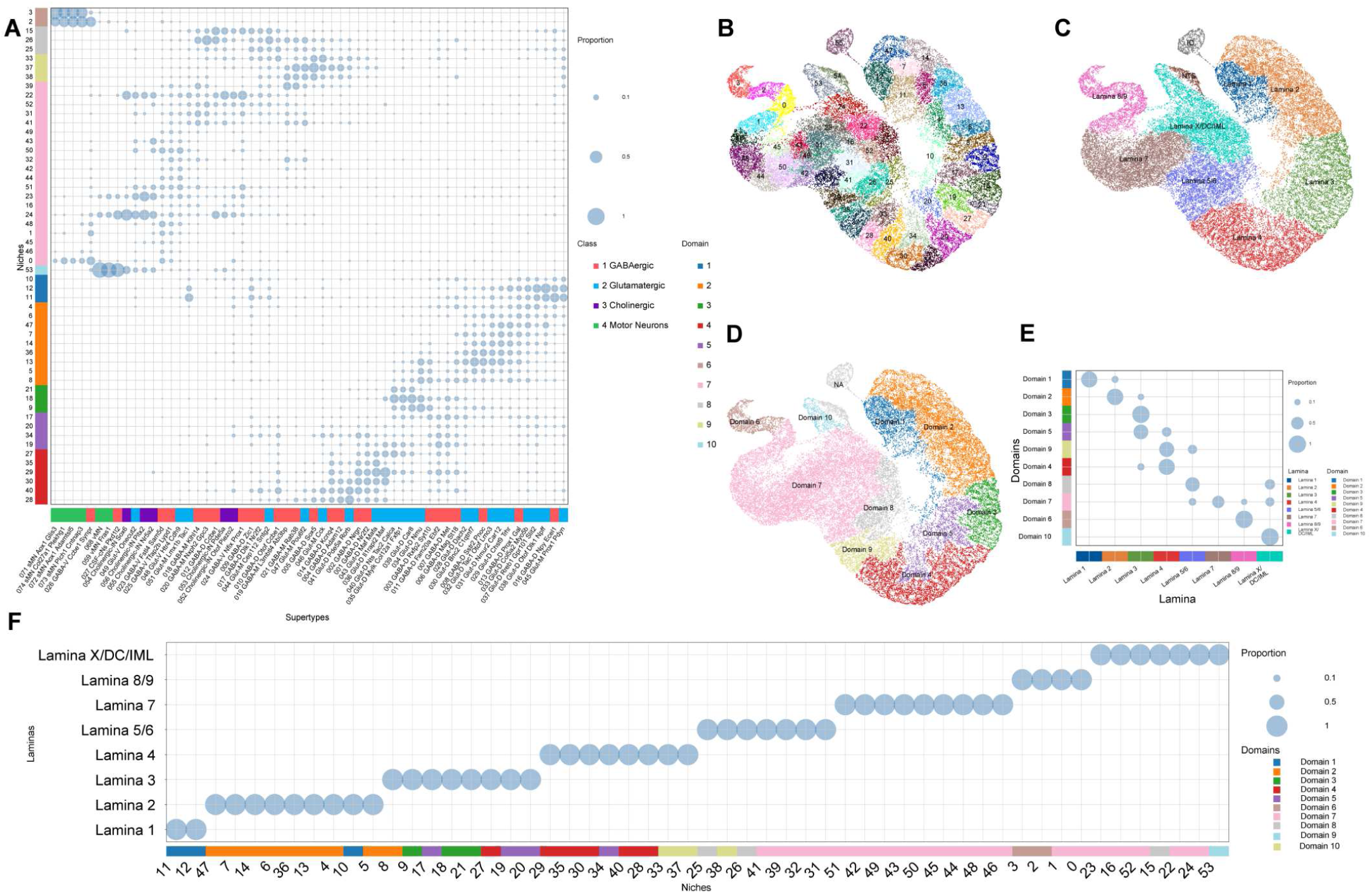
Niche composition across supertypes, spatial domains, and laminae. **(A)** Bubble plot showing distribution of supertypes across niches. The bubble size indicate proportion of cells in each supertype assigned to each niche. **(B-D)** UMAP based on the niche analysis latent space colored by (B) niche id, (C) lamina assignment, (D) niche domain. **(E)** Bubble plot comparing automatically derived niche domains based on the cell type composition and assigned laminae. Bubble size indicates the proportion of cells from each domain assigned to a given lamina. **(F)** Bubble plot indicating niche to lamina assignments.

**Supplementary Figure 9.**
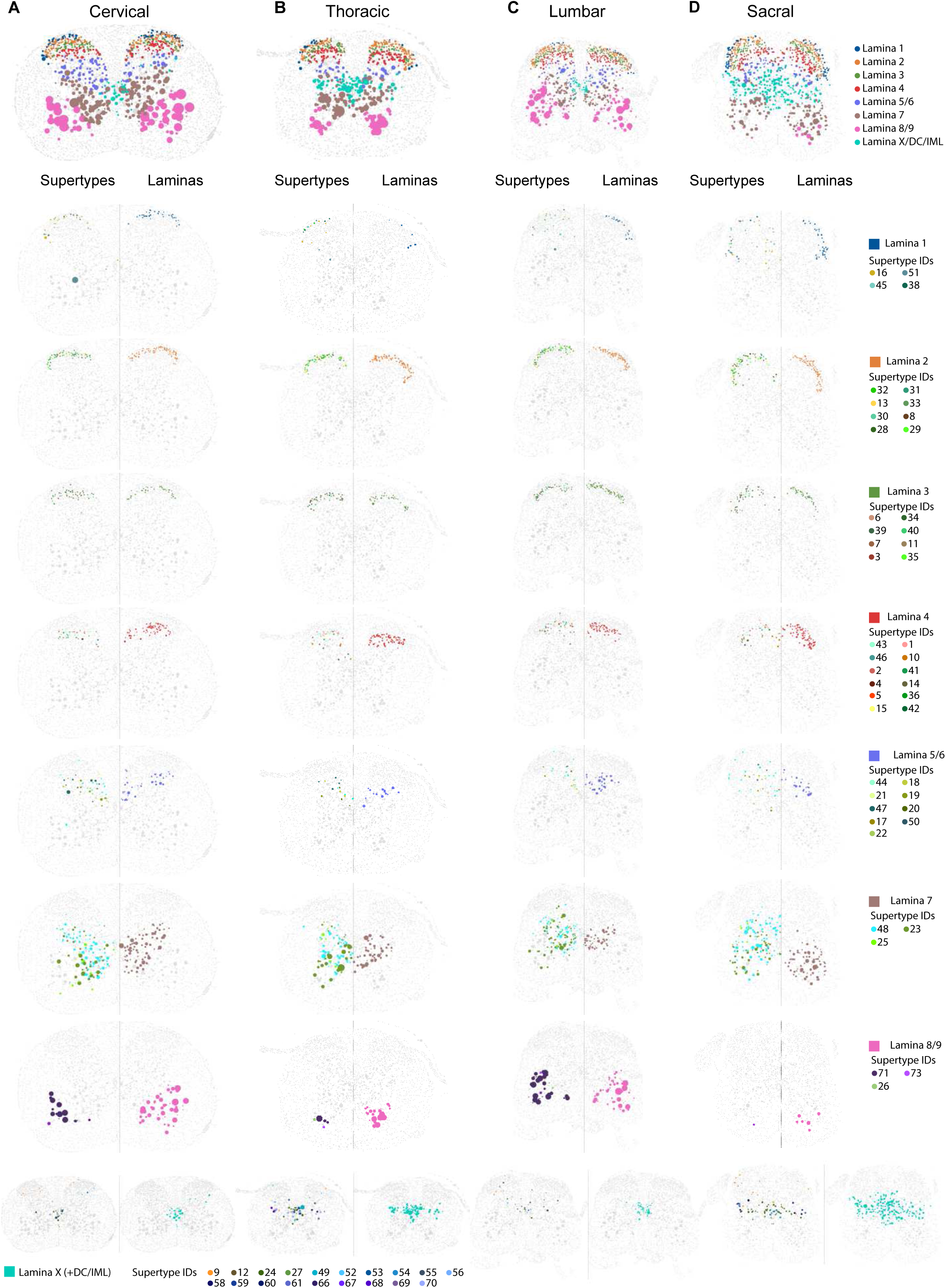
Spatial distribution of laminae and defining supertypes in spinal cord. Representative sections demonstrating laminae and supertypes defining these laminae. Each column represents **(A)** Cervical, **(B)** thoracic, **(C)** lumbar and **(D)** sacral spinal cord, each row corresponds to individual lamina. Each cell is represented by a dot with a size correlated to the cell area.

**Supplementary Figure 10.**
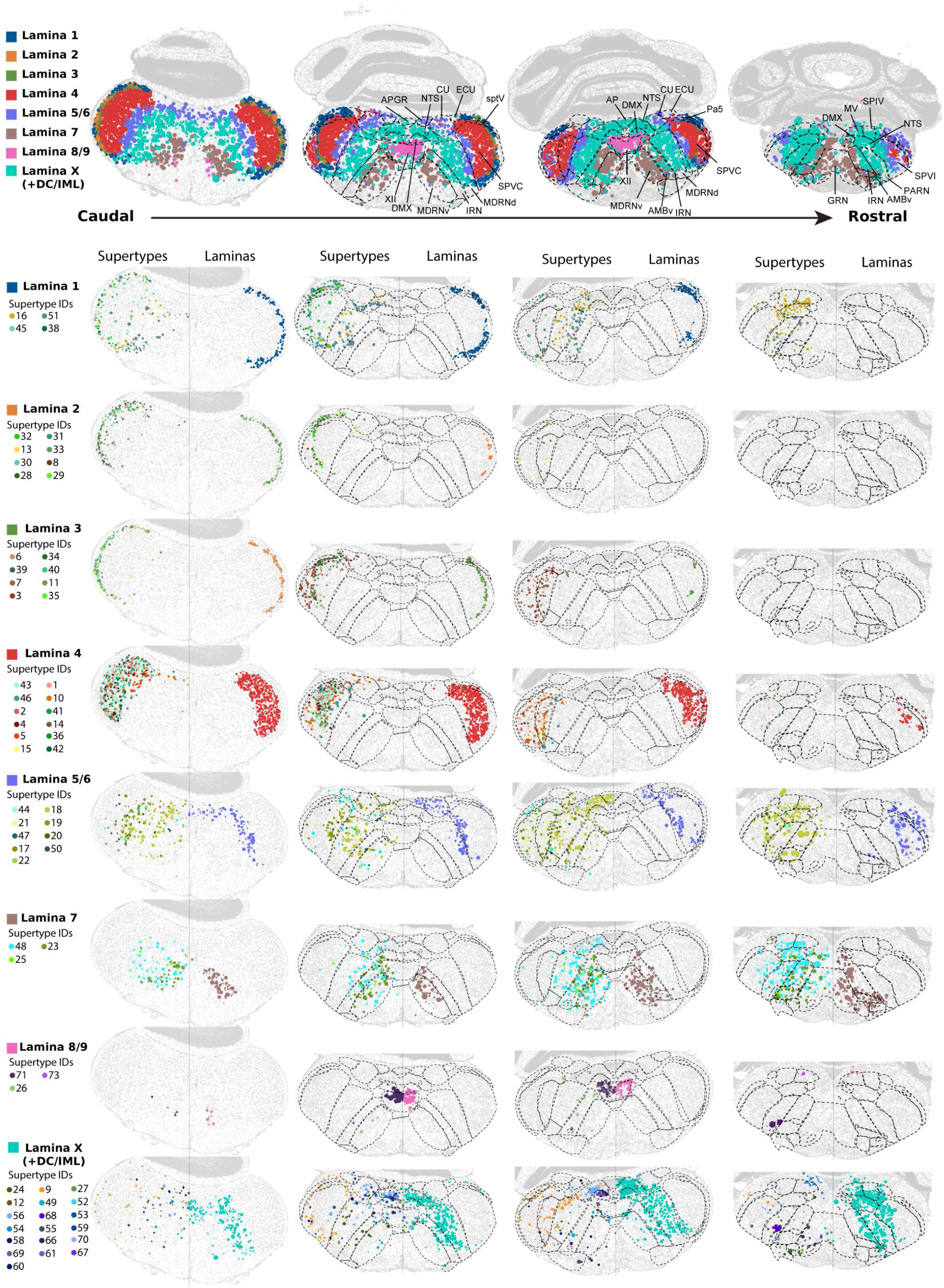
Laminar organization of neuronal supertypes across the brainstem. Representative sections demonstrating inferred laminae and supertypes defining these laminae. Each column represents a section of the brainstem from caudal to rostral, each row corresponds to individual lamina. Each cell is represented by a dot with a size correlated to the cell area.

**Supplementary Figure 11.**
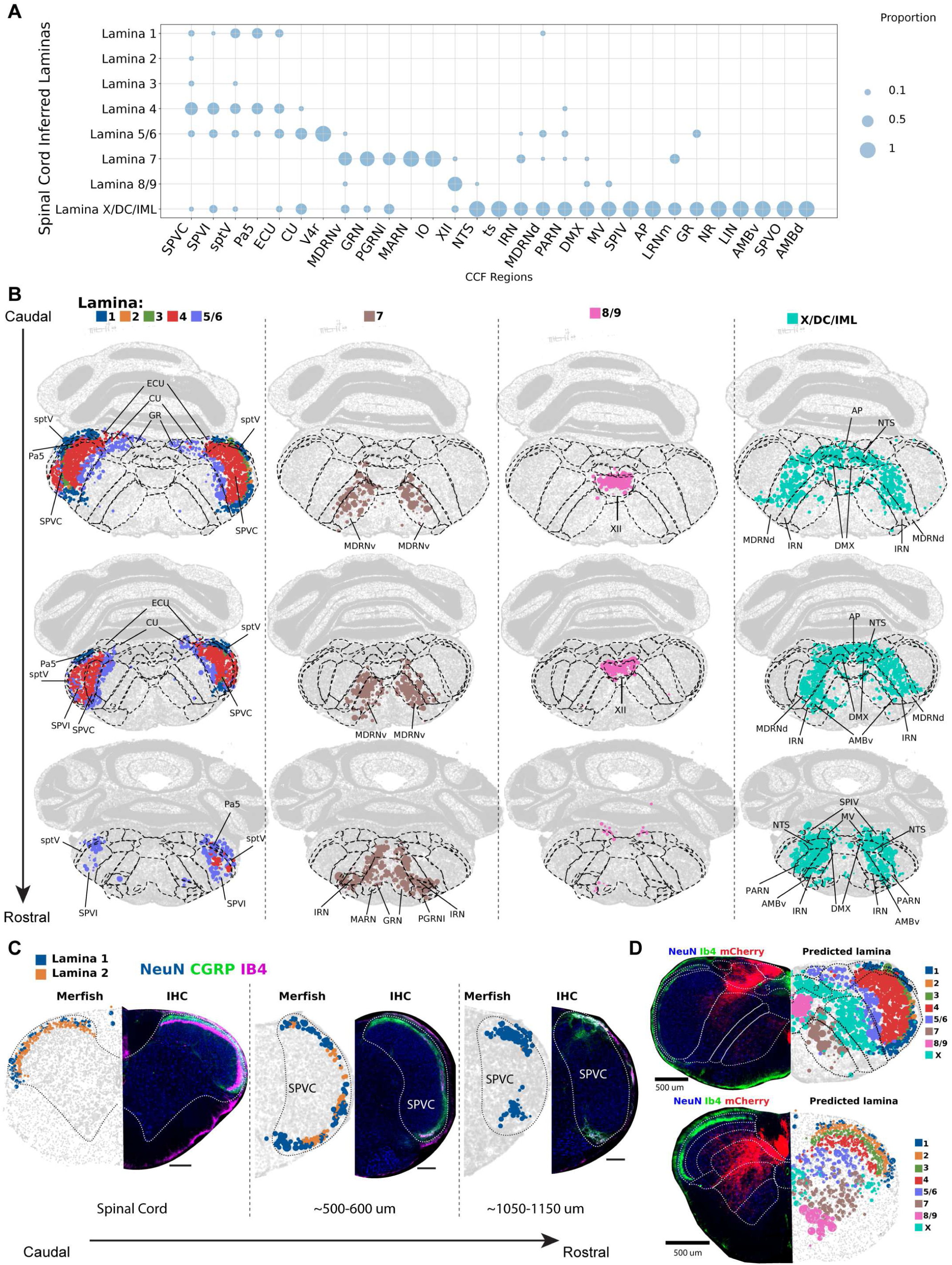
Correspondence between spinal cord-inferred laminae and anatomical regions in brainstem. **(A)** Bubble plot showing the distribution of each of the anatomical regions from CCF-defined anatomical regions between spinal cord-inferred laminae. The data is normalized per column, which means that the bubble size corresponds to the proportion of cells annotated from each of the CCF regions mapped to the given interfered lamina. **(B)** Representative sections of brainstem from caudal to rostral showing the most enriched CCF anatomical regions for each lamina. Each cell is represented by a dot with a size correlated to the cell area. **(C)** Co-registration of spatially assigned laminar identities with immunohistochemical markers across the rostral-caudal extent of the spinal cord and caudal brainstem. Each pair of panels shows MERFISH-assigned cells (left) colored by laminar identity (Lamina 1, blue; Lamina 2, orange) alongside the corresponding IHC image (right) stained for NeuN (blue), CGRP (green), and IB4 (magenta). Sections are shown from caudal spinal cord through rostral brainstem at the indicated rostral-caudal positions (∼500–600 µm, and ∼1050–1150 µm from the caudal reference). Scale bar is 250um. **(D)** Validation of laminar assignments across all Rexed laminae using corticofugal tract labeling and IB4. Each pair of panels shows MERFISH-assigned cells (right) colored by laminar identity (Laminae 1–9 and X as indicated in the legend) alongside the corresponding IHC image (left) stained for NeuN (blue), corticofugal projections (red), and IB4 (green).

**Supplementary Figure 12.**
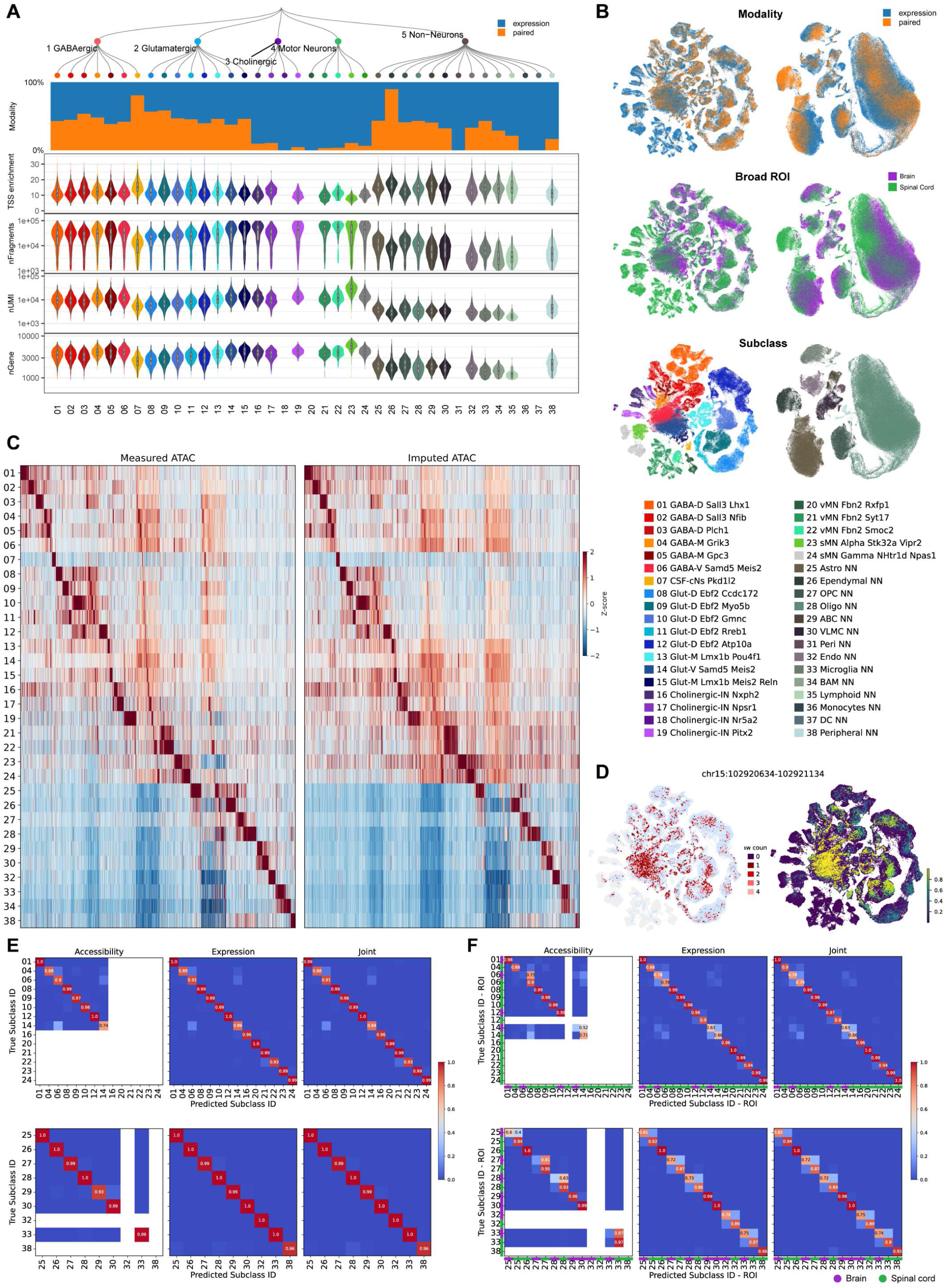
MultiVI integration of paired and unpaired single-cell chromatin accessibility and gene expression data across CNS cell types. **(A)** Quality control metrics for all cells included in the MultiVI model, organized by subclass and grouped by major cell class (dendrogram, top). The stacked bar plot shows the proportion of cells contributed by expression-only (blue) versus paired multiome (orange) modalities per subclass. Violin plots below show the distributions of TSS enrichment score, number of ATAC fragments (nFragments, log scale), number of UMIs (nUMI, log scale), and number of detected genes (nGene) per subclass for cells with paired modalities, colored by subclass identity. **(B)** UMAP embeddings of all neuronal (left) and non-neuronal (right) cells in the MultiVI joint latent space, colored by: sequencing modality (expression-only in blue, paired multiome in orange; top), broad region of origin (brain in purple, spinal cord in green; middle), and subclass identity (bottom). The separation and intermingling of cell populations across these annotations reflects the model’s ability to jointly embed cells across modalities and tissues. **(C)** Heatmaps comparing measured (and normalized) (left) versus MultiVI-imputed (right) chromatin accessibility across all 80,439 peaks (columns) for paired cells, grouped by subclass (rows, subclasses 01–38). Color indicates Z-scored accessibility (red = high, blue = low). Concordance between measured and imputed accessibility validates the MultiVI imputation across cell types. **(D)** Example locus-level view of chromatin accessibility at chr15:102,920,634–102,921,134 (a *Hoxc* locus peak). Left: UMAP with cells colored by measured accessibility raw count at this peak (white to dark red). Right: UMAP with cells colored by MultiVI-imputed accessibility (purple to yellow scale, 0–1.0), illustrating accurate imputation of region-specific chromatin accessibility. **(E)** Confusion matrix showing subclass classification accuracy using accessibility-only, expression-only, and joint (multimodal) MultiVI embeddings, for neuronal subclasses (01–24, top row) and non-neuronal subclasses (25–38, bottom row). Each cell shows the row-normalized proportion of true instances of a given subclass (y-axis) predicted as each subclass (x-axis), such that each row sums to 1. Diagonal values represent classification accuracy per-subclass. Numeric annotations are shown only when accuracy > 0.40. The joint embedding consistently achieves higher on-diagonal scores, demonstrating that multimodal integration improves cell type classification. **(F)** As in (E), but classification is performed at the level of subclass–region combinations (Subclass ID × ROI), distinguishing brain versus spinal cord identity within each subclass. The joint embedding resolves region-specific subclass identities that are ambiguous from either modality alone.

**Supplementary Figure 13.**
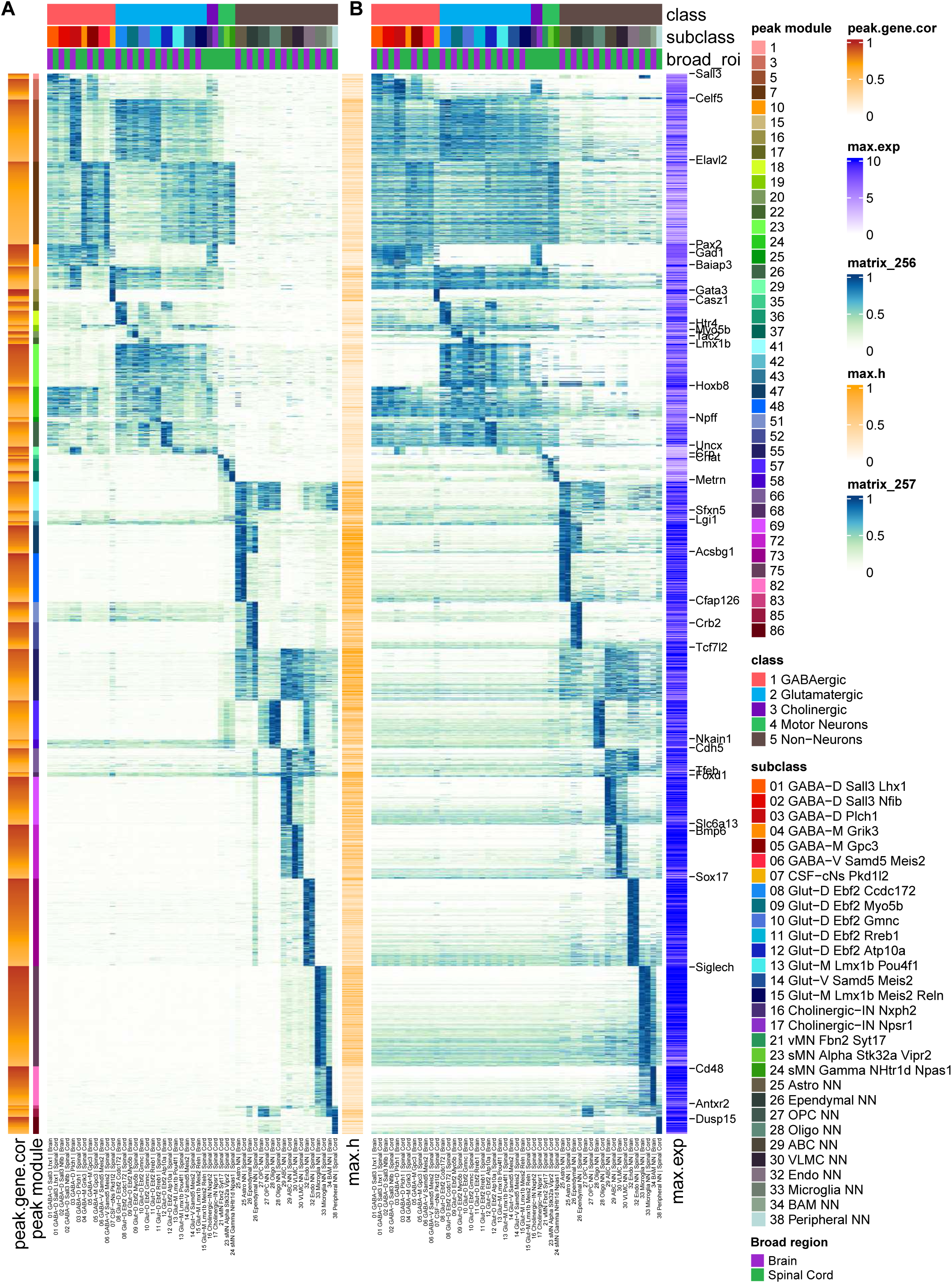
Chromatin accessibility modules and gene expression across cell types. **(A)** Heatmap of chromatin accessibility peaks (rows) across subclasses by region (columns), grouped by peak module. Rows are ordered by peak module identity (indicated by the color bar on the left, modules 1–86) and peak-to-gene correlation score (peak.gene.cor, orange-to-white scale, 0–1). Columns are annotated at the top by cell class, subclass, and broad region of origin (brain or spinal cord). Color intensity within the heatmap reflects the maximum accessibility score (max.h, orange scale, 0–1) per peak per cluster. **(B)** Paired heatmap displaying gene expression levels (blue scales, 0–1) for genes correlated with the chromatin accessibility peaks shown in (A). Rows correspond to the same peak modules as in (A), with select marker genes annotated on the right. Maximum gene expression values (max.exp, purple scale, 0–10) are indicated at the right. Together, panels (A) and (B) link cis-regulatory chromatin modules to their associated gene expression programs across transcriptomically defined CNS cell populations, spanning both brain and spinal cord.

**Supplementary Figure 14.**
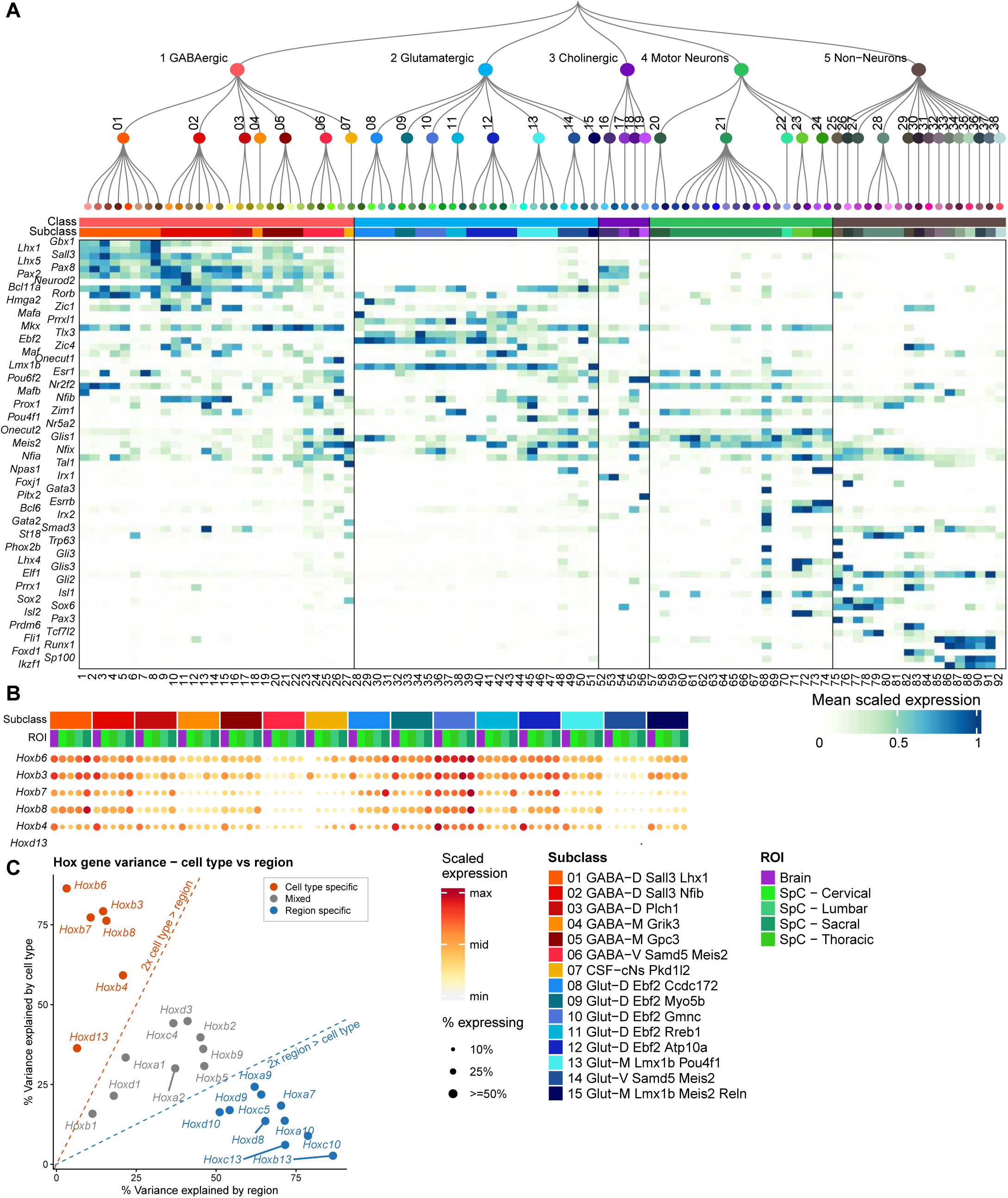
Transcription factor expression and cell type specific Hox gene expression across supertypes. **(A)** Heatmap of mean scaled expression for curated transcription factors (rows) across supertypes (columns). Columns are organized by a hierarchical dendrogram. Transcription factors shown include key class- and subclass-specific regulators such as *Lhx1*, *Lhx5*, *Pax6*, *Neurod2*, *Rorb*, *Tlx3*, *Ebf2*, *Lmx1b*, *Nfib*, *Nr5a2*, *Pitx2*, *Gata3*, *Sox2*, *Pax3*, *Tcf7l2*, *Runx1*, *Foxd1*, and *Ikzf1*, among others. **(B)** Dot plot of *Hox* gene expression that are cell type specific (*Hoxb6*, *Hoxb3*, *Hoxb7*, *Hoxb8*, *Hoxb4*, *Hoxd13*) across cell type subclasses (columns), annotated by subclass identity and region of origin (ROI: Brain, spinal cord cervical, lumbar, sacral, and thoracic). Dot size indicates the percentage of cells expressing the gene and dot color reflects scaled expression level. **(C)** Scatter plot quantifying the relative contribution of cell type identity versus anatomical region to *Hox* gene expression variance. Each point represents a single *Hox* gene, with the x-axis showing the percentage of variance explained by region and the y-axis the percentage explained by cell type. Points are colored by classification as cell-type-specific (orange), region-specific (blue), or mixed (grey). Dashed diagonal lines indicate 2-fold thresholds distinguishing cell-type-dominant from region-dominant variance. *Hox* genes with predominantly cell-type-driven expression (e.g., *Hoxb6*, *Hoxb3*, *Hoxb8*) are labeled in orange, while region-driven genes (e.g., *Hoxc13*, *Hoxb13*, *Hoxc10*) are labeled in blue.

**Supplementary Figure 15.**
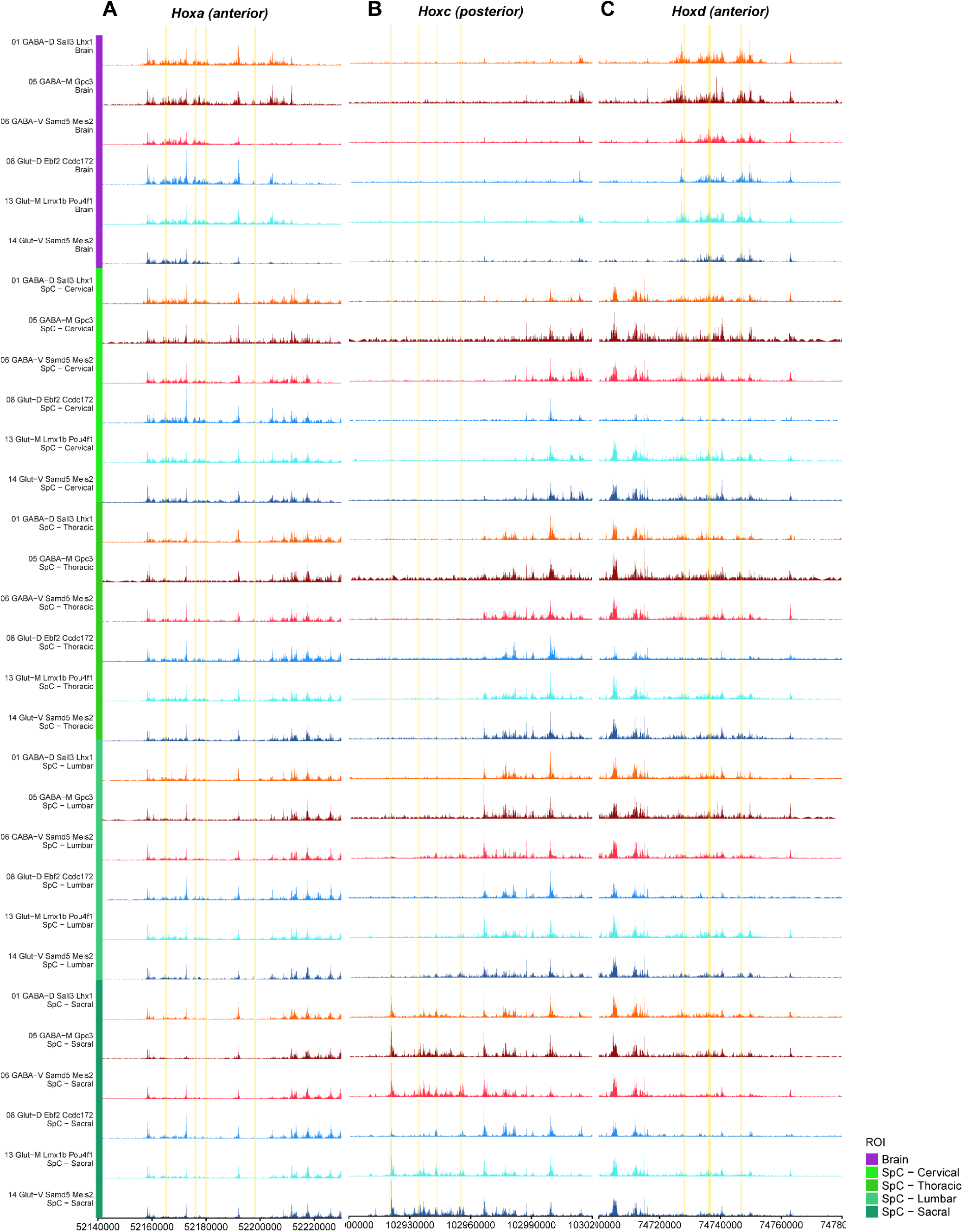
Chromatin accessibility at *Hox* gene loci across CNS cell types and spinal cord regions. (A-C) Genome browser tracks displaying chromatin accessibility (ATAC-seq) signal at three *Hox* gene cluster loci: (A) the *Hoxa* cluster (rostral; chr6: ∼52,140,000–52,230,000), (B) the *Hoxc* cluster (caudal; chr15: ∼102,920,000–103,020,000), and (C) the *Hoxd* cluster (rostral; chr2: ∼74,720,000–74,780,000). Each row represents a distinct combination of subclass and anatomical region of origin, grouped sequentially by region: Brain, spinal cord cervical (SpC – Cervical), spinal cord thoracic (SpC – Thoracic), spinal cord lumbar (SpC – Lumbar), and spinal cord sacral (SpC – Sacral). Within each region, six cell type subclasses are shown: GABA-D Sall3 Lhx1 (orange), GABA-M Gpc3 (dark red), GABA-V Samd5 Meis2 (pink/red), Glut-D Ebf2 Ccdc172 (blue), Glut-M Lmx1b Pou4f1 (cyan), and Glut-V Samd5 Meis2 (dark blue). Track color corresponds to cell type subclass identity. Vertical yellow dashed lines highlight specific regulatory peaks of interest shared across cell types and regions. Systematic differences in accessibility across spinal cord levels reflect the regional *Hox* expression codes that define rostrocaudal positional identity in the CNS.

**Supplementary Figure 16.**
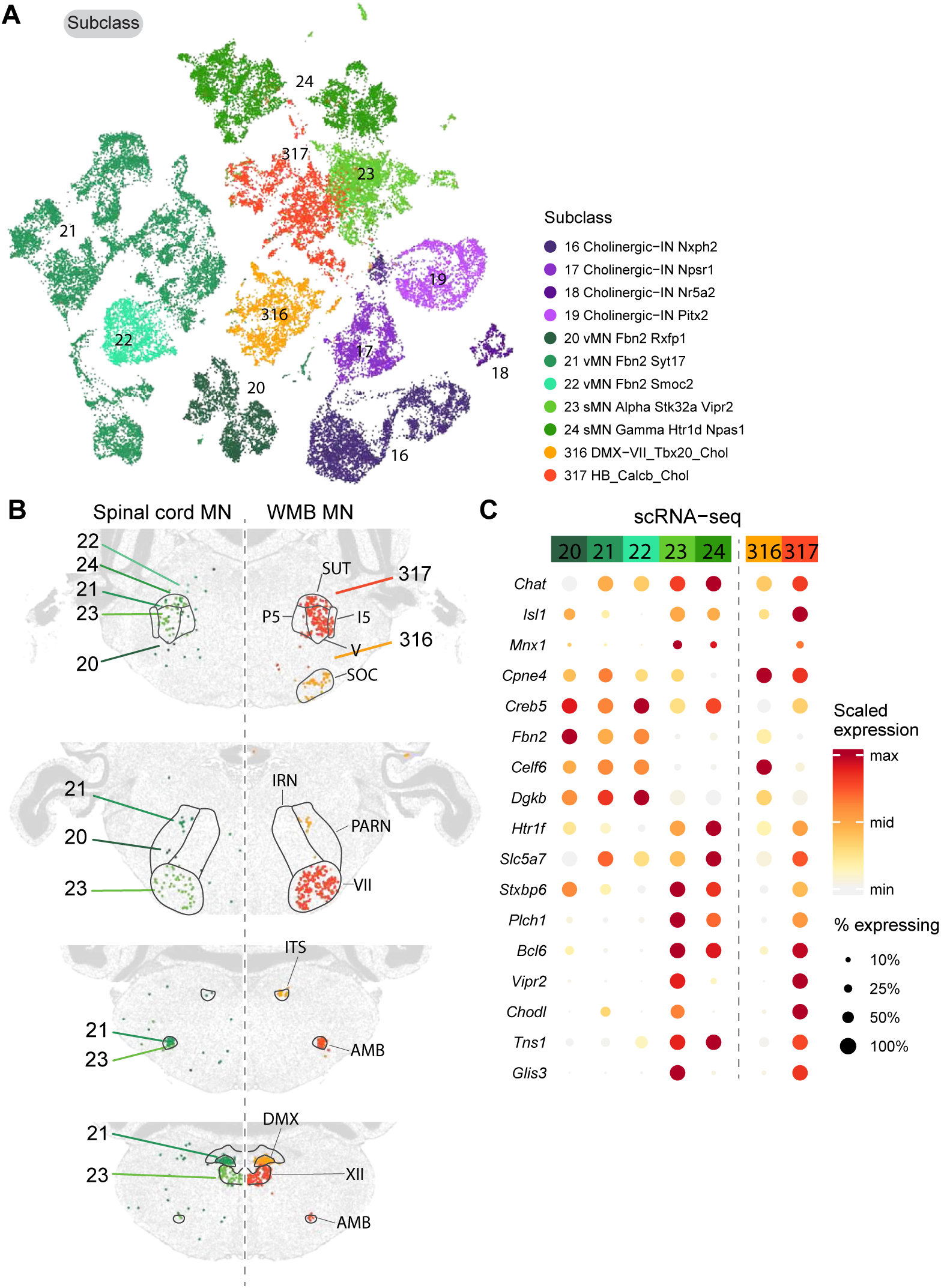
Brainstem motor neuron mapping validation. **(A)** Joint UMAP embedding of motor neuron and cholinergic interneuron populations from the spinal cord reference atlas and the Yao et al. whole-brain atlas (subclasses 316 and 317). Cells are colored by subclass identity as indicated in the legend. **(B)** MERFISH spatial sections spanning the rostral-caudal extent of the medulla, showing cells assigned to motor neuron and cholinergic interneuron subclasses. Colored dots indicate assigned subclass identity; grey cells are all other cell types shown as background. Black outlines indicate the boundaries of anatomically annotated cranial motor nuclei: hypoglossal nucleus (XII), dorsal motor nucleus of the vagus (DMX), nucleus ambiguus (AMB), facial motor nucleus (VII), trigeminal motor nucleus (V, P5-I5), superior olivary complex (SOC), inferior reticular nucleus (IRN), paragigantocellular reticular nucleus (PARN), and subthalamic nucleus (SUT). **(C)** Dot plot showing the expression of motor neuron marker genes across spinal cord motor neuron subclasses in the scRNA-seq reference atlas. Columns represent subclasses as indicated, grouped by spinal cord taxonomy (left, subclasses 20–24) and whole-brain atlas subclasses present in the dataset (right, subclasses 316–317). Dot color indicates mean log-normalized expression. Dot size indicates the proportion of cells within each subclass expressing the gene.

**Supplementary Figure 17.**
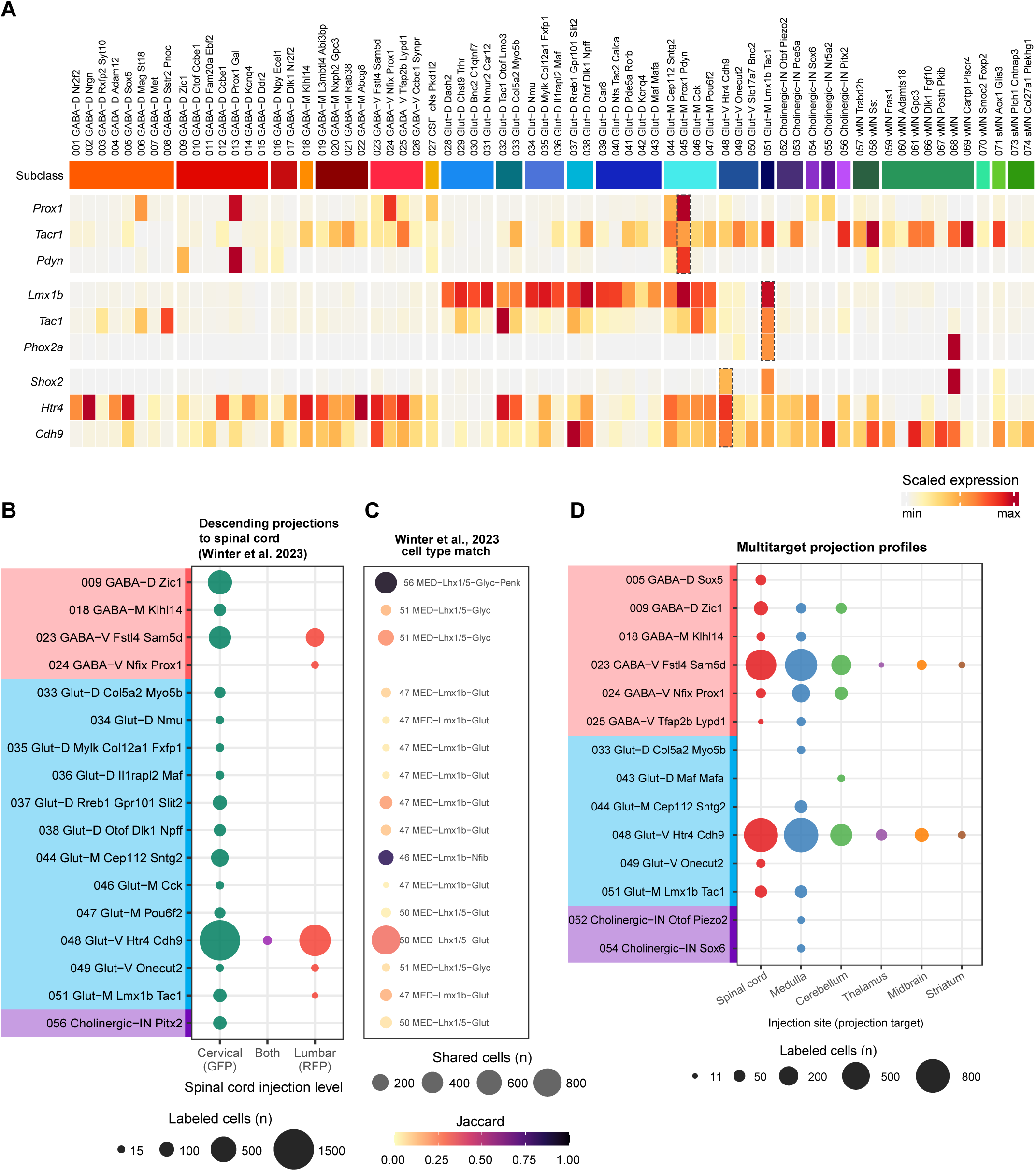
Retrograde tracing datasets link molecularly defined supertypes to projection identities. **(A)** Heatmap of scaled expression for ten projection-neuron marker genes (*Calca*, *Tacr1*, *Nts*, *Tac2*, *Lmx1b*, *Tac1*, *Phox2a*, *Shox2*, *Htr4*, *Cdh9*) across all neuronal supertypes (001–074), grouped by subclass. \**Phox2a* is expressed in a subgroup of 051 Glut-M Lmx1b Tac1. **(B)** Dotplot showing brain neurons retrogradely labeled from cervical (GFP, green) and lumbar (RFP, red) spinal cord injections (Winter et al. 2023^15^), assigned to neuronal supertypes in the shared latent space. Dot size indicates number of labeled cells from a spinal cord injection. Color strip on the left indicates neuronal class identity as in Figure 1. **(C)** Dotplot showing Jaccard similarity between each supertype and its best-matching Winter et al. 2023 cluster. Dot size indicates number of labeled cells; dot color indicates Jaccard similarity. **(D)** Dotplot showing medullary neurons retrogradely labeled from injections at multiple target sites, assigned to supertypes in the shared latent space.

